# Anatomy of a mega-radiation: Biogeography and niche evolution in *Astragalus*

**DOI:** 10.1101/2023.06.27.546767

**Authors:** R.A. Folk, J.L.M. Charboneau, M. Belitz, T. Singh, H.R. Kates, D.E. Soltis, P.S. Soltis, R.P. Guralnick, C.M. Siniscalchi

## Abstract

*Astragalus* (Fabaceae), with more than 3,000 species, represents a successful radiation of morphologically highly similar species found across the Northern Hemisphere. It has attracted attention from systematists and biogeographers, who have asked what factors might be behind the extraordinary diversity of this important arid-adapted clade and what sets it apart from close relatives with far less species richness. Here, for the first time using extensive taxonomic sampling in a phylogenetic analysis, we ask whether (1) *Astragalus* is uniquely characterized by bursts of radiation or is instead similar to related taxa. Then we test whether the species diversity of *Astragalus* is attributable specifically to its predilection for (2) cold and arid habitats or (3) particular soils. Finally, we test (4) whether *Astragalus* originated in central Asia as proposed and (5) whether niche evolutionary shifts were associated with the colonization of other continents. Our results point to the importance of heterogeneity in the diversification of *Astragalus*, with upshifts associated with the earliest divergences but attributable to no abiotic factor or biogeographic regionalization tested here. The only potential correlate with diversification we identified was chromosome number. We find strong evidence for a central Asian origin and direct dispersals from this region responsible for much of the present-day distribution, highlighting the importance of central Asia as a biogeographic gateway. In contrast to diversification shifts, biogeographic shifts have a strong association with the abiotic environment. Our most important result was a fundamental divide in soil types and diurnal temperature variation between the Eastern and Western Hemisphere species; this divergence does not reflect differences in available habitat among these biogeographic domains but may reflect unique local gains of edaphic and abiotic stress adaptations. While large clades are logistically difficult to tackle, our investigation shows the importance of phylogenetic and evolutionary studies of “mega-radiations.” Our findings reject any simple key innovation behind the dominance and richness of *Astragalus* and underline the often nuanced, multifactorial processes leading to species-rich clades.

## INTRODUCTION

*Astragalus* (milk-vetch or locoweed; Fabaceae: Papilionoideae; Azani et al., 2017a), with ∼3,100 species (https://powo.science.kew.org/; https://astragalusofworld.com/), is currently the largest clade in land plants treated at generic rank. While mega-genera arise primarily from taxonomic history (Frodin, 2004) and are not directly comparable to less species-rich genera (Hennig, 1966; Sanderson and Wojciechowski, 1996), they are clearly related to morphological evolution because they often arise from a paucity of easily recognizable morphological variation that would delimit more manageable units, ultimately adding to the logistical challenge of their study. Aside from species number, the *Astragalus* clade is remarkable as a major nodulating legume radiation in temperate arid areas, particularly central Asia and western North America where it is one of the most ecologically important legumes. *Astragalus* species, relatively uniform in terms of morphology (primarily herbaceous perennials) and niche biology (primarily arid and semi-arid areas), are morphologically highly similar to other members of the Astragalean clade (e.g., *Oxytropis*, *Colutea*), offering no clear adaptive explanation as to their extraordinary diversity and ecological importance.

The ecological dominance, species richness, and narrow endemism of *Astragalus* in many semi-arid areas of the Northern Hemisphere have attracted the attention of biogeographers and ecologists (Scherson et al., 2008; Samad et al., 2014; Hardion et al., 2016; Amini et al., 2019; Azani et al., 2019; Záveská et al., 2019; Plenk et al., 2020; Li et al., 2021; Wang et al., 2021; Maassoumi and Ashouri, 2022). Perhaps the major biogeographic finding of molecular phylogenetic studies in *Astragalus*, suspected previously on the basis of chromosome counts (Ledingham, 1960; Barneby, 1964), is the recovery of a clade comprising nearly all North American *Astragalus*, informally termed the Neo-Astragalus clade (Wojciechowski et al., 1993, 1999), which renders species in the Eastern Hemisphere a paraphyletic group. The Neo-Astragalus clade is distinguished by chromosomal processes. Polyploidy, probably often allopolyploidy (Bartha et al., 2013; Záveská et al., 2019; Plenk et al., 2020), is thought to be the dominant process in Eastern Hemisphere species (Lewis et al., 2005), where counts of *n =* 8, 16, interpreted as euploid, predominate. By contrast, aneuploidy prevails in the Neo-Astragalus clade (Wojciechowski et al., 1993, 1999), with the aneuploid series *n =* 11–15 (Spellenberg, 1976) forming a “nearly perfect correlation” with biogeography, according to Wojciechowski et al. (1999).

Rapid chromosomal evolution, in the form of long aneuploid series seen in some plant groups, has been associated with responses to environmental stress (De Storme and Mason, 2014; Ackerfield et al., 2020) and has been interpreted as promoting recombination in the face of ecological change more generally (Levin, 1975; Grant, 1981). While Neo-Astragalus is not clearly morphologically delimited (Wojciechowski et al., 1993), the strong biogeographic and chromosomal pattern is in line with niche biology. Barneby (1964) observed that, by contrast with the dominance of Eurasian *Astragalus* in the highlands, North American species that he believed had developed in situ seem to specialize in challenging dry lowland environments. The genus is also associated with poor soils, for which its nodulating ability (Afkhami et al., 2018; Kates et al., 2022) would serve as a key adaptation. Edaphic specialization is thought to be a major force in plant biogeography (Raven, 1964; Axelrod, 1972; Lichter-Marck and Baldwin, 2023), and while large-scale investigations of edaphic ecology remain few in nodulating legumes (Tamme et al., 2021; Doby et al., 2022; Siniscalchi et al., 2022), soil environments control the distribution of root nodule symbiosis by limiting the availability of both plant and especially bacterial partners and affecting partner choice (Sprent et al., 2017; Dludlu et al., 2018; Rathi et al., 2018; Ardley and Sprent, 2021). Differing availability of symbiotic partners at the regional scale is thought to be a major factor overall limiting the dispersal process in nodulating plants (Parker et al., 2006; Simonsen et al., 2017). Therefore, it appears that a continental shift to the Americas, likely before the radiation of today’s major clades, was associated with ecological specialization to challenging low-nutrient soil and harsh climatic conditions, along with more rapid chromosomal evolution.

Finally, related to questions of biogeography and chromosomal change are diversification mechanisms in *Astragalus*. Given its high species richness, the traditional explanation was that *Astragalus* underwent particularly high diversification rates (Barneby, 1964; Polhill, 1981; Lewis et al., 2005), and the genus was among the earliest in plants where explicit rates of diversification were calculated from molecular phylogenetic data (Sanderson and Wojciechowski, 1996). Remarkably, the primary finding of Sanderson and Wojciechowski was that *Astragalus* is not characterized by extraordinary rates of diversification, but instead shares uniform, relatively high rates of diversification with other members of the broader “Astragalean” clade (tribe Galageae pro parte; Lewis et al., 2005), a group distinctive in its ecology but lacking in any clear morphological innovations that might relate to diversification. Use of more recent probabilistic methods to estimate diversification in *Astragalus* have focused on recently diverged subclades (Azani et al., 2019, using BAMM) or on crown dates of diversification events using time calibration (Hardion et al., 2016; Bagheri et al., 2017). A reevaluation of the uniform diversification hypothesis of Sanderson and Wojciechowski (1996) is warranted with more complete sampling and robust phylogenetic trees enabling the investigation of diversification rates both within and between genera, as well as diversification methods that more fully use the branch length information from molecular phylogenetics beyond species count data.

Both intrinsic and extrinsic factors that shape diversity in *Astragalus* have been posited, but empirical work has been piecemeal. Areas of species richness and endemism have recently been reported for Eurasian *Astragalus* (Maassoumi and Ashouri, 2022), and *Astragalus* has also been the subject of study in terms of biogeography and diversification, particularly in specific subclades (Sanderson and Wojciechowski, 1996; Scherson et al., 2008; Azani et al., 2019; Su et al., 2021). A needed next step is to test, across the entire clade, the associations between biogeographic processes such as lineage dispersal, chromosome evolution, and shifts in niche occupancy. Here, we use a combination of comparative methods and spatial phylogenetics to dissect the drivers of diversity in *Astragalus.* We test five main hypotheses: we first test whether (1) the high species diversity of *Astragalus* is due to uniform, relatively high speciation (cf. Sanderson and Wojciechowski, 1996) and not to bursts of diversification within the genus. We then test whether (2) *Astragalus* diversification is primarily associated with cold and arid habitats or with (3) shifts in soil type. Finally, we test whether (4) shifts in habitat are associated with shifts between biogeographic regions; and (5) whether *Astragalus* originated in West Asia with direct connections to other realms from this ancestral region.

## METHODS

### Taxon sampling

The taxa sampled are part of the NitFix sequencing initiative ((https://nitfix.org/; Kates et al., 2022), with molecular data reanalyzed for the purpose of this study. Specimens were sampled from herbarium specimens using a sample management protocol reported previously (Folk et al., 2021a). In total, 939 taxa were included, with 847 in the ingroup. Outgroup sampling was extensive across the Astragalean clade sensu Lewis et al. (2005) in order to yield a strong biogeographic foundation. Representatives were included of *Clianthus, Colutea, Eremosparton, Erophaca, Lessertia, Smirnowa, Sphaerophysa, Sutherlandia, Swainsona,* and the *Astragalus* segregates *Biserrula*, *Podlechiella,* and *Phyllolobium. Oxytropis*, biogeographically similar to *Astragalus*, was extensively sampled (71 taxa). Samples of the segregate genera *Ophiocarpus* and *Barnebyella* (Lewis et al., 2005) were not available, but these are almost certainly better treated as *Astragalus* (Kazempour Osaloo et al., 2003). As the most distant outgroup, *Wisteria* (sensu Lewis et al. 2005, a member of the inverted repeat-lacking clade [IRLC] but outside the Astragalean clade; Wojciechowski et al., 1999, 2004) was used to root the trees. Supplementary Table S1 provides accession information; assignments to clade and sections given the phylogenetic results (see below) are available in Supplementary Figure S1. Sampling statistics for the phylogeny are reported in Results.

### Molecular methods

DNA extracts were prepared from herbarium specimens according to a modified CTAB protocol (Doyle and Doyle, 1987) reported previously (Folk et al., 2021a) and performed at the University of Florida (Gainesville, FL, USA). Specimens were then cleaned of remaining secondary compounds with SPRI beads and submitted for preparation of Illumina TruSeq libraries, omitting a size selection step to account for DNA degradation. We used a Hyb-Seq approach (Weitemier et al., 2014) targeting 100 low-copy nuclear loci, with data generated as part of the NitFix sequencing initiative (Kates et al., 2022). Library construction and target capture were performed by Rapid Genomics (Gainesville, FL, USA). Details concerning the baitset have been reported previously (Folk et al., 2021a; Kates et al., 2022).

### Phylogenomic assembly

Sequences were assembled using the HybPiper v1.3 pipeline (Johnson et al., 2016). Given the wide phylogenetic scope of the probe set used for hybridization, spanning the four taxonomic orders that comprise the nitrogen-fixing clade of angiosperms, target recovery was achieved with the use of an amino acid reference for each locus, instead of DNA, with the BLAST option for read mapping. HybPiper flags loci for which multiple contigs are assembled as potential paralogs on a per-species basis, deciding the putative “ortholog” by assessing sequencing depth and contig length. The hybpiper_stats.py script was used to extract assembly information; given the low percentage of flagged paralogs (96% of the samples had 5% or fewer of loci flagged as potentially paralogous; paralogy flags are summarized in Supplementary Fig. S6), we did not employ further paralog resolution methods, instead using the potential “ortholog” selected by the pipeline. Assembly success in terms of number of assembled loci is summarized in Supplementary Fig. S7. General assembly stats are available on the project GitHub (https://github.com/ryanafolk/astragalus_niche_biogeo/blob/main/tree_quality_control/qc_astragalus_rename.csv).

### Phylogenomic analysis

Individual gene trees were obtained using maximum likelihood in RAxML v.8.2.11 (Stamatakis, 2014), with 100 bootstraps and using the GTR+G nucleotide substitution model. A species tree was obtained with the multispecies coalescent method in ASTRAL-III (Zhang et al., 2018), with support values calculated as local posterior probabilities (LPP). As coalescent species trees do not have terminal branch lengths without population sampling and measure branch length partly in terms of population size, branch lengths were re-optimized in RAxML using the option “-f e” to yield branch lengths in expected per-site substitutions.

The penalized likelihood method implemented in TreePL (Smith and O’Meara, 2012) was used to date the tree. Secondary calibration points were extracted from Azani et al.’s (2019) chronogram, consisting of three calibrations: a root constraint (min: 32.33 mya, max: 44.82 mya), *Oxytropis* crown group (min: 1.3 mya, max: 6.3 mya), and *Oxytropis/Astragalus* split (min: 11.93 mya, max: 20.33 mya). The “prime” option was used to calculate the initial parameters, and the analysis was conducted with cross-validation and a smoothing rate of 100.

### Assembly of locality data

We assembled a dataset of georeferenced occurrence records for species of *Astragalus* by first downloading all occurrence records in Fabales from the biodiversity discovery platforms iDigBio (accessed May 21, 2020) and the Global Biodiversity Information Facility (GBIF, 2020). Using a list of all recognized Fabales species names and their associated synonyms, we aggregated all records of synonyms to accepted species names. This Fabales dataset was then filtered to include only species in *Astragalus* and *Oxytropis* and further sampled outgroups. Next, occurrence records were subjected to a data cleaning process that relied heavily on the R package CoordinateCleaner (Zizka et al., 2019). In this process, records were removed if they were missing coordinates, or if latitude values were not between -90 and 90 or if longitude values were not between -180 and 180. We also removed records if coordinates had equal longitude and latitude values, were within 500 m of the geographic centroid of political countries or provinces, were within 500 m of the GBIF headquarters, had either zero longitude or latitude, or were within 100 m of a global database of biodiversity institutions that includes zoos, botanical gardens, herbaria, universities, and museums (Zizka et al., 2019). Records were also removed if the minimum absolute distance between a record and all other records of the same species was greater than 1000 km. Duplicate records with the same longitude and latitude values were filtered to only retain one record.

### Assembly of niche attributes

We assembled a set of niche attributes selected to best capture properties of interest for *Astragalus,* such as climate and soil, while selecting parameters that show minimal correlation at global scales (Pearson’s *R*^2^ < 0.7). Using previously published code (Folk et al., 2019), we extracted niche attributes for 18 predictors representing climate, soil, elevation, and landcover. Bioclimatic data (Hijmans et al., 2005) were BIO1, BIO2, BIO3, BIO4, BIO7, BIO12, BIO15, BIO17. Variable choice from the larger set of 19 Bioclim variables was guided by mean annual temperature (BIO1) and annual precipitation (BIO12), as well as variability in temperature (BIO2–4, BIO7) and precipitation (BIO15 and BIO17). Elevational data were derived from GTOPO30 (https://doi.org/10.5066/F7DF6PQS). Soil data derived from SoilGrids (Hengl et al., 2017) were nitrogen and carbon content, sand and coarse fragment percent, and the most likely of the 29 WRB 2006 (World Reference Base) soil classifications (see https://www.fao.org/soils-portal/data-hub/soil-classification/world-reference-base/en/) for each grid cell. Finally, landcover data (Tuanmu and Jetz, 2014) comprised herbaceous percent land cover (i.e., grassland) and coniferous (“needle-leaf”) and broad-leaf forest percent land cover. For the quantitative variables (all attributes except for soil type classification), species’ niche occupancy was summarized by the mean of all extracted occurrence values. For soil type, a categorical variable, the mode was used. Ancestral reconstructions of environmental data were implemented in the R package phytools, with model choice via AIC_c_; the model set included Brownian motion, Ornstein-Uhlenbeck [OU], and early-burst models for quantitative variables; OU models were selected without exception. For categorical soil type only the equal-rates model could be implemented due to the high number of states.

### Biogeography

Coding of geographic regions followed a previously developed shape file and Python code for regionalization (Folk et al., 2021b). Briefly, considering the regions most important to *Astragalus,* North and South America were recognized as distinct regions, as was Africa, including the southern Mediterranean. Western Europe and the northern Mediterranean were considered a region combined with boreal Eurasia, distinct from West Asia (the most important area for *Astragalus* in terms of species richness), the latter delimited from Anatolia east to the Altai, north to the Pontic area and south to the Arabian Peninsula following arguments in (Folk et al., 2021b) (Fig. 1). East Asia and South Asia were recognized together as a further distinct region. In total, seven regions were recognized as summarized in Fig. 1. Biogeographic scoring was manually checked with Plants of the World Online (http://powo.science.kew.org/) and information from the *Astragalus of the World* website (https://astragalusofworld.com/). We used a model comparison framework in BioGeoBEARS (Matzke, 2013) to select biogeographic models and favored models without jump parameters (DIVA-like, BAYAREA-like, and DEC models) following previous arguments (Ree and Sanmartín, 2018). The maximum range size was set to five, following the largest range size observed in extant taxa.

**Fig. 1.**
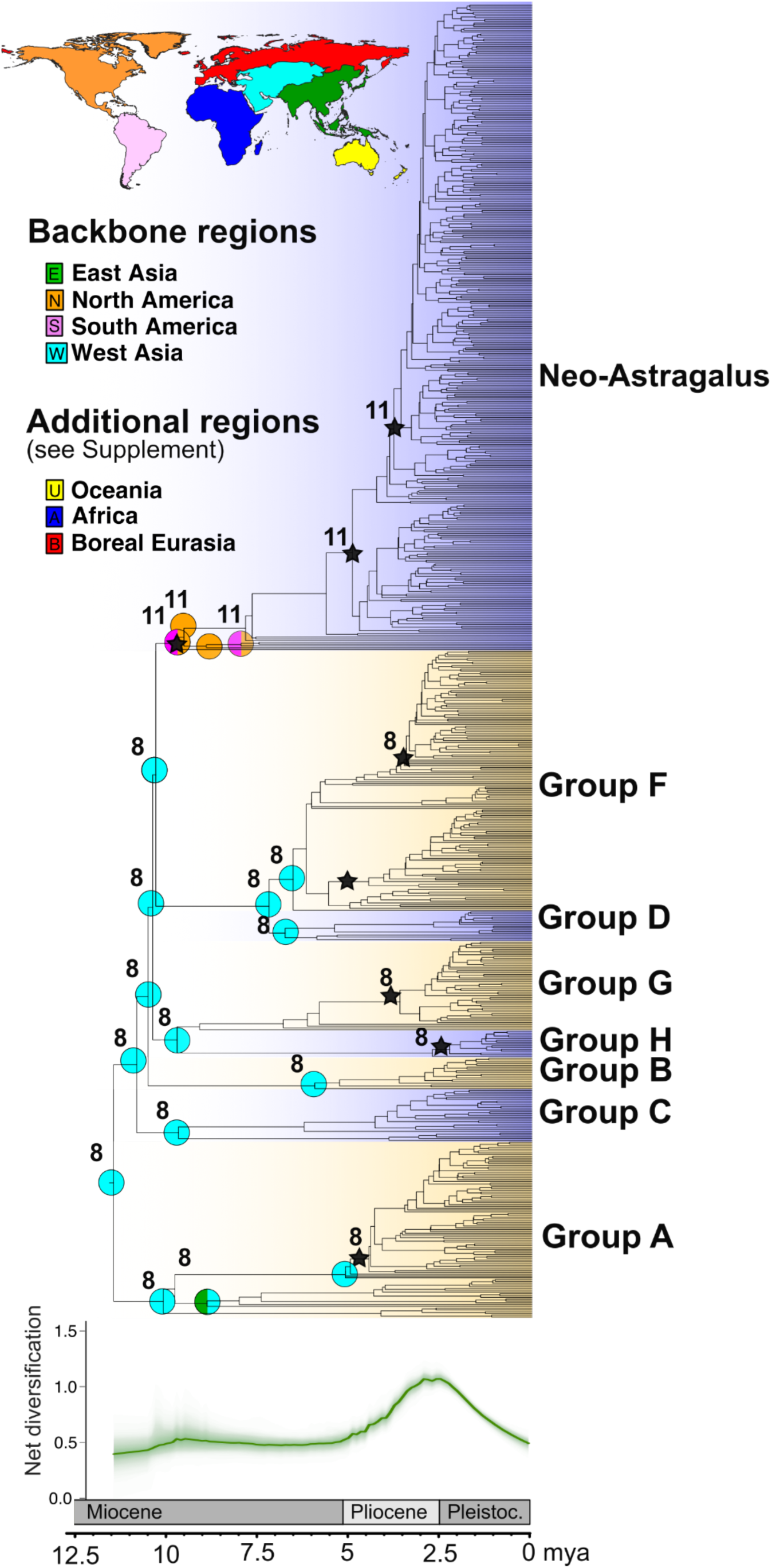
Biogeography and diversification of *Astragalus.* Ancestral biogeographic regions are indicated by colored circles according to the map legend; only *Astragalus* s.s. is shown here, and only backbone nodes and crown nodes of named clades are plotted. See Supplementary Fig. S5 for full result. Black stars denote all recovered BAMM diversification shifts; see Supplementary Fig. S2 for branches colored by diversification rate. Boldface numbers indicate reconstructed ancestral chromosome numbers, plotted for all nodes with diversification shift and/or biogeographic region plotted; see Supplementary Fig. S4 for full result.

### Examining polyploidy

Here we used both reported chromosome counts and ploidy estimates derived directly from sequence data to investigate chromosome evolution in *Astragalus.* Both represent alternative views of polyploidy, and determinations based on genomic data represent new techniques with substantial recent interest (Viruel et al. 2019; Viruel et al. 2023). We gathered chromosome data from CCDB ((Rice et al., 2015); http://ccdb.tau.ac.il/) and the *Astragalus* taxonomic database (https://astragalusofworld.com/), yielding 418 taxa with data. For ploidal determinations from sequence data, we used nQuire (Weiß et al., 2018; Viruel et al., 2019). nQuire uses heterozygous site count data to distinguish biallelic, triallelic, and tetraällelic inheritance patterns. Considering only sites with two bases observed, the distribution of observed base ratios in read pileups (e.g., 1:1, 1:2, 1:3, etc.) are plotted in a histogram and compared with expected frequencies under diploid, triploid, and tetraploid models using a model choice framework. While this method shows promise for analyzing sequence data, three assumptions underlying the method should be noted: (1) nQuire only considers diploid, triploid, and tetraploid models and cannot directly test higher ploidal levels, (2) it cannot test for aneuploidy, and (3) it was not originally intended for the case of allopolyploidy. nQuire has been shown to recover ploidal levels fairly well compared to direct measurement as long as ploidal levels are low, with increasing difficulty recognizing high ploidal levels from pentaploidy upwards (Viruel et al., 2019), but this positive result likely depends on the genomic complexity of study species and sequencing experimental parameters, and therefore requires further research. The implications of the other assumptions are not well-understood. nQuire outputs several model choice statistics; here, calls were based on AIC. To obtain the read pileups, we used BWA (Li and Durbin, 2009) to map reads back to a reference comprising a 50% majority consensus of all taxa contained in the hybridization probe set. Sorted BAM files were prepared with SAMtools (Li et al., 2009) and used as input in the nQuire pipeline. Final ploidy calls were used for ancestral reconstruction in phytools; the model set included equal-rates and all-rates-different models with the latter selected.

### Chromosome number reconstruction

Chromosome ancestral states were calculated using the ChromEvol model (Glick and Mayrose, 2014) as implemented in RevBayes (Höhna et al., 2016), considering rates of chromosome gains, losses, and polyploidization. The calibrated tree was trimmed to match the species for which counts were available, using the “drop.tip” function in the R package ape (Paradis et al., 2004). The analysis was run for 50,000 generations, with a 25% burnin. Analysis results were summarized and plotted using the R package RevGadgets (Tribble et al., 2022).

### Diversification rates

Estimation of diversification rates has been controversial (Moore et al., 2016; Louca and Pennell, 2020), so we used a combination of approaches here including both parametric and semiparametric methods and estimates across timescales including the present-day (cf. Louca and Pennell, 2020). To obtain semiparametric estimates of speciation rate (cf. Title and Rabosky, 2019) across *Astragalus* species, we used tip speciation rates according to the DR statistic of (Jetz et al., 2012), calculated using the function of Sun et al. (2020). To estimate shifts in diversification rate regimes, we used BAMM v. 2.5.0 (Rabosky, 2014). Priors followed recommendations from the R package BAMMtools, with additional settings as follows: expected shifts = 1 per documentation, segLength (likelihood grain) = 0.02, minimum clade size for shifts = 2 (i.e., shifts constrained to internal branches). Incomplete sampling was accounted for using the taxonomic data in *Astragalus of World* (https://www.astragalusofworld.com/). Because clade membership of unsequenced taxa is not certain (see Results), we assigned different sampling probabilities for Neo-Astragalus (0.4472) and the Eurasian clades (0.2041), as this was the lowest phylogenetic level at which applying taxonomic species count data for previously unsequenced species was feasible. Because outgroup sampling practices violate sampling assumptions of birth-death models, diversification estimation was limited to the ingroup, *Astragalus s.s*. MCMC was run in four chains for 64.4 million generations (stopped based on ESS > 400 and strong convergence), δT = 0.01, swap period = 1,000 generations, and sampling every 100,000th generation; further parameters followed BAMM documentation.

Tests of trait-associated diversification were implemented both within the BAMM method and external to BAMM using tip speciation rates under a model-free paradigm. BAMM rate-trait association tests were implemented in STRAPP (Rabosky and Huang, 2016), which is a semi-parametric permutation test using assigned BAMM diversification regimes to test for trait-associated regime shifts. Semi-parameteric tip rate tests used ESSim (Harvey and Rabosky, 2018) for continuous data and FiSSE (Rabosky and Goldberg, 2017) for binary data. All three tests are designed to control for evolutionary pseudoreplication, which can result in spurious significance in SSE-type models (see Rabosky and Goldberg, 2015). Multistate data (ploidy, soil type) were recoded to binary characters to satisfy requirements for FiSSE as described in Results.

### Data availability

Scripts and data to produce the analyses are available on GitHub at https://github.com/ryanafolk/astragalus_niche_biogeo. Sequence data are deposited at SRA as noted in Appendix S1 (BioProject XXX). Trees have also been deposited on TreeBase at XXX.

## RESULTS

### Sampling

We included 939 accessions, including 92 outgroups in the final phylogenetic tree. Of these, 373 taxa were assignable to Neo-Astragalus (44.7% of the total described diversity based on accepted taxa in https://astragalusofworld.com/) and 474 taxa assignable to the Eurasian (“Old World”) clades (20.4% of the total). While the Eurasian species are therefore undersampled, species included here represent all of the recognized major clades (Groups A to F; Kazempour Osaloo et al., 2003, 2005; Azani et al., 2017b; Su et al., 2021). Of the names matched to recognized groups in the *Astragalus of World* taxonomic database (https://astragalusofworld.com/), this sampling effort comprises 16.3% of species in Group A, 15.1% of Group B, 50% of Group C, 16.5% of Group D, 12.6% of Group H, 7.6% of Group G, and 14.2% of Group F. On the basis of sequencing quality and phylogenetic placement, 21 taxa in the phylogeny were excluded from downstream analysis; exclusions mostly pertained to the Neo-Astragalus clade, and these are enumerated in the GitHub project repository (https://github.com/ryanafolk/astragalus_niche_biogeo). For downstream analyses using the phylogeny (e.g., biogeographical and environmental analyses), we also randomly subsampled the tree terminals to one accession per species (thus eliminating multiple subspecific taxa and other cases of multiple samples per species), resulting in a tree with 818 tips.

### Phylogenetic relationships

Relationships within the Astragalean clade differed from summary relationships reported by Wojciechowski et al. (2000), Lewis et al. (2005), and Azani et al. (2019), but agree with those of Su et al. (2021), with a strongly supported monophyletic *Astragalus* s.s. (LPP [Local Posterior Probability] 0.99) sister to the remainder of the Astragalean clade, comprising *Oxytropis* + Coluteoid clade sensu Su et al. (2021). These relationships were primarily well-resolved, although the clade supporting *Biserrula epiglottis* as sister to the remainder of the Astragalean clade exclusive of *Astragalus s.s*. was only moderately supported (LPP 0.79).

Within *Astragalus,* we recovered an overall set of clades similar to those of (Kazempour Osaloo et al., 2005). The clade names in Fig. 1 follow the system of Groups A–H for Eurasian species in Kazempour Osaloo et al. (2003, 2005). The backbone was well-supported with all LPP > 0.96. As seen earlier, Group A is sister to the rest of the genus. Groups C and B, whose branching order was unresolved in earlier studies, are successively sister to the remainder of *Astragalus.* Group G, recovered as a paraphyletic grade of two clades named G1 and G2 in Kazempour Osaloo et al. (2005) and as a polytomy with Group G3 in Kazempour Osaloo et al. (2003), was recovered as monophyletic here, although with limited shared sampling among these studies. Group E was not recovered; two species sampled here (*A. sinicus*, *A. alpinus*), also included in Kazempour Osaloo et al. (2005), were recovered here in (respectively) a clade of Asian species sister to the remainder of Group C, and embedded in Group D. Group H was recovered as sister to Group G without support (LPP < 0.5). Clades corresponding to Groups D and F were recovered as sister (LPP 0.88), and this clade was sister to Neo-Astragalus (LPP 0.96). The monophyly of Groups A–D were not supported (LPP < 0.5; although confidently supported subclades were recovered), but the remainder were well-supported (LPP > 0.95).

Additional species that had not been previously sequenced have been tentatively assigned to these groups in an *Astragalus* taxonomic database (https://astragalusofworld.com/). Our results partly agree with these taxonomic assignments, with many exceptions; exhaustive taxonomic subsectional and clade information can be found in Supplementary Fig. S1. Group F contained the highest number of species previously assigned to other groups, especially of species assigned to Group B (Supplemental Fig. S1).

Our results were mostly similar to recent studies of Asian *Astragalus* (Azani et al., 2017b; Su et al., 2021), with our Group F corresponding to the Hypoglottis and Diholcos clades of these studies. The Contortuplicata and Hamosa clades correspond to the Asian clade recovered within our Group C, and the Phaca clade corresponds to our Group A. Our study finds Neo-Astragalus as an independent clade not placed within Diholcos (Group F), but Azani et al. (2017b) and Su et al. (2021) had limited sampling of Neo-Astragalus. The Glottis clade, sister to all other *Astragalus* in Azani et al. (2017b) and Su et al. (2021), was recovered here as closer to *Oxytropis* among the outgroups (represented by *Astragalus epiglottis* = *Biserrula epiglottis*). We also did not recover a Pseudosesbanella clade as sister to the remainder of *Astragalus* exclusive of the Glottis clade (Su et al., 2021); instead, the species we sampled of the Pseudosesbanella clade as previously recognized are early-diverging within Group A.

### Biogeographic results

The favored DEC model (ΔAIC_c_ 97 compared to second-best BAYAREA-LIKE model) recovered a strongly supported origin of *Astragalus* in West Asia ∼11.47 mya (crown ages hereafter), somewhat younger than, but consistent with, estimates from previous studies focusing on late Miocene origins. For example, (Azani et al., 2019) recovered a crown age of 14.16 mya (11.47–16.95 95% HPD) and also reconstructed West Asian origins; Su et al. (2021) recovered a crown age of 12.51 mya (9.45–16.00 95% HPD). Among named clades traditionally recognized in *Astragalus*, all named Eastern Hemisphere clades (Groups A–H) retained the ancestral West Asia distribution at the crown divergence.

Dispersals to the other two important regions for the Eastern Hemisphere grade of *Astragalus*, boreal Eurasia and Africa, occur almost solely through West Asia and primarily in the last 2–3 my. Likewise, despite a strong presence in eastern Asia, primarily in the Himalaya-Qinghai-Tibet area, *Astragalus* shows no examples of the Eastern Asia-Eastern North America floristic disjunction. Almost without exception, eastern Asian clades descend by vicariance from ancestors with Eastern and West Asian distributions; *Astragalus* clade A has a single example of a boreal-eastern Asia vicariant event.

Almost all *Astragalus* species of the Americas have long been known to form a clade (Wojciechowski et al., 1993), and our more strongly sampled results are consistent with this. In the DEC model, a single dispersal from West Asia led to a broad ancestral distribution across the Americas, reconstructed at 9.84 mya (older than the date of 4.4 mya recovered in Scherson et al. [2008] and 4.36 mya in Azani et al. [2019]), immediately followed by a vicariance across North and South America, with the North American lineage leading to most of the diversity of Neo-Astragalus. Most South American species represent at least 13 subsequent dispersals to South America from North America with dates between 2.5–4 million years, four of which represent clades of 2-17 species. While this scenario conflicts with that of Scherson et al. (2008), who found evidence for two dispersals from North to South America, the difference is attributable partly to differing resolution of clades (neither South American clade was recovered here) but especially to improved sampling in this study, including several South American taxa never studied before. Overall, this timeline and the predominant north-to-south direction are consistent with current understanding of the history of amphitropical disjunctions in plants (reviewed in Simpson et al., 2017). Based on our current sampling of this clade, only a single species of Neo-Astragalus left North America, with *Astragalus polaris* extending from Alaska to the Russian Far East.

Approximately 15 North American species, generally of more mesic environments (Barneby 1964), fall outside the Neo-Astragalus clade. These examples of secondary connections with North America, which include both endemics and species with large Northern Hemisphere distributions, are primarily descended from broad-ranging ancestors distributed in boreal regions, and unlike Neo-Astragalus, date to no earlier than the early Pleistocene, as predicted respectively by Barneby (1964) and Wojciechowski et al. (1999).

### Phylogenetic signal in quantitative niche attributes

All measured niche attributes showed strong evidence of phylogenetic niche conservatism (all highly significant, *λ* > 0.5 in all cases; see Supplementary Table S2). While strong phylogenetic structuring may be the rule for niche attributes, in most cases this mostly involved small, recently diverged clades (Fig. 2; Supplementary Fig. S3; cf. Wiens et al. [2010] for discussion of niche conservatism and phylogenetic scale). However, ancestral reconstruction (Fig. 2a–b; performed under the favored OU model) revealed that diurnal temperature range and particularly isothermality (BIO2 and BIO3) were remarkable in clearly delimiting the Neo-Astragalus clade of the Americas from Eurasian clades (Fig. 2b); no other niche attributes corresponded so strongly with geography. While strong diurnal temperature swings are associated with arid environments, this phylogenetic distribution was highly distinct from that seen for aridity, temperature, and precipitation (Fig. 2a; Supplementary Fig. S3). Likewise, elevation (Supplementary Fig. S3), although an important factor, does not delimit the major clades of *Astragalus.* Biogeographic region alone explains a remarkable 51.8 and 51.7% of the variance in diurnal range and isothermality, respectively (ANOVA adjusted *R^2^*). Sampling random cells from the predictor layers in the range of Neo-Astragalus and the Eurasian clades as a null demonstrates that this difference cannot be attributed to overall continental climatic differences in these variables (*t-*test, BIO2: *p* = 0.6394; BIO3: *p* = 0.4816).

**Fig. 2.**
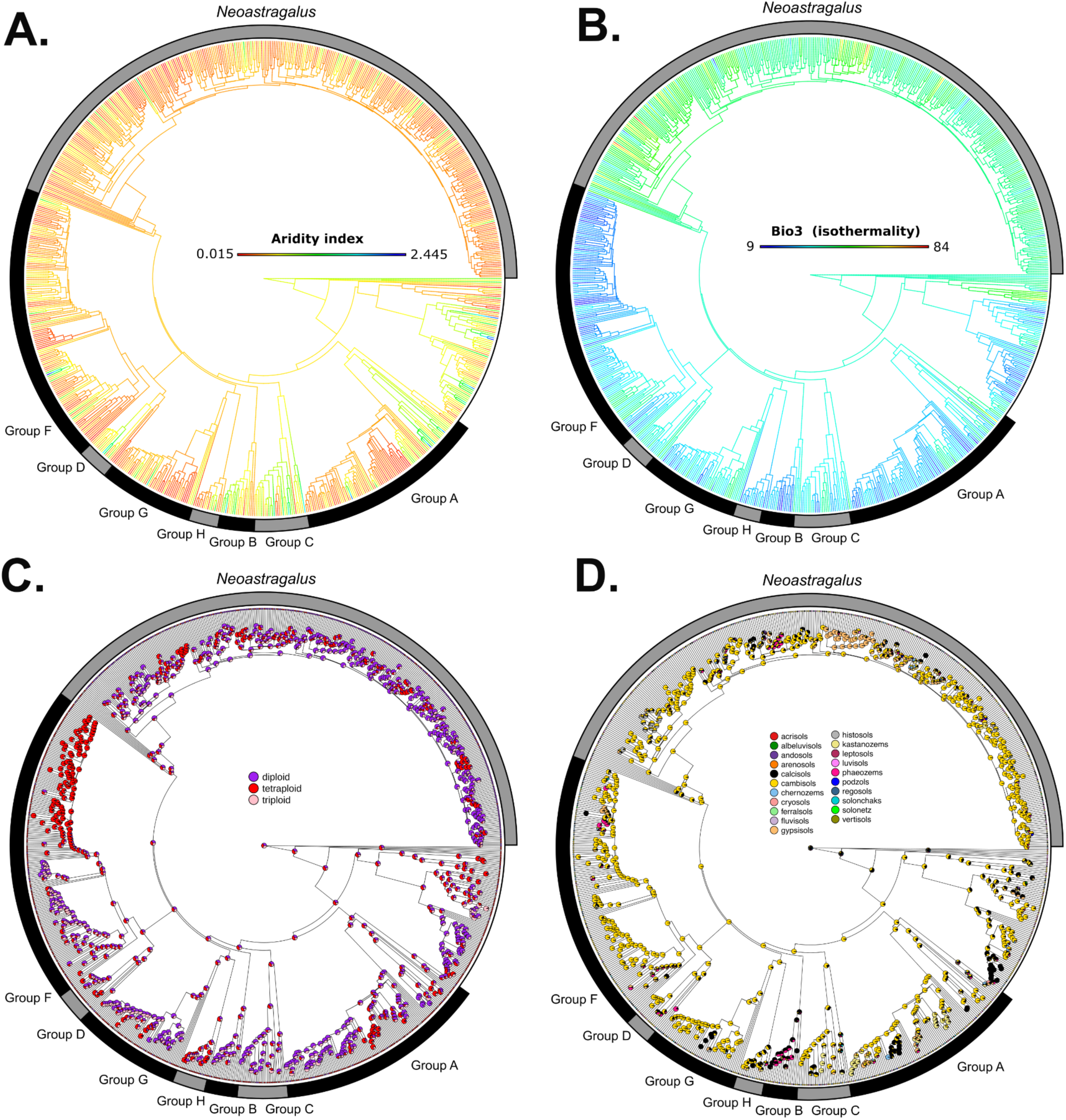
Ancestral reconstruction results for aridity (A), isothermality (B), ploidy (C), and soil type (D). Further reconstructions are available in Supplementary Fig. S3. Clade names follow Fig. 1.

Since environmental trends were also associated with clade membership (Fig. 2a) as well as biogeographic domain, and could have simply been driven by phylogenetic structuring of distributions, we performed two-way MANOVA (here, including only the quantitative niche attributes), showing that taxonomic group (following the *Astragalus of World* database, *p* < 2e-16) and biogeographic domain (*p* < 2e-16) have independent explanatory power for niche occupancy, as does their interaction (*p* = 0.0007413). This suggests that there is both clade- and area-based niche specialization.

### Phylogenetic signal in soil type

The only other environmental characteristic that delimited Neo-Astragalus from Eurasian species and showed similar deep-level phylogenetic structuring to BIO2 and BIO3 was FAO soil type classification, which is diagnosed on the basis of physical and chemical characteristics that are rooted primarily in soil formation processes such as erosion and alluvial deposition. Direct measures of soil fertility such as carbon and nitrogen content did not show similar phylogenetic structuring, suggesting this result was independent of direct nutrient challenge. Overall, calcisols, cambisols, chernozems, kastanozems, leptosols, luvisols, and regosols, most of which are poor soils of northern and arid regions, account for nearly all species diversity (Fig. 3e–f). Compared to Eurasian species, Neo-Astragalus is highly enriched for calcisols (defined partly by high calcium content and characteristic of the southwestern arid US), kastanozems (a humus-rich soil associated with grassland), and luvisols (a typical clay-rich temperate soil type; Fig. 3e). Eurasian species primarily occur in cambisols and regosols (Fig. 3f), soils associated with high erosion that are also found to a lesser extent in Neo-Astragalus. Cambisols, the most common soil association for the genus and a common soil type in cold and arid regions, are reconstructed with high probability as ancestral in *Astragalus* and *Oxytropis*, their MRCAs, and the entire backbone of *Astragalus* including Neo-Astragalus and all other major named clades (Fig. 2d).

**Fig. 3.**
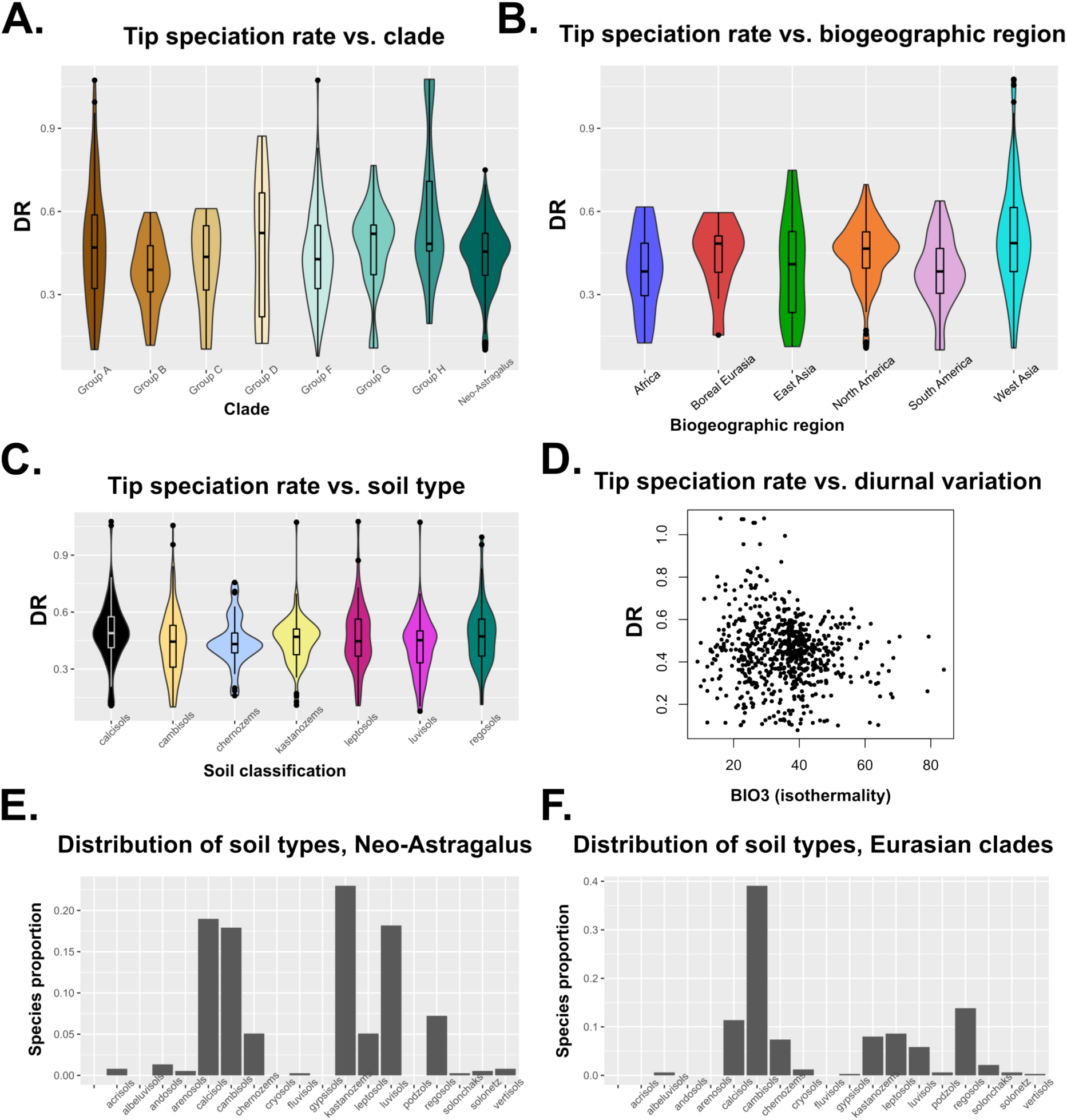
(**A-D**) Distribution of tip speciation rates, vs. named clade (A), vs. biogeographic region (B; colors follow Fig. 1), vs. soil type (C; colors follow Fig. 2d), and vs. isothermality (D). (**E-F**) Distribution of species occupancy of soil types for Neo-Astragalus (E) and for the Eurasian clades (F).

### Chromosome reconstruction

Chromosome reconstructions in ChromEvol (Fig. 1, Supplementary Fig. S4) definitively supported a base number of *n* = 8 for the Astragalean clade. This base number was largely retained throughout the Eastern Hemisphere grade and outgroup genera. The exceptions, considering only internal phylogenetic nodes, are a shift (with low posterior probability) to *n* = 6 in a small clade of *Oxytropis* and a shift to *n* = 7 in a small clade within *Astragalus* Group C. Finally, a shift to *n =* 11 occurred in the ancestor of Neo-Astragalus, which was followed by a shift to *n* = 12 in a clade of South American species and three clades of North American species, and to *n =* 13 in a clade of South and North American species (Supplemental Fig. S4). ChromEvol also detected numerous ancestral polyploidization events. These were distributed in small clades in *Oxytropis* and in *Astragalus* Groups B, G, and D (Supplemental Fig. S4).

### Ploidy reconstruction

Ploidy as reconstructed from sequence data (Fig. 2c) showed strong phylogenetic signal, but unlike the result recovered in ChromEvol, the reconstruction across the backbone of *Astragalus* was indecisive. Diploidy was reconstructed across Neo-Astragalus and a small subclade of *Astragalus* Group F. Ancestral reconstruction of the shift between ploidy calls (Fig. 2c) was uncertain but with a slightly greater likelihood of ancestral polyploidy followed by a shift to diploidy that most likely occurred in a subclade of Neo-Astragalus. As well as being biologically unreasonable (see Discussion for a potential interpretation), this scenario disagrees with model-based chromosome reconstructions based on chromosome counts (above) which point to ploidy shifts primarily in Eurasian species.

### Diversification

The hypothesis of uniform diversification regimes was definitively rejected (posterior probability ∼0 for zero shifts). Rates varied through time, with a shallow peak in the late Miocene (∼10 mya) and a strong peak at the Plio-Pleistocene boundary (∼2.5 mya; Fig. 1). The former burst is approximately contemporaneous with the divergence of Groups A–H and Neo-Astragalus, while the latter represents nested radiations early in the history of each major clade (Fig. 1). While eight shifts were recovered in the best diversification regime configuration (Fig. 1), these were not clearly associated with any attributes studied here. BIO2 (BAMM STRAPP *p* = 0.094; ESSim *p* = 0.7373) and BIO3 (STRAPP *p* = 0.056; ESSim *p* = 0.5435; Fig. 3d), the environmental characteristics closely associated with Neo-Astragalus, were not significantly associated with diversification. Soil characteristics also were not associated with diversification (STRAPP *p* = 0.07; FiSSE *p* = 0.9391, the latter with soil types recoded as calcisols vs. other soil types to yield a binary contrast; Fig. 3c). Ploidal levels did not differ in diversification rate, whether based on chromosome counts (STRAPP *p* = 0.799; here counts > *n* = 13 considered polyploid following the ChromEvol result) or sequence data (STRAPP *p* = 0.096, FiSSE *p* = 0.2408). Finally, tip speciation rates showed no obvious relationship with biogeographic region (Fig. 3b) and did not differ among the major recognized clades of *Astragalus* (Fig. 3a). Overall, despite BAMM detecting bursts of diversification, they are seen in most of the species-rich clades (Fig. 1) and resulted in similarly high rates of diversification across *Astragalus* (Supplementary Fig. S2). The only dataset significant with respect to diversification was chromosome number (STRAPP *p* = 0.027; note no chromosome shifts occur directly on branches experiencing diversification bursts except for the stem lineage of Neo-Astragalus: Fig. 1). These results are consistent with evidence for variable but overall high diversification rates in *Astragalus* that are not obviously linked to particular ecological strategies or other aspects investigated here, consistent with the idea of a non-adaptive radiation (Givnish, 2015).

## DISCUSSION

### *Biogeography of* Astragalus

As suggested by Maassoumi and Ashouri (2022), West Asia and especially the area comprising modern-day Iran, is the center of Eurasian *Astragalus* diversity, both geographically and in terms of species richness and diversity of major lineages. Our results add to this conclusion by suggesting West Asia as the ancestral area and as an essential crossroads for achieving its modern-day distribution. Dispersal to the Americas and other regions of Eurasia and Africa mostly depend on ancestors that were distributed in West Asia. Only transitions between North and South America are substantially independent of West Asia; otherwise movement into East Asia, Africa, and Boreal Eurasia occur from West Asia without exception in the biogeographic reconstruction.

We found more complex connections between North and South America than that recovered in previous studies. It has been long known that South American species are quite similar to North American species, seemingly more so than to each other, and certainly much more so than to the more numerous central Asian relatives (Johnston, 1947). This problem was previously studied by Scherson et al. (2008), who suggested a North American origin followed by two dispersals to South America. However, this study was limited by sampling (only 44 species of Neo-Astragalus) and did not use statistical biogeographic methods; likewise studies sampling standard Sanger markers have generally suffered from limited resolution caused by very low nucleotide divergence (Wojciechowski et al., 1999). Largely by virtue of much denser taxon sampling, although still with challenges regarding statistical support for subclades in Neo-Astragalus, we have been able to recover a more complex story, favoring vicariance rather than a solely North American scenario for the earliest history of the clade and supporting a larger number of transitions from North to South America. Otherwise, our results are similar to what was expected for *Astragalus*, with a decisive West Asian origin.

### *Climate evolution in* Astragalus

The most clear and surprising result of our investigation of niche attributes was the importance of diurnal temperature variation and its close tracking of biogeographic provinces. Given that we did not find a direct association of diurnal temperature range with ploidy or diversification, biogeographic area remains as the only significant associate with this niche attribute. While infrequently studied in plants, the importance of diurnal temperature range for determining plant distributions has been emphasized in previous studies. Ramsay (2001) and Zhao et al. (2018) found diurnal temperature variation to be an important predictor of plant growth form, a result they attributed to physiological stress adaptation; likewise, there are interactions between diurnal temperature and plant germination (Liu et al., 2013) and with plant size (Myster and Moe, 1995). In some investigations, latitudinal diversity patterns have been partly attributed to diurnal temperature variation in plants and microbes, attributed to filtering out species with narrow physiological tolerances within highly variable biomes (Hu et al., 2019; Lancaster and Humphreys, 2020). At the level of species rather than clades, important variables identified in the *Astragalus* of Eurasia have focused on seasonal rather than diurnal variation (Yang et al., 2020) or soil and topographical parameters (Safaei et al., 2018; Aghajanlou et al., 2021; Baumberger et al., 2021), so phylogenetic scale is an important consideration (Graham et al., 2018). Studies in Neo-Astragalus have focused on temperature seasonality but also identified diurnal variation; Jones et al. (2021) identified BIO4 as the most important PCA loading in the *A. sabulosus* group of the Colorado Plateau; similarly BIO4 was the second-most important variable in a distribution modeling study of *A. utahensis* but diurnal temperature variation was also an important predictor (Baer and Maron, 2020).

Our *Astragalus* results indicate the importance of diurnal variation, but remain as a surprise for two reasons. First, the strict biogeographic divide is remarkable and does not represent an equivalent divide in the background environmental variation of the Eastern and Western Hemispheres. This suggests that dispersal to the Americas was associated with niche specialization. Second, the primary correlate identified in the literature with diurnal variation is growth form, with high diurnal variation classically associated with low, cushion-like growth forms (Ramsay, 2001). *Astragalus*, by contrast, is remarkable for its morphological conservatism (Sanderson and Wojciechowski, 1996), and therefore has no corresponding growth-form contrast between the Hemispheres. The exact physiological adaptations relating to diurnal temperature variation occurring in *Astragalus* are unclear; nevertheless, the status of this clade as including some of the most important nodulating plants in its range suggests a photosynthetic explanation rooted in water stress. As argued in several previous works (Adams et al., 2016; Pellegrini et al., 2016; Doby et al., 2022), plants engaging in root nodule symbiosis exist on a distinct area of the leaf economic spectrum as embodied by their differing nitrogen investments (McKey, 1994; Crews, 1999). High nitrogen investment in photosynthesis in legumes appears to relate primarily to confronting water stress (Adams et al., 2016), suggesting that nodulation may serve as part of a spectrum of adaptations in *Astragalus* that explains their success in distinctive freezing arid areas.

### Soil biology

This study represents the first in-depth investigation of soil biology across *Astragalus* (see Wang and Wang, 2013; Wang et al., 2021 for studies on individual species), and indeed, to our knowledge, the first broad-scale phylogenetic investigation of recognized FAO soil classifications in a large number of plant species. Aside from diurnal temperature, soil classification (which captures primarily how soil formed rather than nutrient availability) turned out to be one of the most phylogenetically conserved habitat attributes, much more so than individual chemical attributes like carbon and nitrogen (which have been identified as important in *Astragalus* at the species rather than clade level; cf. Safaei et al., 2018; Yang et al., 2020).

Edaphic ecology, while still understudied (Rajakaruna, 2004, 2018; Anacker, 2014; Hulshof and Spasojevic, 2020), is often singled out as one of the most important biogeographic factors in plants (Raven, 1964; Axelrod, 1972). While legume biogeography is most often understood in terms of biomes and continents (Lewis et al., 2005), as nodulating plants, edaphic ecology is also of clear importance to legume distributions (Sprent et al., 2017). Soil rhizosphere environments, and the physiological limitations that soil environments impress upon rhizosphere bacteria and fungi, represent one potential constraint on plant distributions (Parker et al., 2006; Simonsen et al., 2017). Spatially robust microbiome studies on nodulating plants remain few; one study performed in *Astragalus* considered soil properties but found geographic distance the most important predictor of biogeographic structure for rhizospheres (Li et al., 2021). Nevertheless, evidence from other legume species suggests that edaphic ecology may be a key factor dictating symbiotic choice (Rathi et al., 2018). Besides symbiont choice, the edaphic setting interacts with water stress because soils differ in their water retention characteristics, potentially leading to an increased importance of root nodule symbioses. Given the few large-scale studies of edaphic characteristics in nodulating plants (Siniscalchi et al., 2022), progress documenting basic edaphic specializations are needed first if we are to make progress in understanding major contrasts such as that seen in *Astragalus* and test whether this can be attributed to differences in root nodule symbiosis or other factors.

### *Chromosome variation in* Astragalus

Similarly to previous investigations, this study showed base chromosome numbers are closely associated with biogeographic patterns in *Astragalus*, with a major base number shift associated with radiation in the Americas and more limited base shifts associated with many South American species. There was marginal evidence for an association between chromosome number (but not ploidy) and diversification rates, but diversification shifts are not directly associated with base number shifts. Investigations into the relationships between chromosome evolution and either biogeography (e.g., Rice et al., 2019) or diversification (e.g., Landis et al., 2018) have yielded mixed results. Polyploidy in particular has often been seen as a major source of evolutionary innovation in the face of stress and aneuploidy, while less well studied, has similarly been associated with abiotic stress (Folk et al., 2020; Van de Peer et al., 2021). The major association between chromosomes and biogeographic province in *Astragalus* may similarly relate to survival of abiotic stress through mechanisms promoting genetic variation.

An outcome of broader interest is a strong discrepancy between ploidy as assessed by chromosome numbers and by sequence data; while we present these data for the benefit of other potential users it is likely that many novel nQuire determinations presented here in taxa that have never been assessed for chromosome number are incorrect. The use of sequence data has attracted interest as it may offer a path to inferring ploidal levels from herbarium specimens where living material is impossible to obtain (Viruel et al., 2019), although unfortunately these are the same situations where validation of ploidy results will be most challenging. Some of the discordance between the empirical and inferred estimates of ploidy may relate to the nQuire software itself and its application to our samples. First, nQuire cannot test for ploidal levels higher than tetraploid; thus, if some samples are pentaploid or greater (known in *Astragalus*; http://ccdb.tau.ac.il/), the software will not perform appropriately (Viruel et al. 2019). Second, distinguishing among alternative ploidies requires sufficient sequence coverage, which is dependent on the size and complexity of the genome under study but is at least 35-40X in benchmarking studies for nQuire, with demonstration of the software’s effectiveness conducted at a minimum of 100X (Weiß et al. 2018); coverage levels vary in this study among samples and loci, and insufficient or inconsistent coverage will make it impossible identify the correct ploidal level. Finally, nQuire was developed for ‘intraspecific polyploidy’ (Weiß et al., 2018), which is interpretable as autopolyploidy; both auto- and allopolyploidy occur in *Astragalus* (e.g., Doyle 2012; Záveská et al. 2019; Bagheri et al. 2022), and the presence of allopolyploids may compromise the ability of nQuire to distinguish among ploidal levels. Other studies have likewise found discordances between estimates of ploidy based on the nQuire pipeline and other methods, such as flow cytometry (e.g., Viruel et al. 2019; Jantzen et al. 2022).

One possible and relatively simple way to explain the discrepancy between chromosome counts and sequence-based ploidal determinations, although in need of verification with genomic data, would be diploidization partly via aneuploidy from a polyploid ancestor. Higher basic numbers as seen in Neo-Astragalus were often traditionally interpreted as old polyploids with subsequent aneuploid reduction (Grant 1981: p 367) A corollary prediction would be reduced genome size relative to ploidy-based expectations, although this is not currently testable due to a lack of genome size data in Neo-Astragalus (Pellicer and Leitch, 2020). Paralogy and loss specific to the phylogenetic loci employed here, which are primarily housekeeping loci with copy number likely under selection, would be another possibility. In addition to the predominance of diploid calls in Neo-Astragalus, this clade also concentrates the highest values of putative paralogous genes, as identified by HybPiper (Supplementary Fig. S6). This could suggest the occurrence of a confounding polyploidization-diploidization event in the history of this clade obscuring sequence-based ploidy calls, although we lack sufficient gene sampling to investigate this hypothesis. Overall, investigators should use caution in directly determining ploidal levels from sequence data due to genome reduction and other confounding processes as well as methodological assumptions, especially where determinations cannot be groundtruthed by chromosome counts, and more investigations are needed of this still relatively experimental approach.

### *Diversification of* Astragalus

Sanderson and Wojciechowski (1996) were the first to quantify diversification rates in *Astragalus* and related genera; their primary hypothesis was that the Astragalean clade was characterized by relatively high but uniform rates of diversification. The analyses they performed were more restricted than those considered here, primarily testing differences between genera rather than at the species level within genera. Nevertheless, the hypothesis of uniform diversification rates is rejected here both considering rates within *Astragalus* and between *Astragalus* and other genera. Early-diverging clades within *Astragalus* share diversification regimes with outgroup taxa, while the Coluteoid clade and *Oxytropis* each have distinct diversification rate regimes, as do numerous core clades of *Astragalus*.

The rates we found, with a mean of between 0.5 and 1 speciation event per lineage per million years depending on the period (Fig. 1) and rates as high as 7 speciation events/my (Supplementary Fig. S2), are in line with those identified previously for the genus (cf. Wojciechowski et al., 1999; Scherson et al., 2008) and on the high end of what is known for plants (reviewed in Lagomarsino et al., 2016), well-befitting the high diversity of the genus. Although the rates are high, the timeline of diversification is conventional for a temperate radiation (reviewed in Folk et al., 2020). Two peaks of diversification in the late Miocene and Plio-Pleistocene accord with earlier work in *Astragalus* (Azani et al., 2019) and are typical of other major temperate clades (Arakaki et al., 2011; Folk et al., 2019; Sun et al., 2020).

The present work could not identify any clear abiotic factors among temperature, precipitation, topography, or soil parameters that could be associated with diversification. While diversification appears to be relatively heterogeneous, nevertheless our overall scenario accords with Sanderson and Wojciechowski (1996) that *Astragalus* and the entire Astragalean clade have relatively high diversification rates. Sanderson and Wojciechowski (1996) appeal to parallel origins of growth form to withstand abiotic stress and similar origins of chemical adaptations against herbivory to explain the high diversity of the group, yet abiotic ecological factors at least appear to be rejected. As well as assessing biotic interactions and chemical or physiological aspects, future work should focus on broadening the phylogenetic scope outside the Astragalean clade, which may identify alternative drivers from what we have seen here (Graham et al., 2018), as Sanderson and Wojciechowski (1996) focused on the Astragalean clade as a whole.

### Astragalus, Oxytropis *and Astragalean clade relationships*

Most previous phylogenetic work in *Astragalus* has used few genetic loci and focused either on representative sampling (Wojciechowski et al., 1999; Scherson et al., 2008) or on more extensive sampling of focal clades (Scherson et al., 2008; Bartha et al., 2013; Azani et al., 2017b, 2019; Bagheri et al., 2017; Amini et al., 2019) and areas (Su et al., 2021). Our analysis primarily corroborates previous phylogenetic findings across *Astragalus* while greatly improving sampling of Eurasian species. The phylogenetic placements we recovered validate and better resolve the “Old-World groups” of Kazempour Osaloo et al. (2003, 2005). As mentioned in Results, while the delimitation of named clades is similar to previously published work, the well-supported backbone we recover is discordant with two recent studies that also successfully resolved backbone relationships with high support (Azani et al., 2017b; Su et al., 2021) and interestingly was more similar to earlier work (Kazempour Osaloo et al., 2005). Two explanations may account for discordant results compared to Azani et al. (2017b) and Su et al. (2021): first, the aforementioned studies sampled very few Neo-Astragalus species, thus lacking the sampling to confidently place Neo-Astragalus relative to Eastern Hemisphere clades, therefore possibly explaining the anomalous placement of Neo-Astragalus. Second, these studies share a focus on high-copy portions of plant genomes with special evolutionary properties, with Su et al. (2021) using whole plastid genomes and Azani et al. (2017b) using ITS (the nuclear internal transcribed spacer) and plastid *trnK*/*matK*; Kazempour Osaloo et al. (2005) only included ITS sequences. Plastid genes are uniparentally inherited, not recombinationally independent from one another, and are often treated as single genes for the purpose of phylogenetic analysis (Doyle 1992); likewise both ribosomal and chloroplast genes are subject to gene conversion (Baldwin et al. 1995). Hence, the previous primary use of nonrecombining genomic compartments with special inheritance properties underlines the value of alternative evidence from the nuclear genome. Future studies should investigate the potential for cytonuclear phylogenetic conflict, which may involve deep relationships (Sun et al., 2015; Stull et al., 2020, 2023), and which could be expected given that hybridization is thought to be present in *Astragalus* (Bartha et al., 2013).

Finer-scale clades largely correspond to sectional taxonomy with many exceptions (Supplementary Fig. S1), and results will be of service in revising subgeneric taxonomy with additional focused sampling. However, Barneby’s sectional classification of Neo-Astragalus is not supported, with many clades recovered with support that are discordant with sectional delimitation (see Supplementary Fig. S1). However, limited backbone support in Neo-Astragalus in this analysis means that much of the Neo-Astragalus tree remains unresolved despite the increase in marker and taxon sampling. These groups should be tested further with intensive phylogenomic analyses and additional marker sets to improve resolution.

Among our outgroups, we sampled 71 species of *Oxytropis*, a monophyletic closely related segregate of *Astragalus* more prevalent in the Arctic (Meyers, 2012; Sandanov et al., 2022) that has been studied previously using small numbers of chloroplast and nuclear ribosomal markers (Archambault and Strömvik, 2012; Meyers, 2012; Kholina et al., 2016, 2020; Tekpinar et al., 2016a; b), as well as phylogenomic data to a limited extent (Shahi Shavvon et al., 2017). Biogeography of the *Oxytropis* species of the Americas was investigated by Archambault and Strömvik (2012) and Tekpinar et al. (2016b); both studies suggested multiple dispersals to the Americas from Eurasia, but were limited by very little nucleotide variation and were unable to target specifically the origin of this dispersal. We found four dispersals to North America, two each in the two major *Oxytropis* clades recovered. Interestingly, two of them in one of the clades (MRCA[*Oxytropis pilosa*, *Oxytropis yunnanensis*]) accord with *Astragalus* in being dispersals from West Asia (Supplementary Fig. S5), but the remaining two dispersal events in the other clade are classic East Asia-eastern North America disjunctions (Wen, 1999), a biogeographic pattern never observed in *Astragalus.* Overall, *Oxytropis* would be the excellent subject of further work as the relationships we recovered do not accord well with subgeneric taxa (e.g., Malyshev, 2008; Archambault and Strömvik, 2012), but we lack the sampling to test this further.

In terms of subtribal relationships, our analysis divides the Astragalean clade (currently tribe Galegeae pro parte) into two sister taxa: *Astragalus* and the Coluteoid clade, a series of genera around and including *Oxytropis*; relationships in the Coluteoid clade are similar to Zhang et al. (2009). Our results support the segregation of the sister genera *Biserrula* and *Podlechiella* (Kazempour Osaloo et al., 2003), both primarily distributed in the Mediterranean basin, as well as the primarily north Asian *Phyllolobium*.

### Conclusions

Our results confidently identify West Asia as the area of origin of *Astragalus* and the key area for practically all major continental biogeographic events. Temperature niche, edaphic ecology, and chromosome processes are remarkably distinct in the Neo-Astragalus clade of the Americas, yet neither these nor biogeographic regions could be related clearly to variation in diversification rates.

Our major niche result, the biogeographic importance of diurnal temperature variation, has never been suspected in *Astragalus* and is not often studied in plants. Yet, like other factors, diurnal temperature variation does not have a clear relationship to diversification processes despite being closely associated with a species-rich clade. These results highlight the need to broadly study abiotic parameters at large scales in major radiations, and to explicitly test their (potential lack of) importance in the diversification process. Further study would clarify whether *Astragalus* corresponds to a non-adaptive radiation (Rundell and Price, 2009; Givnish, 2015), which it seems to match well on the basis of abiotic factors.

## Acknowledgments

This work was funded by the National Science Foundation, DEB-1916632.

## REFERENCES

Ackerfield, J., A. Susanna, V. Funk, D. Kelch, D. S. Park, A. H. Thornhill, B. Yildiz, et al. 2020. A prickly puzzle: Generic delimitations in the *Carduus* - *Cirsium* group (Compositae: Cardueae: Carduinae). Taxon 69: 715–738.

Adams, M. A., T. L. Turnbull, J. I. Sprent, and N. Buchmann. 2016. Legumes are different: Leaf nitrogen, photosynthesis, and water use efficiency. Proceedings of the National Academy of Sciences of the United States of America 113: 4098–4103.

Afkhami, M. E., D. Luke Mahler, J. H. Burns, M. G. Weber, M. F. Wojciechowski, J. Sprent, and S. Y. Strauss. 2018. Symbioses with nitrogen-fixing bacteria: nodulation and phylogenetic data across legume genera. Ecology 99: 502.

Aghajanlou, F., H. Mirdavoudi, M. Shojaee, E. Mac Sweeney, A. Mastinu, and P. Moradi. 2021. Rangeland management and ecological adaptation analysis model for *Astragalus curvirostris* Boiss. Horticulturae 7: 67.

Amini, E., S. Kazempour-Osaloo, A. A. Maassoumi, and H. Zare-Maivan. 2019. Phylogeny, biogeography and divergence times of *Astragalus* section *Incani* DC. (Fabaceae) inferred from nrDNA ITS and plastid *rpl32-trnL* (UAG) sequences. Nordic Journal of Botany 37: e02059.

Anacker, B. L. 2014. The nature of serpentine endemism. American Journal of Botany 101: 219–224.

Arakaki, M., P.-A. Christin, R. Nyffeler, A. Lendel, U. Eggli, R. M. Ogburn, E. Spriggs, et al. 2011. Contemporaneous and recent radiations of the world’s major succulent plant lineages. Proceedings of the National Academy of Sciences of the United States of America 108: 8379–8384.

Archambault, A., and M. V. Strömvik. 2012. Evolutionary relationships in *Oxytropis* species, as estimated from the nuclear ribosomal internal transcribed spacer (ITS) sequences point to multiple expansions into the Arctic. Botany 90: 770–779.

Ardley, J., and J. Sprent. 2021. Evolution and biogeography of actinorhizal plants and legumes: A comparison. The Journal of Ecology 109: 1098–1121.

Axelrod, D. I. 1972. Edaphic aridity as a factor in angiosperm evolution. The American Naturalist 106: 311–320.

Azani, N., M. Babineau, C. D. Bailey, H. Banks, A. R. Barbosa, R. B. Pinto, J. S. Boatwright, et al. 2017a. A new subfamily classification of the Leguminosae based on a taxonomically comprehensive phylogeny: The Legume Phylogeny Working Group (LPWG). Taxon 66: 44–77.

Azani, N., A. Bruneau, M. F. Wojciechowski, and S. Zarre. 2019. Miocene climate change as a driving force for multiple origins of annual species in *Astragalus* (Fabaceae, Papilionoideae). Molecular Phylogenetics and Evolution 137: 210–221.

Azani, N., A. Bruneau, M. F. Wojciechowski, and S. Zarre. 2017b. Molecular phylogenetics of annual *Astragalus* (Fabaceae) and its systematic implications. Botanical Journal of the Linnean Society. Linnean Society of London 184: 347–365.

Baer, K. C., and J. L. Maron. 2020. Ecological niche models display nonlinear relationships with abundance and demographic performance across the latitudinal distribution of *Astragalus utahensis* (Fabaceae). Ecology and Evolution 10: 8251–8264.

Bagheri, A., A. A. Maassoumi, M. R. Rahiminejad, J. Brassac, and F. R. Blattner. 2017. Molecular phylogeny and divergence times of *Astragalus* section *Hymenostegis*: An analysis of a rapidly diversifying species group in Fabaceae. Scientific reports 7: 14033.

Bagheri, A., A. Akhavan Roofigar, Z. Nemati, and F. R. Blattner. 2022. Genome size and chromosome number evaluation of *Astragalus* L. sect. Hymenostegis Bunge (Fabaceae). Plants 11: 435.

Baldwin, B. G., M. J. Sanderson, J. M. Porter, M. F. Wojciechowski, C. S. Campbell, and M. J. Donoghue. 1995. The ITS region of nuclear ribosomal DNA: A valuable source of evidence on angiosperm phylogeny. Annals of the Missouri Botanical Garden 82: 247–277.

Barneby, R. C. 1964. Atlas of North American *Astragalus*. Mem NY Bot Gard 13: 1–1188.

Bartha, L., N. Dragoş, A. Molnár V., and G. Sramkó. 2013. Molecular evidence for reticulate speciation in *Astragalus* (Fabaceae) as revealed by a case study from sect. *Dissitiflori*. Botany 91: 702–714.

Baumberger, T., A. Baumel, P.-J. Dumas, J. Ugo, L. Keller, E. Dumas, T. Tatoni, et al. 2021. Is a restricted niche the explanation for species vulnerability? Insights from a large field survey of *Astragalus tragacantha* L. (Fabaceae). Flora 283: 151902.

Crews, T. E. 1999. The presence of nitrogen fixing legumes in terrestrial communities: Evolutionary vs ecological considerations. Biogeochemistry 46: 233–246.

De Storme, N., and A. Mason. 2014. Plant speciation through chromosome instability and ploidy change: Cellular mechanisms, molecular factors and evolutionary relevance. Current Plant Biology 1: 10–33.

Dludlu, M. N., S. B. M. Chimphango, C. H. Stirton, and A. M. Muasya. 2018. Distinct edaphic habitats are occupied by discrete legume assemblages with unique indicator species in the Cape Peninsula of South Africa. Journal of Plant Ecology 11: 632–644.

Doby, J. R., D. Li, R. A. Folk, C. M. Siniscalchi, and R. P. Guralnick. 2022. Aridity drives phylogenetic diversity and species richness patterns of nitrogen-fixing plants in North America. Global Ecology and Biogeography 31: 1630–1642.

Doyle, J. J., and J. L. Doyle. 1987. A rapid DNA isolation procedure for small quantities of fresh leaf tissue. Phytochem. Bull. 19: 11–15.

Doyle, J. J. 1992. Gene trees and species trees: Molecular systematics as one-character taxonomy. Systematic Botany 17: 144–163.

Doyle, J. J. 2012. Polyploidy in Legumes. In P. S. Soltis, and D. E. Soltis [eds.], Polyploidy and Genome Evolution, 147–180. Springer Berlin Heidelberg, Berlin, Heidelberg.

Folk, R. A., H. R. Kates, R. LaFrance, D. E. Soltis, P. S. Soltis, and R. P. Guralnick. 2021a. High-throughput methods for efficiently building massive phylogenies from natural history collections. Applications in Plant Sciences 9: e11410.

Folk, R. A., C. M. Siniscalchi, and D. E. Soltis. 2020. Angiosperms at the edge: Extremity, diversity, and phylogeny. Plant, Cell & Environment 43: 2871–2893.

Folk, R. A., R. L. Stubbs, N. J. Engle-Wrye, D. E. Soltis, and Y. Okuyama. 2021b. Biogeography and habitat evolution of Saxifragaceae, with a revision of generic limits and a new tribal system. Taxon 70: 263–285.

Folk, R. A., R. L. Stubbs, M. E. Mort, N. Cellinese, J. M. Allen, P. S. Soltis, D. E. Soltis, and R. P. Guralnick. 2019. Rates of niche and phenotype evolution lag behind diversification in a temperate radiation. Proceedings of the National Academy of Sciences of the United States of America 116: 10874–10882.

Frodin, D. G. 2004. History and concepts of big plant genera. Taxon 53: 753–776.

GBIF.org (22 May 2020) GBIF Occurrence Download https://doi.org/10.15468/dl.yfddmm

Givnish, T. J. 2015. Adaptive radiation versus ‘radiation’ and ‘explosive diversification’: why conceptual distinctions are fundamental to understanding evolution. The New Phytologist 207: 297–303.

Glick, L., and I. Mayrose. 2014. ChromEvol: Assessing the pattern of chromosome number evolution and the inference of polyploidy along a phylogeny. Molecular biology and evolution 31: 1914–1922.

Graham, C. H., D. Storch, and A. Machac. 2018. Phylogenetic scale in ecology and evolution. Global Ecology and Biogeography 27: 175–187.

Grant, V. 1981. Plant Speciation. Columbia University Press, New York.

Hardion, L., P.-J. Dumas, F. Abdel-Samad, M. Bou Dagher Kharrat, B. Surina, L. Affre, F. Médail, et al. 2016. Geographical isolation caused the diversification of the Mediterranean thorny cushion-like *Astragalus* L. sect. *Tragacantha* DC. (Fabaceae). Molecular Phylogenetics and Evolution 97: 187– 195.

Harvey, M. G., and D. L. Rabosky. 2018. Continuous traits and speciation rates: Alternatives to state-dependent diversification models. Methods in Ecology and Evolution 9: 984–993.

Hengl, T., J. Mendes de Jesus, G. B. M. Heuvelink, M. Ruiperez Gonzalez, M. Kilibarda, A. Blagotić, W. Shangguan, et al. 2017. SoilGrids250m: Global gridded soil information based on machine learning. PloS One 12: e0169748.

Hennig, W. 1966. Phylogenetic Systematics. University of Illinois Press.

Hijmans, R. J., S. E. Cameron, J. L. Parra, P. G. Jones, and A. Jarvis. 2005. Very high resolution interpolated climate surfaces for global land areas. International Journal of Climatology 25: 1965– 1978.

Höhna, S., M. J. Landis, T. A. Heath, B. Boussau, N. Lartillot, B. R. Moore, J. P. Huelsenbeck, and F. Ronquist. 2016. RevBayes: Bayesian phylogenetic inference using graphical models and an interactive model-specification language. Systematic Biology 65: 726–736.

Hu, A., Y. Nie, G. Yu, C. Han, J. He, N. He, S. Liu, et al. 2019. Diurnal temperature variation and plants drive latitudinal patterns in seasonal dynamics of soil microbial community. Frontiers in Microbiology 10: 674.

Hulshof, C. M., and M. J. Spasojevic. 2020. The edaphic control of plant diversity. Global ecology and Biogeography 19: 320.

Jantzen, J. R., P. J. F. Guimarães, L. C. Pederneiras, A. L. F. Oliveira, D. E. Soltis, and P. S. Soltis. 2022. Phylogenomic analysis of *Tibouchina* s.s. (Melastomataceae) highlights the evolutionary complexity of Neotropical savannas. Botanical Journal of the Linnean Society 199: 372–411.

Jetz, W., G. H. Thomas, J. B. Joy, K. Hartmann, and A. O. Mooers. 2012. The global diversity of birds in space and time. Nature 491: 444–448.

Johnson, M. G., E. M. Gardner, Y. Liu, R. Medina, B. Goffinet, A. J. Shaw, N. J. C. Zerega, and N. J. Wickett. 2016. HybPiper: Extracting coding sequence and introns for phylogenetics from high-throughput sequencing reads using target enrichment. Applications in Plant Sciences 4: apps.1600016.

Johnston, I. M. 1947. *Astragalus* in Argentina, Bolivia and Chile. Journal of the Arnold Arboretum. Arnold Arboretum 28: 336–374.

Jones, M. R., D. E. Winkler, and R. Massatti. 2021. The demographic and ecological factors shaping diversification among rare *Astragalus* species. Diversity & Distributions 27: 1407–1421.

Kates, H. R., B. C. O’Meara, R. LaFrance, G. W. Stull, E. K. James, D. Conde, S. Liu, et al. 2022. Two shifts in evolutionary lability underlie independent gains and losses of root-nodule symbiosis in a single clade of plants. bioRxiv: 2022.07.31.502231.

Kazempour Osaloo, S., A. A. Maassoumi, and N. Murakami. 2003. Molecular systematics of the genus *Astragalus* L. (Fabaceae): Phylogenetic analyses of nuclear ribosomal DNA internal transcribed spacers and chloroplast gene ndhF sequences. Plant Systematics and Evolution 242: 1–32.

Kazempour Osaloo, S., A. A. Maassoumi, and N. Murakami. 2005. Molecular systematics of the Old World *Astragalus* (Fabaceae) as inferred from nrDNA ITS sequence data. Brittonia 57: 367–381.

Kholina, A. B., M. M. Kozyrenko, E. V. Artyukova, D. V. Sandanov, and E. A. Andrianova. 2016. Phylogenetic relationships of the species of *Oxytropis* DC. subg. *Oxytropis* and *Phacoxytropis* (Fabaceae) from Asian Russia inferred from the nucleotide sequence analysis of the intergenic spacers of the chloroplast genome. Russian Journal of Genetics 52: 780–793.

Kholina, A. B., M. M. Kozyrenko, E. V. Artyukova, V. V. Yakubov, M. G. Khoreva, E. A. Andrianova, and O. A. Mochalova. 2020. Phylogenetic relationships of *Oxytropis* section *Arctobia* of northeast Asia according to sequencing of the intergenic spacers of chloroplast and ITS of nuclear genomes. Russian Journal of Genetics 56: 1424–1434.

Lagomarsino, L. P., F. L. Condamine, A. Antonelli, A. Mulch, and C. C. Davis. 2016. The abiotic and biotic drivers of rapid diversification in Andean bellflowers (Campanulaceae). The New Phytologist 210: 1430–1442.

Lancaster, L. T., and A. M. Humphreys. 2020. Global variation in the thermal tolerances of plants. Proceedings of the National Academy of Sciences of the United States of America 117: 13580– 13587.

Landis, J. B., D. E. Soltis, Z. Li, H. E. Marx, M. S. Barker, D. C. Tank, and P. S. Soltis. 2018. Impact of whole-genome duplication events on diversification rates in angiosperms. American Journal of Botany 105: 348–363.

Ledingham, G. F. 1960. Chromosome numbers in *Astragalus* and *Oxytropis*. Canadian Journal of Genetics and Cytology 2: 119–128.

Levin, D. A. 1975. Pest pressure and recombination systems in plants. The American Naturalist 109: 437–451.

Lewis, G. P., B. Schrire, B. Mackinder, and M. Lock. 2005. Legumes of the World. Royal Botanic Gardens, Kew.

Lichter-Marck, I. H., and B. G. Baldwin. 2023. Edaphic specialization onto bare, rocky outcrops as a factor in the evolution of desert angiosperms. Proceedings of the National Academy of Sciences of the United States of America 120: e2214729120.

Li, H., and R. Durbin. 2009. Fast and accurate short read alignment with Burrows-Wheeler transform. Bioinformatics 25: 1754–1760.

Li, H., B. Handsaker, A. Wysoker, T. Fennell, J. Ruan, N. Homer, G. Marth, et al. 2009. The Sequence Alignment/Map format and SAMtools. Bioinformatics 25: 2078–2079.

Liu, K., J. M. Baskin, C. C. Baskin, H. Bu, G. Du, and M. Ma. 2013. Effect of diurnal fluctuating versus constant temperatures on germination of 445 species from the eastern Tibet Plateau. PloS One 8: e69364.

Li, Y., Y. Yang, T. ’e Wu, H. Zhang, G. Wei, and Z. Li. 2021. Rhizosphere bacterial and fungal spatial distribution and network pattern of *Astragalus mongholicus* in representative planting sites differ the bulk soil. Applied Soil Ecology 168: 104114.

Louca, S., and M. W. Pennell. 2020. Extant timetrees are consistent with a myriad of diversification histories. Nature 580: 502–505.

Maassoumi, A. A., and P. Ashouri. 2022. The hotspots and conservation gaps of the mega genus *Astragalus* (Fabaceae) in the Old-World. Biodiversity and Conservation 31: 2119–2139.

Malyshev, L. I. 2008. Phenetics of the subgenera and sections in the genus *Oxytropis* DC. (Fabaceae) bearing on ecology and phylogeny. Contemporary Problems of Ecology 1: 440–444.

Matzke, N. J. 2013. Probabilistic historical biogeography: new models for founder-event speciation, imperfect detection, and fossils allow improved accuracy and model-testing. Frontiers of Biogeography 5(4). https://doi.org/10.21425/F5FBG19694

McKey, D. 1994. Legumes and nitrogen: The evolutionary ecology of a nitrogen-demanding lifestyle. Advances in Legume Systematics Part 5, 211–228.

Meyers, Z. J. 2012. A contribution to the taxonomy and phylogeny of *Oxytropis* section *Arctobia* (Fabaceae) in North America. Masters. University of Alaska Fairbanks.

Moore, B. R., S. Höhna, M. R. May, B. Rannala, and J. P. Huelsenbeck. 2016. Critically evaluating the theory and performance of Bayesian analysis of macroevolutionary mixtures. Proceedings of the National Academy of Sciences of the United States of America 113: 9569–9574.

Myster, J., and R. Moe. 1995. Effect of diurnal temperature alternations on plant morphology in some greenhouse crops—a mini review. Scientia Horticulturae 62: 205–215.

Paradis, E., J. Claude, and K. Strimmer. 2004. APE: Analyses of Phylogenetics and Evolution in R language. Bioinformatics 20: 289–290.

Parker, M. A., W. Malek, and I. M. Parker. 2006. Growth of an invasive legume is symbiont limited in newly occupied habitats. Diversity & Distributions 12: 563–571.

Pellegrini, A. F. A., A. C. Staver, L. O. Hedin, T. Charles-Dominique, and A. Tourgee. 2016. Aridity, not fire, favors nitrogen-fixing plants across tropical savanna and forest biomes. Ecology 97: 2177–2183.

Pellicer, J., and I. J. Leitch. 2020. The Plant DNA C-values database (release 7.1): an updated online repository of plant genome size data for comparative studies. The New Phytologist 226: 301–305.

Plenk, K., W. Willner, O. N. Demina, M. Höhn, A. Kuzemko, K. Vassilev, and M. Kropf. 2020. Phylogeographic evidence for long-term persistence of the Eurasian steppe plant *Astragalus onobrychis* in the Pannonian region (eastern Central Europe). Flora 264: 151555.

Polhill, R. M. 1981. Galegeae. Advances in Legume Systematics: 367–370.

Rabosky, D. L. 2014. Automatic detection of key innovations, rate shifts, and diversity-dependence on phylogenetic trees. PloS one 9: e89543.

Rabosky, D. L., and E. E. Goldberg. 2017. FiSSE: A simple nonparametric test for the effects of a binary character on lineage diversification rates. Evolution 71: 1432–1442.

Rabosky, D. L., and E. E. Goldberg. 2015. Model inadequacy and mistaken inferences of trait-dependent speciation. Systematic Biology 64: 340–355.

Rabosky, D. L., and H. Huang. 2016. A robust semi-parametric test for detecting trait-dependent diversification. Systematic Biology 65: 181–193.

Rajakaruna, N. 2018. Lessons on evolution from the study of edaphic specialization. The Botanical Review 84: 39–78.

Rajakaruna, N. 2004. The edaphic factor in the origin of plant species. International Geology Review 46: 471–478.

Ramsay, P. M. 2001. Diurnal temperature variation in the major growth forms of an Ecuadorian páramo plant community. In P. M. Ramsay [ed.], The Ecology of Volcán Chiles: High-altitude Ecosystems on the Ecuador-Colombia Border, 101–112.

Rathi, S., N. Tak, G. Bissa, B. Chouhan, A. Ojha, D. Adhikari, S. K. Barik, et al. 2018. Selection of *Bradyrhizobium* or *Ensifer* symbionts by the native Indian caesalpinioid legume *Chamaecrista pumila* depends on soil pH and other edaphic and climatic factors. FEMS Microbiology Ecology 94: fiy180.

Raven, P. H. 1964. Catastrophic selection and edaphic endemism. Evolution 18: 336–338.

Ree, R. H., and I. Sanmartín. 2018. Conceptual and statistical problems with the DEC+J model of founder-event speciation and its comparison with DEC via model selection. Journal of Biogeography 45: 741–749.

Rice, A., L. Glick, S. Abadi, M. Einhorn, N. M. Kopelman, A. Salman-Minkov, J. Mayzel, et al. 2015. The Chromosome Counts Database (CCDB) - a community resource of plant chromosome numbers. The New Phytologist 206: 19–26.

Rice, A., P. Šmarda, M. Novosolov, M. Drori, L. Glick, N. Sabath, S. Meiri, et al. 2019. The global biogeography of polyploid plants. Nature Ecology & Evolution 3: 265–273.

Rundell, R. J., and T. D. Price. 2009. Adaptive radiation, nonadaptive radiation, ecological speciation and nonecological speciation. Trends in Ecology & Evolution 24: 394–399.

Safaei, M., M. Tarkesh, H. Bashari, and M. Bassiri. 2018. Modeling potential habitat of *Astragalus verus* Olivier for conservation decisions: A comparison of three correlative models. Flora 242: 61–69.

Samad, F. A., A. Baumel, M. Juin, D. Pavon, S. Siljak-Yakovlev, F. Médail, and M. Bou Dagher Kharrat. 2014. Phylogenetic diversity and genome sizes of *Astragalus* (Fabaceae) in the Lebanon biogeographical crossroad. Plant Systematics and evolution 300: 819–830.

Sandanov, D. V., A. S. Dugarova, E. P. Brianskaia, I. Y. Selyutina, N. I. Makunina, S. V. Dudov, V. V. Chepinoga, and Z. Wang. 2022. Diversity and distribution of *Oxytropis* DC. (Fabaceae) species in Asian Russia. Biodiversity Data Journal 10: e78666.

Sanderson, M. J., and M. F. Wojciechowski. 1996. Diversification rates in a temperate legume clade: Are there ‘so many species’ of *Astragalus* (Fabaceae)? American Journal of Botany 83: 1488–1502.

Scherson, R. A., R. Vidal, and M. J. Sanderson. 2008. Phylogeny, biogeography, and rates of diversification of New World *Astragalus* (Leguminosae) with an emphasis on South American radiations. American Journal of Botany 95: 1030–1039.

Shahi Shavvon, R., S. Kazempour Osaloo, A. A. Maassoumii, F. Moharrek, S. Karaman Erkul, A. R. Lemmon, E. M. Lemmon, et al. 2017. Increasing phylogenetic support for explosively radiating taxa: The promise of high-throughput sequencing for *Oxytropis* (Fabaceae). Journal of Systematics and Evolution 55: 385–404.

Simonsen, A. K., R. Dinnage, L. G. Barrett, S. M. Prober, and P. H. Thrall. 2017. Symbiosis limits establishment of legumes outside their native range at a global scale. Nature Communications 8: 14790.

Simpson, M. G., L. A. Johnson, T. Villaverde, and C. M. Guilliams. 2017. American amphitropical disjuncts: Perspectives from vascular plant analyses and prospects for future research. American Journal of Botany 104: 1600–1650.

Siniscalchi, C. M., H. R. Kates, P. S. Soltis, D. E. Soltis, R. P. Guralnick, and R. A. Folk. 2022. Testing the evolutionary drivers of nitrogen-fixing symbioses in challenging soil environments. bioRxiv: 2022.09.27.509719.

Smith, S. A., and B. C. O’Meara. 2012. treePL: Divergence time estimation using penalized likelihood for large phylogenies. Bioinformatics 28: 2689–2690.

Spellenberg, R. 1976. Chromosome numbers and their cytotaxonomic significance for North American *Astragalus* (Fabaceae). Taxon 25: 463–476.

Sprent, J. I., J. Ardley, and E. K. James. 2017. Biogeography of nodulated legumes and their nitrogen-fixing symbionts. The New Phytologist 215: 40–56.

Stamatakis, A. 2014. RAxML version 8: A tool for phylogenetic analysis and post-analysis of large phylogenies. Bioinformatics 30: 1312–1313.

Stull, G. W., K. K. Pham, P. S. Soltis, and D. E. Soltis. 2023. Deep reticulation: The long legacy of hybridization in vascular plant evolution. The Plant Journal 114: 743–766.

Stull, G. W., P. S. Soltis, D. E. Soltis, M. A. Gitzendanner, and S. A. Smith. 2020. Nuclear phylogenomic analyses of asterids conflict with plastome trees and support novel relationships among major lineages. American Journal of Botany 107: 790–805.

Su, C., L. Duan, P. Liu, J. Liu, Z. Chang, and J. Wen. 2021. Chloroplast phylogenomics and character evolution of eastern Asian *Astragalus* (Leguminosae): Tackling the phylogenetic structure of the largest genus of flowering plants in Asia. Molecular Phylogenetics and Evolution 156: 107025.

Sun, M., R. A. Folk, M. A. Gitzendanner, P. S. Soltis, Z. Chen, D. E. Soltis, and R. P. Guralnick. 2020. Recent accelerated diversification in rosids occurred outside the tropics. Nature Communications 11: 3333.

Sun, M., D. E. Soltis, P. S. Soltis, X. Zhu, J. G. Burleigh, and Z. Chen. 2015. Deep phylogenetic incongruence in the angiosperm clade Rosidae. Molecular Phylogenetics and Evolution 83: 156– 166.

Tamme, R., M. Pärtel, U. Kõljalg, L. Laanisto, J. Liira, Ü. Mander, M. Moora, et al. 2021. Global macroecology of nitrogen-fixing plants. Global Ecology and Biogeography: 30: 514–526.

Tekpinar, A., S. K. Erkul, Z. Aytaç, and Z. Kaya. 2016a. Phylogenetic relationships among native *Oxytropis* species in Turkey using the *trnL* intron, *trnL-F* IGS, and *trnV* intron cpDNA regions. Turkish Journal of Botany 40: 472–479.

Tekpinar, A., S. K. Erkul, Z. Aytaç, and Z. Kaya. 2016b. Phylogenetic relationships between *Oxytropis* DC. and *Astragalus* L. species native to an Old World diversity center inferred from nuclear ribosomal ITS and plastid matK gene sequences. Turkish Journal of Biology 40: 250–263.

Title, P. O., and D. L. Rabosky. 2019. Tip rates, phylogenies, and diversification: what are we estimating, and how good are the estimates? Methods Ecol. Evol.: 821–834.

Tribble, C. M., W. A. Freyman, M. J. Landis, J. Y. Lim, J. Barido-Sottani, B. T. Kopperud, S. Hӧhna, and M. R. May. 2022. RevGadgets: An R package for visualizing Bayesian phylogenetic analyses from RevBayes. Methods in Ecology and Evolution 13: 314–323.

Tuanmu, M.-N., and W. Jetz. 2014. A global 1-km consensus land-cover product for biodiversity and ecosystem modelling: Consensus land cover. Global Ecology and Biogeography 23: 1031–1045.

Van de Peer, Y., T.-L. Ashman, P. S. Soltis, and D. E. Soltis. 2021. Polyploidy: An evolutionary and ecological force in stressful times. The Plant Cell 33: 11–26.

Viruel, J., M. Conejero, O. Hidalgo, L. Pokorny, R. F. Powell, F. Forest, M. B. Kantar, et al. 2019. A target capture-based method to estimate ploidy from herbarium specimens. Frontiers in Plant Science 10: 937.

Viruel, J., O. Hidalgo, L. Pokorny, F. Forest, B. Gravendeel, P. Wilkin, and I. J. Leitch. 2023. A bioinformatic pipeline to estimate ploidy level from target capture sequence data obtained from herbarium specimens. In T. Heitkam, and S. Garcia [eds.], Plant Cytogenetics and Cytogenomics: Methods and Protocols, 115–126. Springer US, New York, NY.

Wang, Q., C. Chen, Z. Pang, C. Li, D. Wang, Q. Ma, and J. Wu. 2021. The role of the locoweed (*Astragalus variabilis* Bunge) in improving the soil properties of desert grasslands. Rangeland Journal 43: 47–54.

Wang, Z.-B., and Q.-Y. Wang. 2013. Cultivating erect milkvetch (*Astragalus adsurgens* Pall.) (Leguminosae) improved soil properties in Loess Hilly and Gullies in China. Journal of Integrative Agriculture 12: 1652–1658.

Weiß, C. L., M. Pais, L. M. Cano, S. Kamoun, and H. A. Burbano. 2018. nQuire: A statistical framework for ploidy estimation using next generation sequencing. BMC Bioinformatics 19: 122.

Weitemier, K., S. C. K. Straub, R. C. Cronn, M. Fishbein, R. Schmickl, A. McDonnell, and A. Liston. 2014. Hyb-Seq: Combining target enrichment and genome skimming for plant phylogenomics. Applications in Plant Sciences 2: apps.1400042.

Wen, J. 1999. Evolution of eastern Asian and eastern North American disjunct distributions in flowering plants. Annual Review of Ecology and Systematics 30: 421–455.

Wiens, J. J., D. D. Ackerly, A. P. Allen, B. L. Anacker, L. B. Buckley, H. V. Cornell, E. I. Damschen, et al. 2010. Niche conservatism as an emerging principle in ecology and conservation biology. Ecology Letters 13: 1310–1324.

Wojciechowski, M. F., M. Lavin, and M. J. Sanderson. 2004. A phylogeny of legumes (Leguminosae) based on analysis of the plastid *matK* gene resolves many well-supported subclades within the family. American Journal of Botany 91: 1846–1862.

Wojciechowski, M. F., M. J. Sanderson, B. G. Baldwin, and M. J. Donoghue. 1993. Monophyly of aneuploid *Astragalus* (Fabaceae): Evidence from nuclear ribosomal DNA internal transcribed spacer sequences. American Journal of Botany 80: 711–722.

Wojciechowski, M. F., M. J. Sanderson, and J.-M. Hu. 1999. Evidence on the monophyly of *Astragalus* (Fabaceae) and its major subgroups based on nuclear ribosomal DNA ITS and chloroplast DNA *trnL* intron data. Systematic Botany 24: 409–437.

Wojciechowski, M. F., M. J. Sanderson, K. P. Steele, and A. Liston. 2000. Molecular phylogeny of the ‘temperate herbaceous tribes’ of papilionoid legumes. Advances in Legume Systematics 9: 277–298.

Yang, M., Z. Li, L. Liu, A. Bo, C. Zhang, and M. Li. 2020. Ecological niche modeling of *Astragalus membranaceus* var. *mongholicus* medicinal plants in Inner Mongolia, China. Scientific Reports 10: 12482.

Záveská, E., C. Maylandt, O. Paun, C. Bertel, B. Frajman, The Steppe Consortium, and P. Schönswetter. 2019. Multiple auto- and allopolyploidisations marked the Pleistocene history of the widespread Eurasian steppe plant *Astragalus onobrychis* (Fabaceae). Molecular Phylogenetics and Evolution 139: 106572.

Zhang, C., M. Rabiee, E. Sayyari, and S. Mirarab. 2018. ASTRAL-III: Polynomial time species tree reconstruction from partially resolved gene trees. BMC Bioinformatics 19: 153.

Zhang, M., Y. Kang, L. Zhou, and D. Podlech. 2009. Phylogenetic origin of *Phyllolobium* with a further implication for diversification of *Astragalus* in China. Journal of Integrative Plant Biology 51: 889– 899.

Zhao, Y., H. Cao, W. Xu, G. Chen, J. Lian, Y. Du, and K. Ma. 2018. Contributions of precipitation and temperature to the large scale geographic distribution of fleshy-fruited plant species: Growth form matters. Scientific Reports 8: 17017.

