## Supplement for "Anatomy of a mega-radiation: Biogeography and niche evolution in *Astragalus*"

**Fig. S1.** Phylogeny, broken in three parts for display as indicated. Branch lengths represent coalescent units, and are arbitrarily plotted as 1 for tips. Support values plotted on branches represent local posterior probabilities (LPP), and are only plotted where they exceed 0.5. Tip labels in *Astragalus* are plotted by group, then section, then binomial. E.g., *Astragalus mongholicus* is within *Astragalus* sect. *Nuculiella* of group A, and therefore is rendered as "GA_Nuculiella_Astragalus_chinensis".

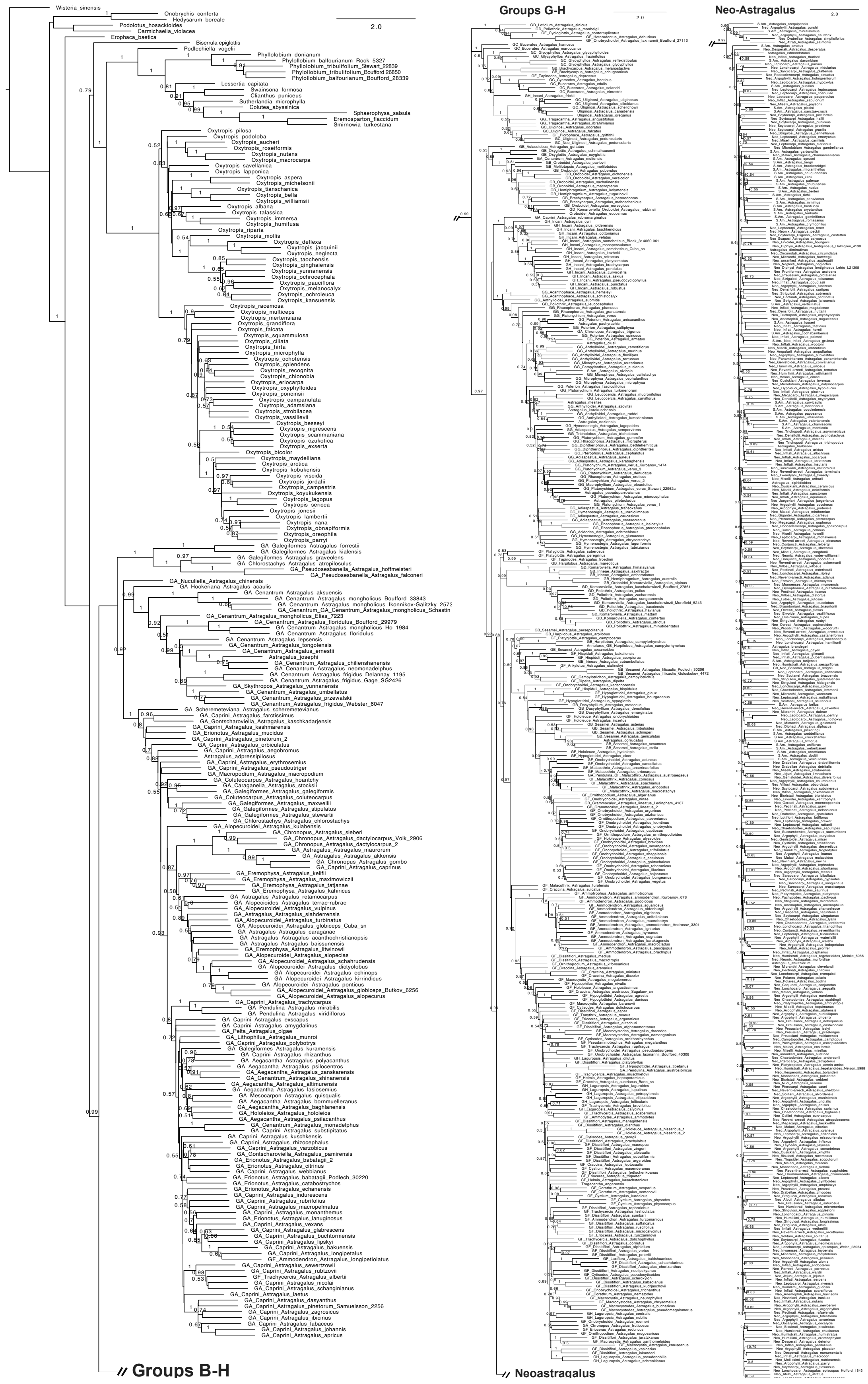

Groups G-H
Neo-Astragalus
2.0
Groups B-H
Neoastragalus

**Fig. S2.** Plot of BAMM net diversification rate shifts. Red dots indicate shifts in BAMM rate regime; branch colors represent net diversification per the legend, in events per million years.

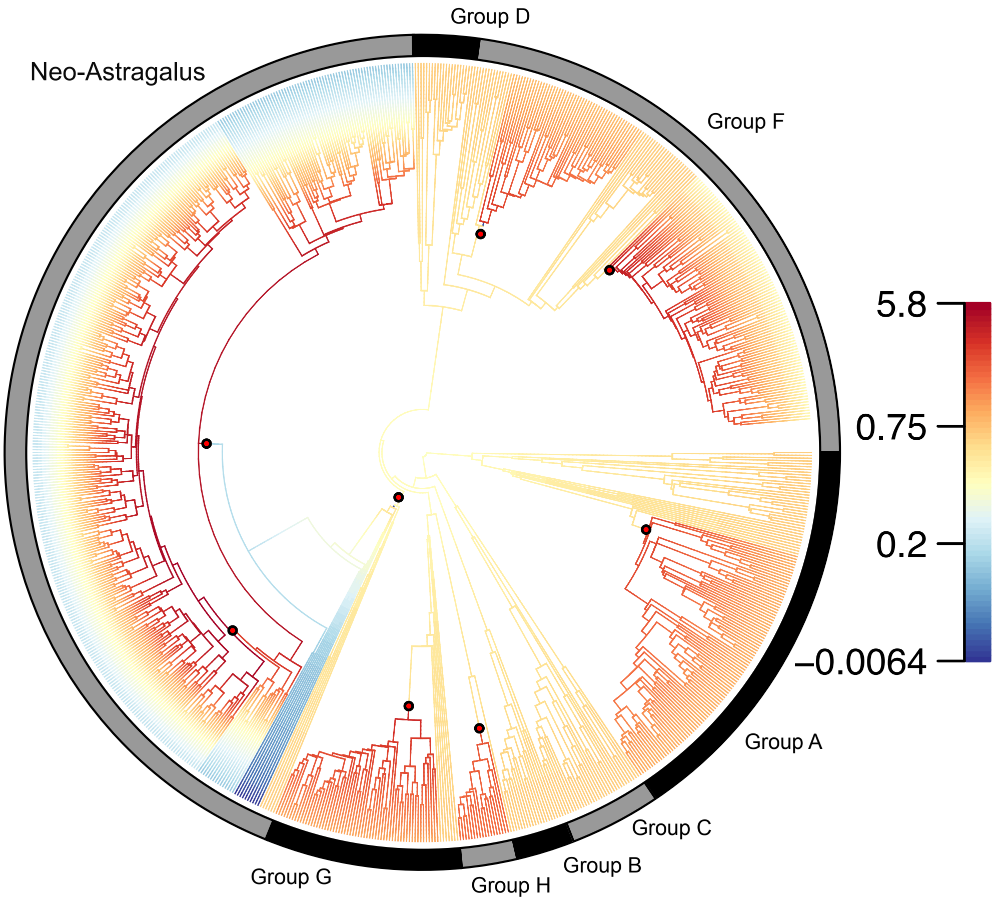

**Fig. S3.** Ancestral reconstructions for the 9 predictor variables not shown in the main text.

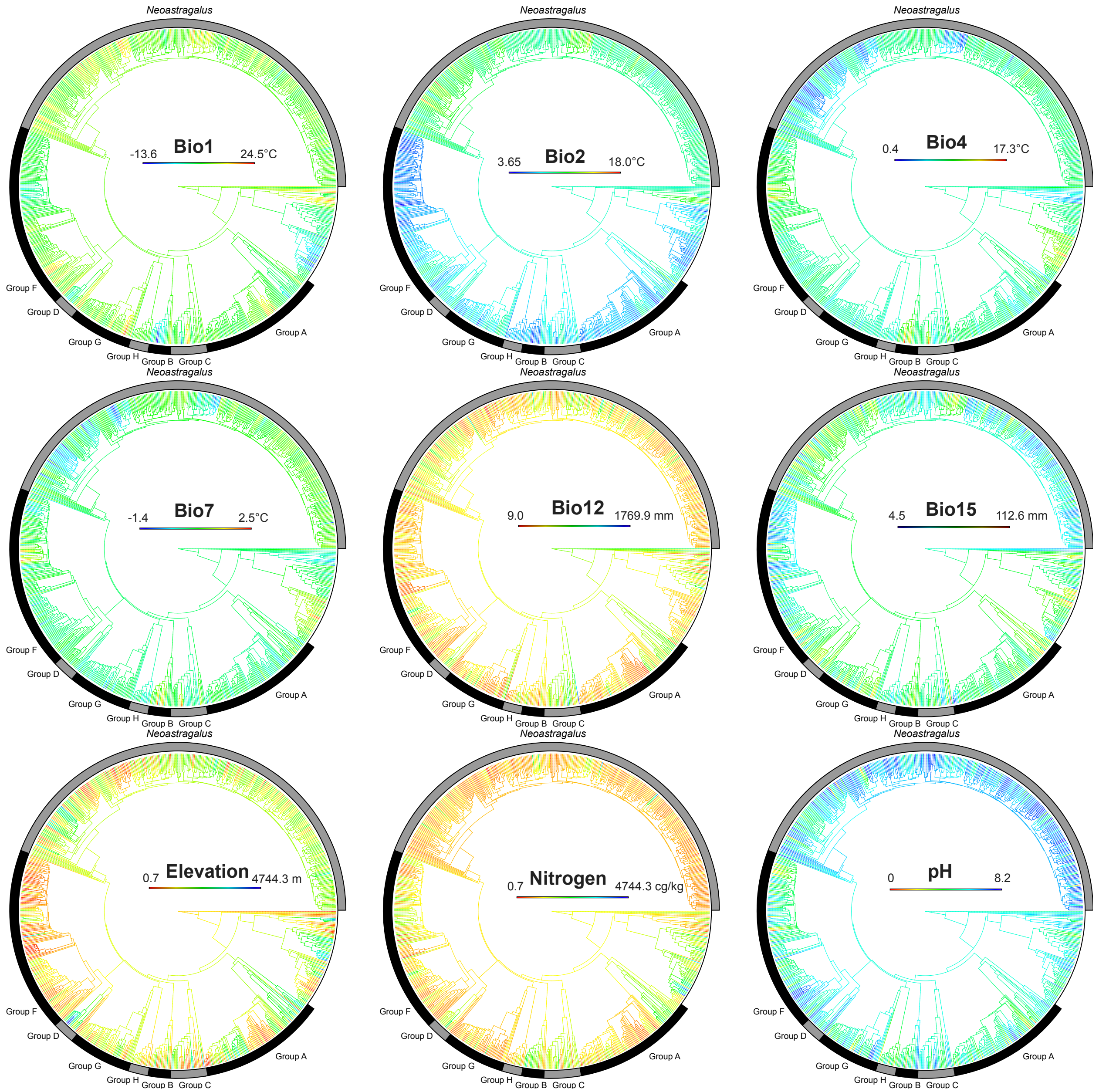

**Fig. S4.** Phylogeny, broken in two parts for display as indicated. Branch shape sizes and labels indicate reconstructed chromosome number; branch colors indicate posterior probability, with redder colors indicating higher values.

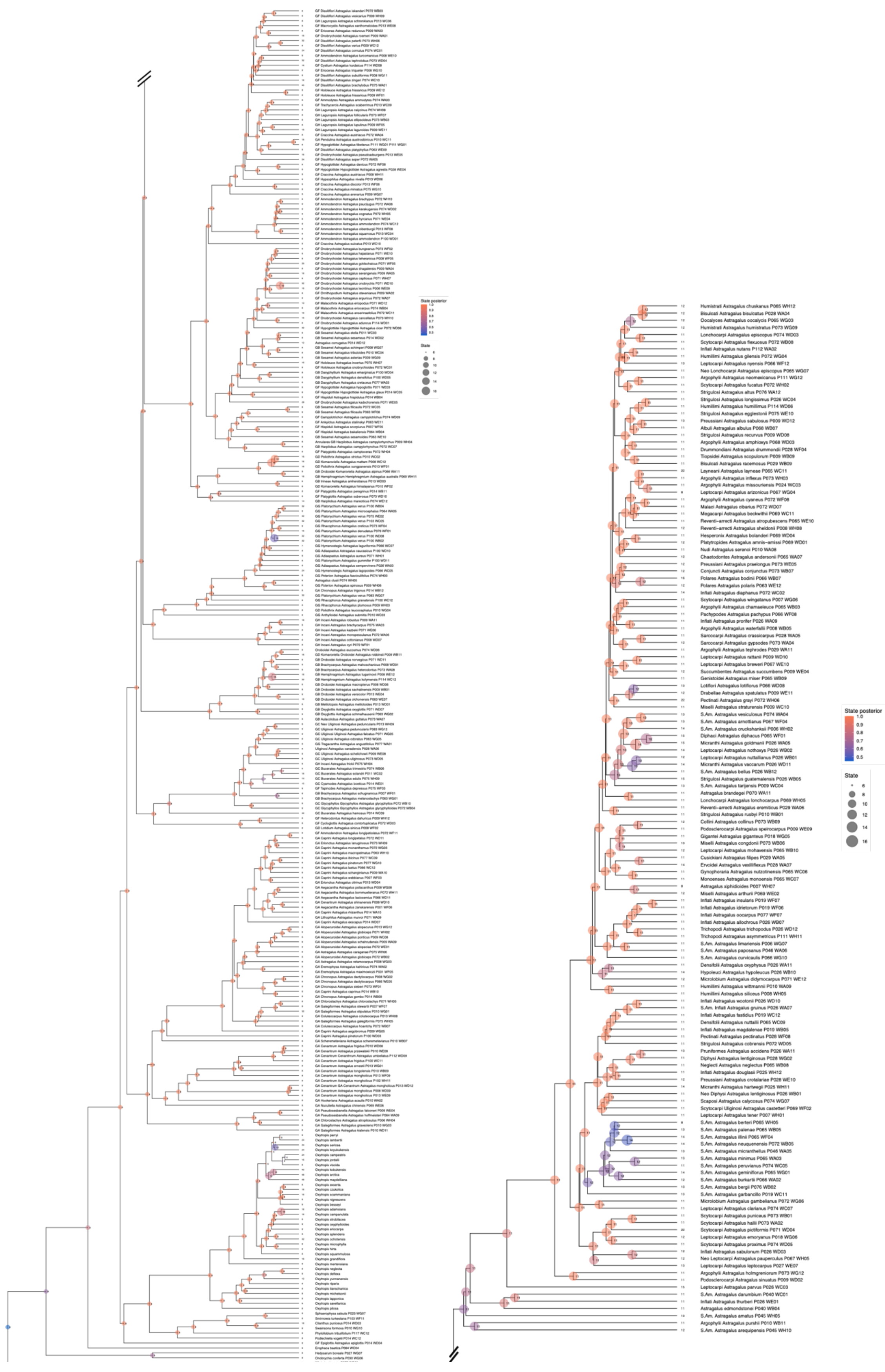

State posterior
1.0
0.9
0.8
0.7
0.6
0.5
State
6
8
10
12
14
16

**Fig. S5.** Complete biogeographic result (cf. Fig. 1 in main text).

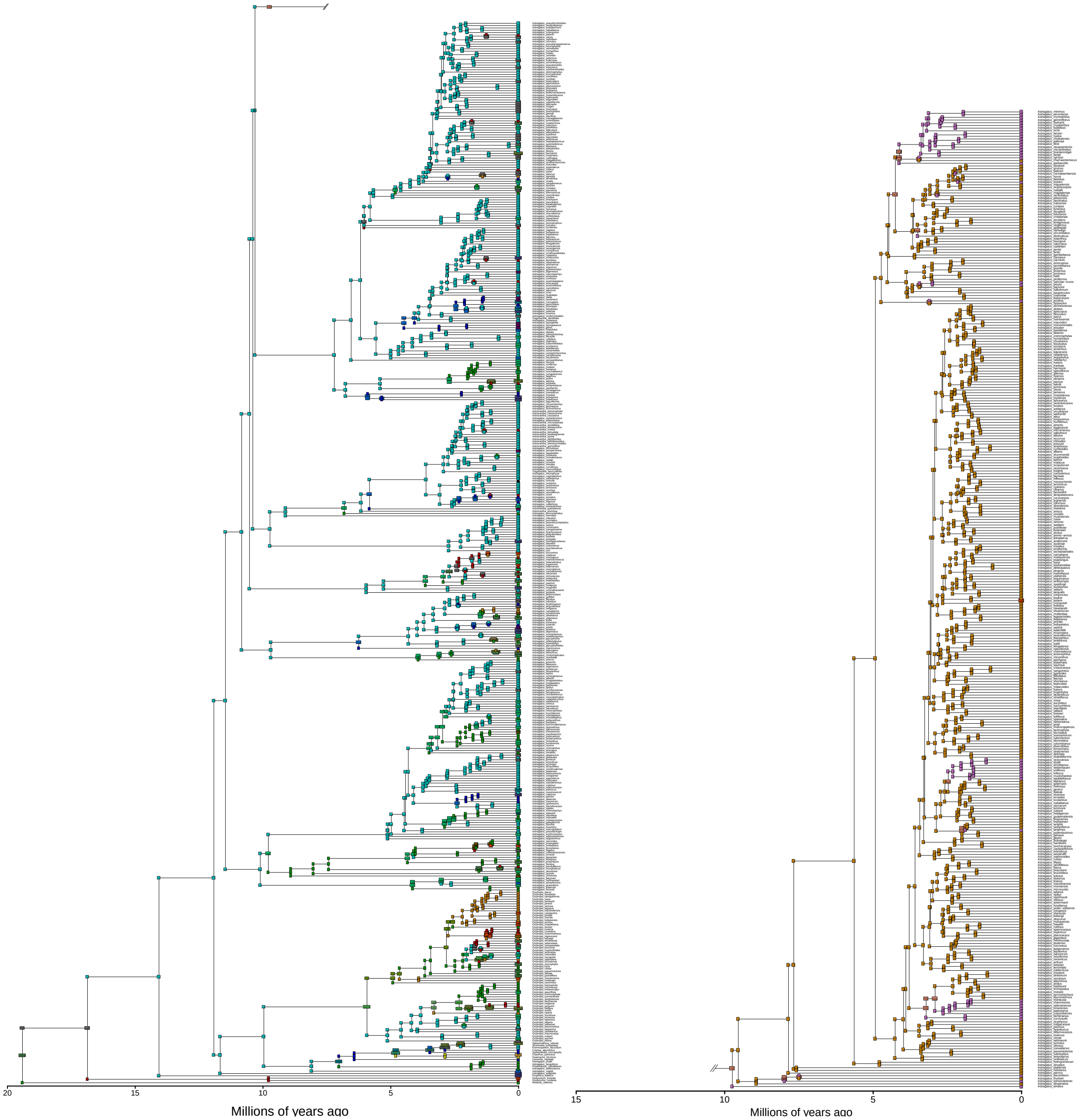

**Fig. S6.** Number of HybPiper paralog flags per sample, mapped across all samples on the pre-curation phylogeny. Warmer colors indicate greater numbers of paralogs.

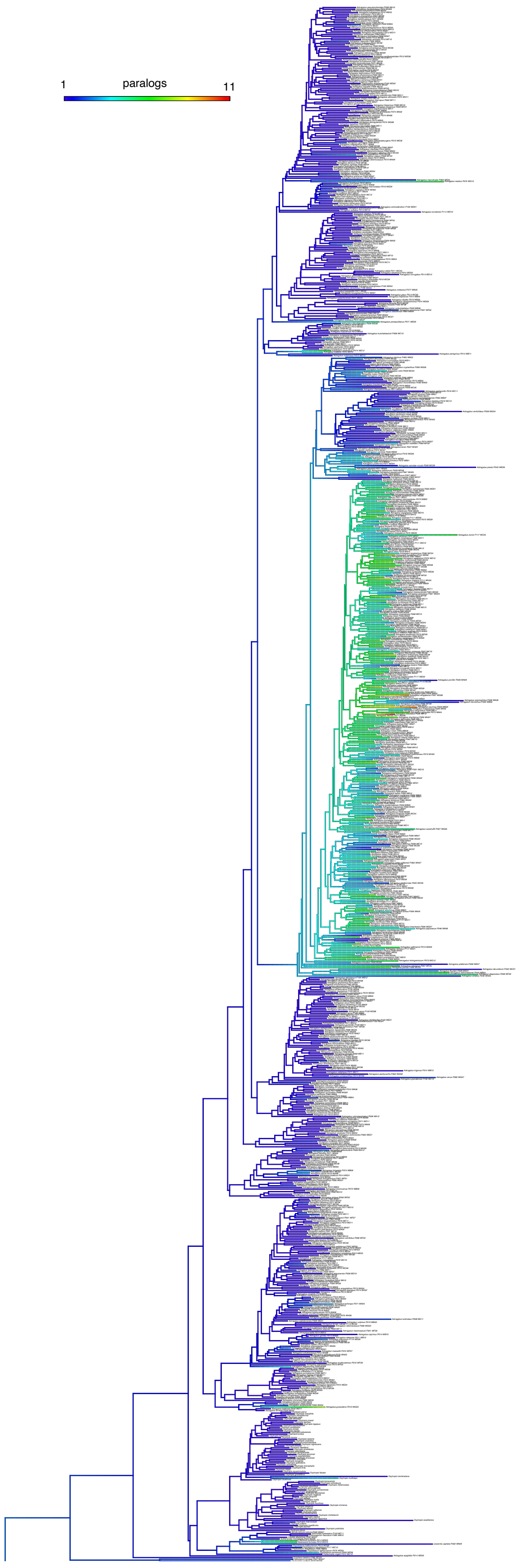
1
paralogs
11

**Fig. S7.** Number of loci assembled to at least 50% of the reference, mapped across all samples on the pre-curation phylogeny. Warmer colors indicate more loci at 50%.

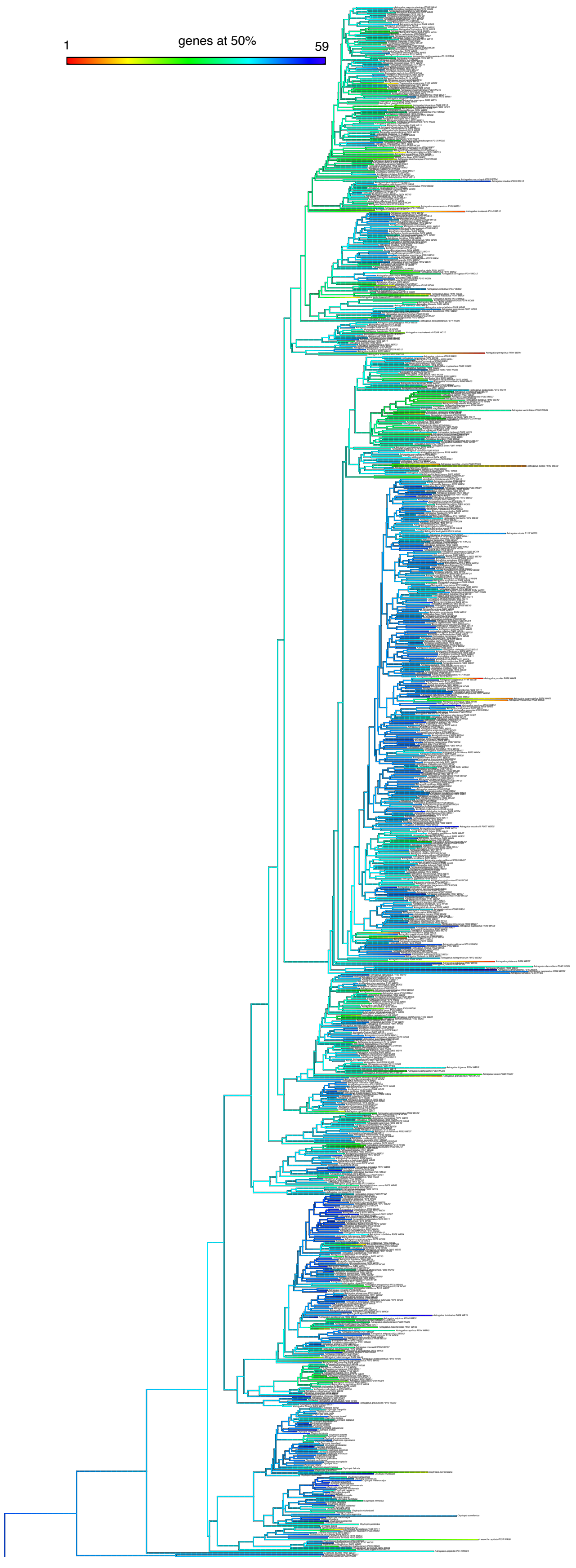
1
genes at 50%
59

| Species (Segregate) | Collector | Collector number | Herbarium | Locality | Year | NitFix UUID |
|---|---|---|---|---|---|---|
| ***Astragalus acanthochristianopsis*** | Podlech | 29236 | HUH | Afghanistan: Herat | 1977 | fd854b03-7a74-4425-bb5f-8c0b1d91807e |
| ***Astragalus acaulis*** | Woo | 425 | HUH | People's Republic of China: Yunnan: De Qin | 2002 | fbafb346-8cba-4158-82c0-caa92154f1f5 |
| ***Astragalus accidens*** | Spellenberg | 2296 | TEX | United States of America: California: Trinity | 1970 | 21984a3d-fb2f-4b43-b8d2-f1ac34079727 |
| ***Astragalus ackermanii*** | Ackerman | 30139 | NY | United States of America: Nevada: Clark | 1979 | d6350763-c09f-49e6-a87b-3e6c78ff5c73 |
| ***Astragalus acutirostris*** | Gentilcore | 886 | NY | United States of America: Nevada: Clark | 2016 | d633166e-5c43-4266-9e13-f8e3e218bde7 |
| ***Astragalus adanus*** | Ertter | 21340 | NY | United States of America: Idaho: Ada | 2013 | df6b4951-13a7-4828-b45e-55eb65408ca1 |
| ***Astragalus adpressipilosus*** | Vvedensky | 6201 | NY | Uzbekistan: Surxondaryo | 1929 | d021d9ad-f028-44c6-8094-0ca7848975ac |
| ***Astragalus aduncus*** | [Cyrillic] | | NY | United States of America: New York | 2018 | cf184ec2-cc82-4d32-ba20-79b7a2b72d61 |
| ***Astragalus adzharicus*** | Sytin | 7129 | NY | Georgia: Ajaria: Khulo | 1986 | d1c60a08-5415-4cc8-af7e-36a1fef22901 |
| ***Astragalus aegobromus*** | Litvinov | s.n. | NY | Turkey: Kars | 1914 | cf1587e1-b97b-4359-a76f-1b52a3b85a95 |
| ***Astragalus aegobromus*** | Litvinov | s.n. | MO | Turkey: Kars: Sarikamish | 1914 | 2ed50c28-a932-4b42-9a94-8fad7801a8ed |
| ***Astragalus aequalis*** | Johnson | 89 | NY | United States of America: Nevada: Clark | 1965 | df4877bf-5177-4e9d-b49c-701fc408754f |
| ***Astragalus afghanomontanus*** | Volk | 2735 | NY | Afghanistan: Bamian | 1952 | ad380c3c-08d6-4f6f-8940-bda2b7f09ea6 |
| ***Astragalus agrestis*** | Krivda | s.n. | FLAS | Canada: Saskatchewan: Choiceland | 1965 | e9da70d3-b54f-46b0-80c8-7e66be95e3d1 |
| ***Astragalus akkensis*** | Lewalle | 13669 | MO | Morocco | 1992 | 1580ade8-cd55-435c-a98a-a9deded28a32 |
| ***Astragalus aksuensis*** | Roldugin | 5376 | HUH | Kazakhstan: Alatau | 1968 | fb508295-95f7-4095-aff0-1f58bd5c44b4 |
| ***Astragalus albens*** | White | 92-154 | TEX | United States of America: California: San Bernardino | 1992 | 21aa2771-430b-442b-a50a-646990221be1 |
| ***Astragalus albertii*** | [None] | | NY | [None] | [None] | cf0b4084-d85c-43f7-a69e-494d644d2fd1 |
| ***Astragalus albicaulis*** | Sagalaev | 19.05 | NY | Russian Federation: Volgograd: Volgograd | 1979 | d1bcc2a6-d807-4df1-8ace-5d93acab50dd |
| ***Astragalus albulus*** | Atwood | 23126 | NY | United States of America: Arizona: Apache | 1997 | d625a1e6-1a02-41c3-9ed3-bc6922833fa6 |
| ***Astragalus algerianus*** | Dubuis | 13217 | MO | Algeria: M'Sila: Bou-Saada | 1986 | 1641c922-767a-4190-8060-9e5f3dc2f380 |
| ***Astragalus alitschuri*** | Gusev | 4583 | HUH | Tajikistan | 1958 | fb00fbee-70d4-4e4f-ab32-a4fd4d3e2417 |
| ***Astragalus allochrous*** | NA | NA | NA | Mexico: Sonora: Agua Prieta | 2003 | 16410fac-8fa7-481d-be8c-e633120a6272 |
| ***Astragalus alopecias*** | Dewey | D-2012 | NY | Kazakhstan | 1977 | cf0a0bb4-c1c7-4719-9ba7-e3b2b192eafa |
| ***Astragalus alopecuroides*** | Jahandiez | 180 | MO | Morocco | 1925 | 15828117-a8ff-43ac-a314-3f6642f30230 |
| ***Astragalus alopecurus*** | Alianskaja | 19.03 | HUH | Russian Federation: Gorno-Altajsk: Onghudaj | 1972 | faf3e866-5f85-4ca1-a889-394136d0d4ff |

| | | | | | | |
|---|---|---|---|---|---|---|
| ***Astragalus alpinus*** | Smith | 1914 | NY | United States of America: Alaska: Fairbanks North Star | 1953 | df46fd23-494a-4f77-a413-9744c245340d |
| ***Astragalus altimurensis*** | Volk | 71/430a | NY | Afghanistan: Paktia: Gardez | 1971 | ad301a32-de96-4d0a-95c2-f25ddc8fd5d3 |
| ***Astragalus altus*** | Spellenberg | 10470 | NY | United States of America: New Mexico: Otero | 1990 | d60ba04d-ab23-4d23-b57b-6b2bf2e2a420 |
| ***Astragalus alvordensis*** | Halse | 6954 | NY | United States of America: Oregon: Harney | 2006 | d605eb4d-c69c-43aa-8cb5-d46ac4866d6f |
| ***Astragalus alyssoides*** | Sytin | 7136 | NY | Azerbaijan: Talysh: Zuvant | 1985 | ad449517-da39-41f0-982e-244cb7dec8c2 |
| ***Astragalus amatus*** | Jiles | 1732 | OS | ChileLimar√≠ | 1950 | e4bcd0e3-fe65-4578-a354-af5324dc7c68 |
| ***Astragalus amblytropis*** | Rosentreter | 4366 | NY | United States of America: Idaho: Lemhi | 1987 | d602e66b-30ae-471d-987d-23df59e5c3dc |
| ***Astragalus americanus*** | Zobel | 112 | FLAS | United States of America: Colorado | 1939 | e9d8bf16-16da-49ff-bac9-6e91de15af95 |
| ***Astragalus amherstianus*** | Shah | 1937 | HUH | Pakistan | 1977 | fd838155-5c15-49bd-a632-e5b8a5a9da66 |
| ***Astragalus ammodendron*** | Androssv | 3301 | NY | Kazakhstan: Unknown: Unknown | 1908 | cf08fca8-7165-442a-ba2c-05d35d19567b |
| ***Astragalus ammodendron*** | Kurbanov | 678 | MO | Turkmenistan: Balkhan | 2001 | 0c8e8aee-4d5a-4c7f-9d6e-c10e3071e475 |
| ***Astragalus ammodytes*** | Veresczagin | 7422 | NY | Kazakhstan | 1925 | d1bb3e7f-1a2a-47aa-865d-d23c491024dc |
| ***Astragalus ammotrophus*** | [Cyrillic] | 63 | NY | Uzbekistan | 1968 | d013fbae-a544-48ce-96c7-333e38e60428 |
| ***Astragalus amnis-amissi*** | Duane Atwood | 10295 | NY | United States of America: Idaho: Custer | 1984 | df3be8be-7267-4d01-9c40-019369b42b3d |
| ***Astragalus amphioxys*** | Atwood | 25392 | NY | United States of America: Arizona: Navajo | 2000 | df3586ab-9e00-43d4-b60f-16dd2d9aea44 |
| ***Astragalus ampullarius*** | Neese | 16818 | NY | United States of America: Utah: Kane | 1985 | cc042c24-a746-44e0-bad7-f6814dd7c813 |
| ***Astragalus amygdalinus*** | Popov | 351 | NY | Uzbekistan | 1926 | d00f73bb-ef5e-44fc-a11d-763dd0b123f6 |
| ***Astragalus andersonii*** | Cronquist | 8453 | NY | United States of America: California: Lassen | 1959 | cb2fd040-c763-4a97-9d0f-9feb83940e97 |
| ***Astragalus anemophilus*** | Morita | 503 | CAS | Mexico: Baja California: Ensenada | 1999 | 163d2ede-6ecc-4449-8205-bbb111ba56ea |
| ***Astragalus angarensis*** | [Cyrillic] | 3302a | MO | Russian Federation: Irkutsk: Balagansky | 1907 | 0c8e29f4-15f1-4b7f-a131-cc2d0f29496a |
| ***Astragalus angustifolius*** | Donmez | 3747 | NY | Turkey | 1993 | ac0b563c-83b5-40ec-8905-d27776c90dbe |
| ***Astragalus angustissimus*** | Goloskokov | s.n. | NY | [Cyrillic] | 1963 | d009e8a4-f51b-4b21-90b2-9a27f86ab354 |
| ***Astragalus anisus*** | Taylor | 5308 | NY | United States of America: Colorado: Gunnison | 1998 | d600ddf7-5af3-4325-bf8d-c36e8b108493 |
| ***Astragalus annularis*** | Wojciechowski | 159 | NY | United States of America: Arizona | 1992 | ac182634-e4dd-4cd6-a321-4e14bfd2d201 |

| | | | | | | |
|---|---|---|---|---|---|---|
| ***Astragalus anserinaefolius*** | Ledingham | 4074 | NY | Islamic Republic of Iran | 1965 | ac1b55e8-93a3-4082-b1bb-9560fc36b9c0 |
| ***Astragalus anserinus*** | Hardy | 832 | NY | United States of America: Utah: Box Elder | 1993 | d6013bed-4ac0-4c7f-80f4-0c616e36c370 |
| ***Astragalus applegatii*** | Kagan | 6068503 | NY | United States of America: Oregon: Klamath | 1985 | d5f642e9-95d2-4fdd-a85e-757329e599b8 |
| ***Astragalus apricus*** | Ledingham | 4092 | NY | Islamic Republic of Iran | 1965 | ac7e4a39-96bf-427a-a0a1-44f7bb17c7f3 |
| ***Astragalus aquilonius*** | Reveal | 4483 | NY | United States of America: Idaho: Lemhi | 1976 | cb2fa16b-1053-4add-894f-33321630df7e |
| ***Astragalus arenarius*** | Asplund | 1076 | NY | Sweden: Skane: Simrishamn | 1928 | d1b0c116-8659-47e1-ad16-de0f09fcc160 |
| ***Astragalus arequipensis*** | Ricardi | | NA | Chile: Antofagasta (II): El Loa | 1961 | e4b6eb72-3464-43af-a7d6-6769760565d9 |
| ***Astragalus arganaticus*** | Goloskokov | 4274 | NY | Kazakhstan | 1956 | cfff2200-3b09-429a-98e7-0184230de39a |
| ***Astragalus argophyllus*** | Massatti | 4830 | NY | United States of America: Wyoming: Fremont | 2006 | cb2b7653-9193-41c3-a9c3-67ae1242f480 |
| ***Astragalus arguricus*** | Sytin | 6528 | NY | Georgia: Samtskhe-Javakheti | 1981 | d1a438d3-8eb3-48c3-8a8b-0f08cd37e1f1 |
| ***Astragalus argyroides*** | USDA-ARS | 314143 | NY | Uzbekistan | 1977 | cffe79f8-a553-4fe8-9072-3574e516091c |
| ***Astragalus aridus*** | Wiggins | 15743 | CAS | Mexico: Baja California: Mexicali | 1960 | 163bf792-235d-40f6-8459-75c032239ed0 |
| ***Astragalus arizonicus*** | Barneby | 18096 | NY | United States of America: Arizona: Yavapai | 1986 | cb27a811-dd3a-48de-a9fd-60f67dd44e4d |
| ***Astragalus armatus*** | Merello | 2842 | MO | Morocco | 2004 | 15800fae-4d35-4eee-bec4-cd5e93ed3b3b |
| ***Astragalus arnottianus*** | Landrum | 8225 | NY | Chile: Santiago Metropolitan | 1993 | d258cec1-c286-4fb2-ae6f-c95f3984c718 |
| ***Astragalus arpilobus*** | Rechinger | 57402 | NY | Islamic Republic of Iran: E. Khorasan: Unknown | 1975 | ab793654-64dd-47cb-8b1b-b1edfff8ae78 |
| ***Astragalus arthurii*** | Hufford | 894 | NY | United States of America: Washington: Asotin | 1995 | d5f1bdd6-2674-44a3-b33d-8d334fbdc2af |
| ***Astragalus asclepiadoides*** | Alexander | 1164 | NY | United States of America: Utah: Emery | 2001 | cb2aa313-5852-47ad-a71b-fad79f4bd63f |
| ***Astragalus askius*** | Ledingham | 4181 | NY | Islamic Republic of Iran: Mazandaran | 1965 | ac793212-7577-4367-b1aa-fd500a8c00ae |
| ***Astragalus asper*** | Barta | s.n. | NY | Austria: Vienna | 2008 | d19a605a-2e0f-4ae6-8d44-38eae2634203 |
| ***Astragalus asterias*** | Lachashvili | 390 | NY | GeorgiaDedoplistskaro | 2006 | d19610e3-d875-49dc-ad85-4803869af55b |
| ***Astragalus asymmetricus*** | Bedell | 97-1 | NY | United States of America: California: San Luis Obispo | 1964 | cb27a7ac-f323-4720-a431-7241f2b3a5ae |
| ***Astragalus atratus*** | Pinzl | 9449 | NY | United States of America: Nevada: Lander | 1991 | d5edfa4e-fbdc-474c-8a3a-6238ddb5c0a8 |

| | | | | | | |
|---|---|---|---|---|---|---|
| ***Astragalus atropilosulus*** | Rwaburindore | 4779 | NA | Uganda: Buhweju | 2000 | e5b14f77-4e08-4331-8eeb-a8d2f63e5992 |
| ***Astragalus atropubescens*** | Gray | 5582 | NY | United States of America: Idaho: Idaho | 2009 | d5ed5b26-0143-45e3-a734-9a4d2f49437e |
| ***Astragalus aureus*** | Ledingham | 5241 | NY | Armenia | 1970 | ac78aecf-6900-4857-b8e5-134362e328ff |
| ***Astragalus austinae*** | Genz | 8526 | NY | United States of America: Nevada: Washoe | 1978 | cb1211a4-609c-4b4a-8299-0ebf39e1ac5f |
| ***Astragalus australis*** | Wetherell | 784 | NY | United States of America: Alaska: Northwest Arctic | 1963 | df30e7e0-7567-479d-81d3-8aec0f9d07ec |
| ***Astragalus austriacus*** | Barta | s.n. | NY | Austria: Niederosterreich | 2009 | d219b02e-83f0-44fc-867f-15e701167812 |
| ***Astragalus austriacus (Astragalus tenuifolius)*** | Sagalaev | s.n. | NY | Russian Federation: Saratov | 1993 | cf984c50-2aef-4206-afb1-1d5b88932378 |
| ***Astragalus austroaegaeus*** | Rechinger | 7496 | NY | Greece | 1935 | d211012b-159e-4f43-868f-99899afe0852 |
| ***Astragalus austrosibiricus*** | Pobedimova | 445 | HUH | Mongolia | 1930 | 0093e40a-6469-4ae2-b897-c327e3f21ceb |
| ***Astragalus babatagii*** | [Cyrillic] | 63 | HUH | Russian Federation | 1959 | 0002cfe7-61fb-4339-974c-36e45fe75969 |
| ***Astragalus babatagii*** | Podlech | 30220 | HUH | Afghanistan: Takhar | 1977 | fd33dff3-3d7f-4d52-991b-9b2bd06597be |
| ***Astragalus baghlanensis*** | Podlech | 30326 b | NY | Afghanistan: Baghlan | 1977 | abf93cc7-8c50-42cf-a06d-196e206c3c74 |
| ***Astragalus baissunensis*** | Vvedensky | 6204 | NY | Uzbekistan: Surxondaryo | 1930 | ceed7a58-ab17-4527-a5b1-8695bf05ea0d |
| ***Astragalus bakaliensis*** | Volk | 1855b | NY | Afghanistan: Kabul | 1959 | abf3f532-55e2-44b1-9b4f-ef7812164eca |
| ***Astragalus bakuensis*** | [Cyrillic] | s.n. | NY | Azerbajdzhan: Transcaucasia | 1933 | cef3b32c-7bf0-448e-b478-dc7c330ee12d |
| ***Astragalus baldshuanicus*** | Vvedensky | 6205 | NY | Uzbekistan | 1929 | ceed240f-9a49-4f18-9447-7b918518199e |
| ***Astragalus baranovii*** | Baranov | 378 | NY | Uzbekistan: Tian Shan: Chimgan | 1926 | ced7732d-130d-49bb-bd28-0a0af564c012 |
| ***Astragalus baxoiensis*** | Boufford | 39460 | HUH | People’s Republic of China: Qinghai | 2007 | ffdefd8c-d9dd-4cf0-88db-4e5674d2e845 |
| ***Astragalus beckwithii*** | Holmgren | 16444 | NY | United States of America: Utah: Box Elder | 2013 | d5e88c4b-14c3-4c10-8dff-43b44664c731 |
| ***Astragalus bellus*** | McVaugh | 24485 | CAS | Mexico: Jalisco | 1970 | 15effd34-c08e-4583-bea8-241b948af8ed |
| ***Astragalus bergii*** | Hunziker | 17706 | NY | Argentina: Cordoba: Punilla | 1964 | d24aa630-4f65-4db6-8006-2a3bd7e3954b |
| ***Astragalus berteri*** | Zoellner | 4357 | NY | Chile | 1970 | d24a88e0-2129-4748-bdf2-596ca8188453 |
| ***Astragalus berterianus*** | Gunckel | 29710 | OS | Chile: Santiago: Polpaico | 1954 | e4b86421-be62-4543-b31f-00911f891bd2 |
| ***Astragalus bethlehemiticus*** | Amdursky | 539 | MO | Israel | 1954 | 0c848f4c-5732-444c-9bda-d149d526e300 |
| ***Astragalus bibullatus*** | Orzell | 9386 | TEX | United States of America: Tennessee: Rutherford | 1989 | 216819b8-9122-4d4d-a2f0-4055c9ea4eed |

| | | | | | | |
|---|---|---|---|---|---|---|
| ***Astragalus bicristatus*** | Thorne | 42077 | NY | United States of America: California: Los Angeles | 1972 | d5e037f5-9d97-4403-ad8b-5cbde60e5b76 |
| ***Astragalus bidentatus*** | Camp | E-4509 | NY | Ecuador: Azuay | 1945 | d2481099-10b4-4e2c-941d-ea94069fd919 |
| ***Astragalus bisulcatus*** | Atwood | 13926 | FLAS | United States of America: Utah: Duchesne | 1990 | e9d0e1d5-9e3b-47a4-ad06-7b652068b2f1 |
| ***Astragalus bodinii*** | Cody | 34777 | NY | Canada: Yukon Territory | 1984 | df27da94-2e99-4197-aee5-189a6ecfeb55 |
| ***Astragalus boeticus*** | Lewalle | 12834 | NA | NA: NA: NA | NA | 155e46d5-9c97-465f-9461-c246d3ec42c8 |
| ***Astragalus bolanderi*** | Helmkamp | 15306 | NY | United States of America: California: Riverside | 2009 | d5e37dd9-cdfb-4151-a71c-438b509708a3 |
| ***Astragalus bornmuellerianus*** | Kaletkina | 29 | NY | Tajikistan | 1966 | ced042b8-ac33-42eb-8652-00e2e61a0773 |
| ***Astragalus bourgaeanus*** | Podlech | 43151 | MO | Morocco: Khenifra | NA | 153b9a7d-038d-46eb-ad11-8d6bdfe0b243 |
| ***Astragalus bourgovii*** | Taylor | 2348 | NY | Canada: British Columbia | 1958 | d5dd237d-bce4-4973-98b0-f1200e00a8bb |
| ***Astragalus brachycarpus*** | Lachashvili | 334 | NY | Georgia: Dedoplistskaro | 2006 | d18a2b27-7ed1-4ddc-9474-a5a92d847453 |
| ***Astragalus brachylobus*** | [Cyrillic] | 7130 | NY | Russian Federation | 1961 | d18628ed-9173-4674-a038-8bc33e7223e3 |
| ***Astragalus brachypus*** | Toeosrorof | s.n. | NY | [Cyrillic] | 1963 | cecf6c91-334b-4824-adc8-110680ba2cc9 |
| ***Astragalus brackenridgei*** | Boeke | 1137 | NY | Peru: Pasco: Pasco | 1977 | d246d886-be5e-405d-898d-563a5e8f117c |
| ***Astragalus brandegei*** | Neesw | 16874 | NY | United States of America: Utah: Wayne | 1985 | cc07b4e0-7501-43a9-b71f-27b0a876dd17 |
| ***Astragalus brauntonii*** | Wallace | s.n. | NY | United States of America: California: Los Angeles | 1980 | d5c3d957-992c-48eb-a8ed-91dd341b2062 |
| ***Astragalus brazoensis*** | Lieb | 296 | NY | United States of America: Texas: Wilson | 1983 | d5c1f64e-ad25-4c71-b1eb-030c0d846236 |
| ***Astragalus brevifolius*** | [Cyrillic] | 2621 | HUH | Russian Federation: Tuva: Mongun-Tayginsky | 1981 | ff10b6a8-3d4f-49dc-bd15-6c187182ac86 |
| ***Astragalus brevipes*** | Sytin | 6531 | NY | Azerbaijan: Nakhichevan | 1982 | cfeebfcb-b0df-4180-9d9c-69d5df605a92 |
| ***Astragalus breweri*** | Liston | 752-1 | NY | United States of America: California: Lake | 1988 | d5bbbc77-1fba-4991-b1c9-9b0928cda52d |
| ***Astragalus bucharicus*** | [Cyrillic] | 87 | HUH | Russian Federation | 1959 | ff10b060-16bb-402f-b143-3dc525bbc4ab |
| ***Astragalus buchtormensis*** | Bespalova | 7047 | NY | Kazakhstan: Karaganda | 1959 | cec83870-6dcc-4f03-9871-a2a7ca153018 |
| ***Astragalus bungeanus*** | Lachashvili | 892 | NY | Georgia: Kvemo Kartli: Gardabani | 2007 | d17fbc4f-0cb7-4fb7-9ab0-1cab3ce524ee |
| ***Astragalus burkartii*** | Ledingham | 4495 | NY | Argentina | 1966 | d16fe1bc-3e16-4722-975d-44c7e9193cbb |

| | | | | | | |
|---|---|---|---|---|---|---|
| ***Astragalus bustillosii*** | Jiles | | NA | Chile: Coquimbo (IV): Limar√≠ | 1958 | e4b6bc69-e24c-49ab-b7e3-d0e6ebc51176 |
| ***Astragalus californicus*** | Barneby | 11510 | NY | United States of America: California: Siskiyou | 1954 | d5b4fa64-a033-4f58-815b-f57530d7649b |
| ***Astragalus calliphysa*** | Edmondson | No. 1636 | NY | Islamic Republic of Iran: Kerman | 1977 | ac75a99b-4eb7-4dbe-b356-9c3b9449aeff |
| ***Astragalus callistachys*** | Ledingham | 4152 | NY | Islamic Republic of Iran: Esfahan | 1965 | ac6aa448-4435-433e-947f-3711a737602f |
| ***Astragalus callithrix*** | Neese | 8784 | NY | United States of America: Nevada: Nye | 1980 | cce88927-e90d-4e7d-ba14-890f74b39e27 |
| ***Astragalus calycinus*** | Tzvelev | 7131 | NY | Russian Federation: Dagestan: Levashinskiy | 1961 | d1806ae7-ea57-4ff8-932f-dee931092d5d |
| ***Astragalus calycosus*** | Maguire | 20,840 | NY | United States of America: Nevada: White Pine | 1941 | ccdb86c3-8165-4620-a0a2-0bc5c67a6ef3 |
| ***Astragalus camptoceras*** | Wink C. & Fedtch P. | 5b | NY | Not shown: Not shown: Not shown | 1922 | cec73a6b-4e61-45e1-b1a8-ae335315f59b |
| ***Astragalus camptopus*** | Smith | 12581 | NY | United States of America: Idaho: Owyhee | 2015 | ccd4e380-42a1-46cd-beaf-869c672fd6a4 |
| ***Astragalus campylorhynchus*** | Grossheim | 14 | NY | Azerbaijan: Nakhichevan: Babek | 1923 | cce9c09d-4723-405e-b31c-f7f96c293992 |
| ***Astragalus campylorhynchus*** | Volk | 1650 | NY | Afghanistan: Vardak | 1951 | ac5a69f3-1e85-4a33-a588-25e44848d54f |
| ***Astragalus campylosema*** | USDA-ARS | 383603 | NY | Turkey: Tunceli | 1977 | ac25b3ef-1aaf-4d34-85e0-d8b5d217e301 |
| ***Astragalus campylotrichus*** | Goloskokov | 4466 | NY | Kazakhstan | 1963 | cceeffd4-dcb5-4920-9af3-abb658cc35f7 |
| ***Astragalus canadensis*** | Ward | s.n. | FLAS | United States of America: Indiana: White | 1945 | e9cf4a9f-de21-4fc3-bd05-b2e30f48ba80 |
| ***Astragalus cancellatus*** | Cocrum | s.n. | NY | [Cyrillic] | 1948 | d17d286c-a7bc-493e-bb1c-f2a85d56665b |
| ***Astragalus caprinus*** | Boulos | 2609 | MO | Tunisia | 1968 | 1533736e-a9f4-4298-af3b-0dfee55816c0 |
| ***Astragalus captiosus*** | Abdaladze | 240 | NY | Georgia: Mtskheta-Mtianeti: Kazbegi | 2004 | d17911ab-a8e7-46e1-9bd3-052569cee771 |
| ***Astragalus caraganae*** | [Cyrillic] | | NY | Armenia | 1954 | cffc22c0-330f-4578-9fda-19c07f3c66e3 |
| ***Astragalus caricinus*** | McKinnon | 379 | NY | United States of America: Washington: Franklin | 1993 | d5b3206a-ffc7-45c6-a209-f8eecbe12e7c |
| ***Astragalus carminis*** | Johnston | 1137 | NA | Mexico | 1940 | 157963a8-0a9b-4788-b066-2d85c3268ff5 |
| ***Astragalus casei*** | Arnold Tiehm | 9369 | TEX | United States of America: Nevada: Esmeralda | 1985 | 25cf0444-abd9-47ec-96e5-57e2c64228e9 |
| ***Astragalus castaneiformis*** | Ledingham | 3230 | NY | United States of America: Arizona | 1962 | ccca8723-26d1-4b3b-9a34-f9ab5ef315fe |
| ***Astragalus castetteri*** | Todsen | s.n. | NY | United States of America: New Mexico: Dona Ana | 1975 | ccbdf821-e8f5-4532-8acb-ba475a4cbe29 |
| ***Astragalus catabostrychos*** | Podlech | 31007 | NY | Afghanistan: Balkh | 1978 | abe9375c-b48d-4d8d-b171-89f5d4808856 |

| | | | | | | |
|---|---|---|---|---|---|---|
| ***Astragalus caucasicus*** | Iashagashvili | 43 | MO | Georgia: East Georgia | 2002 | 0c7c3116-b0e9-47be-b51e-a3d37f35e226 |
| ***Astragalus centralis*** | Botscgantzev | 6208 | HUH | Russian Federation | 1937 | 000a7723-eef3-4e7b-82bb-d3fd4e3127ac |
| ***Astragalus cephalanthus*** | Assadi | 1672 | NY | Islamic Republic of Iran | 1977 | abe4a123-aa5d-42fb-ac06-eeb9b62f9933 |
| ***Astragalus cephalotes*** | Litvinov | s.n. | MO | Turkey: Kars | 1914 | 0c61a0c6-210a-4c91-b1b7-47cd7fecf15a |
| ***Astragalus ceramicus*** | Atwood | 15436 | FLAS | United States of America: Utah: Beaver | 1991 | e9caf60a-10c3-42a1-9086-da53cb00aae3 |
| ***Astragalus cerasinus*** | O'Kane | 2502 | NY | United States of America: Colorado: Conejos | 1986 | d593f077-83d0-4690-a1b2-b2d7ab216727 |
| ***Astragalus cerasocrenus*** | Kurbanov | 306 | MO | Turkmenistan: Ahal | 2001 | 0c739979-ab4c-4198-98ca-5b1762d30efb |
| ***Astragalus chamaeleuce*** | Huber | 5384 | NY | United States of America: Utah: Duchesne | 2014 | d57e7f71-d363-4399-847d-5b666ce1c882 |
| ***Astragalus chamaemeniscus*** | Reveal | 2328 | NY | United States of America: Wyoming: Carbon | 1971 | ccb86491-1ed5-4a48-b737-0da9279c76e0 |
| ***Astragalus chamissonis*** | Marticorena | 969 | NA | Chile: Maule (VII) | 1967 | e4b4e4c7-0ea4-410f-9b1d-d1ebb0008aa0 |
| ***Astragalus chilienshanensis*** | Boufford | 35652 | HUH | People's Republic of China: Sichuan: Yajiang | 2006 | ffffe391-d600-4adc-9cb6-82fe307e550d |
| ***Astragalus chinensis*** | Ledingham | 4815 | NY | Canada: Saskatchewan | 1966 | d69fcd47-70ed-4e11-afac-42a70d9ac6ff |
| ***Astragalus chloodes*** | Goodrich | 27501 | NY | United States of America: Utah: Uintah | 2009 | d697a3b7-d2f1-4095-b863-9c6f40edba0a |
| ***Astragalus chlorostachys*** | Barneby | s.n. | NY | Czeckoslovakia: Pruhonice: [cultivated] | 1972 | aa3c12e0-adf5-48b8-956a-b98532fa76a4 |
| ***Astragalus chorizanthus*** | Volk | 1716 | NY | Afghanistan: Kabul | 1951 | ce214c4f-6e3a-4858-8b56-8fdab9dfa16f |
| ***Astragalus chrysomallus*** | Butkov | 6258 | HUH | Uzbekistan | 1931 | fff5249e-723e-4a52-8158-5e3b96b10f4d |
| ***Astragalus chrysostachys*** | Lamond | 4466 | NY | Islamic Republic of Iran: Kordestan: Sanandaj | 1971 | cff72ae3-ea8f-4233-9e84-75a555e809cb |
| ***Astragalus chubutensis*** | Donat | 178 | NY | Argentina: Santa Cruz: Caleta Olivia | 1929 | d1635734-62b4-4a5f-b9c4-52cdf46e99b6 |
| ***Astragalus chuskanus*** | Rink | 11409 | NY | United States of America: Arizona: Apache | 2012 | ccb170f5-227e-455d-bcfa-92e19e1989ac |
| ***Astragalus cibarius*** | Lesica | 5359 | NY | United States of America: Montana: Carbon | 1991 | ccad2b2c-31ac-4cc6-b852-b15479ebb9a1 |
| ***Astragalus cicer*** | Ashley N. | 15-77 | NY | United States of America: Utah: Sanpete | 1915 | d694a0c5-02fc-454c-b2cb-fde3e6022818 |
| ***Astragalus cimae*** | Holmgren | 8167 | NY | United States of America: Nevada: Nye | 1976 | d68d35fa-9e03-4dc0-9e32-2ab90963add3 |
| ***Astragalus circumdatus*** | Wiggins | 21461 | CAS | Mexico: Baja California | 1971 | 1633b009-2bf5-4327-926d-15327eda5aa8 |

| | | | | | | |
|---|---|---|---|---|---|---|
| ***Astragalus citrinus*** | [Cyrillic] | 6203 | HUH | Turkmenistan | 1931 | 0015709b-0007-4836-81cb-1b1faf84f309 |
| ***Astragalus clarianus*** | Barneby | 11238 | NY | United States of America: California: Sonoma | 1954 | d683f828-e577-4386-bf67-ef0b4a0f2020 |
| ***Astragalus clevelandii*** | Spellenberg | 3739 | NY | United States of America: California: Napa | 1974 | d67c3599-4fb5-4d43-bd1f-0579665b3eac |
| ***Astragalus clusii*** | Borja | s.n. | NY | Spain: Toledo | 1965 | d20c8c0b-b095-43fb-a89a-134101cbbca0 |
| ***Astragalus coahuilae*** | Ripley | 13513 | CAS | Mexico: Coahuila | 1963 | 1572a2de-80b1-4198-82c6-28698d0e126f |
| ***Astragalus cobrensis*** | Keil | 4475 | NY | United States of America: Arizona: Gila | 1969 | cc996174-ba96-4dcd-b097-3465f2fea300 |
| ***Astragalus coccineus*** | Walker | 1414 | NY | United States of America: California: Riverside | 1995 | cc98799e-d99f-4b64-ae1f-a0037d91760d |
| ***Astragalus cochabambensis*** | Dillon | 5060 | NY | Chile: Antofagasta: Antofagasta | 1987 | d29e4ec9-2471-4b38-ad5e-c0bc16e5b9b0 |
| ***Astragalus cognatus*** | Goloskokov | 5770 | NY | Kazakhstan | 1971 | cff36c85-e87a-4a03-8106-44c7649becde |
| ***Astragalus collinus*** | Gray | 5357 | NY | United States of America: Idaho: Nez Perce | 2006 | cc94a1bb-8e04-46e1-9fe5-9c47bbf3835d |
| ***Astragalus coltonii*** | O'Kane | 2348 | NY | United States of America: Colorado: Montezuma | 1986 | cc9475d8-02c6-4d59-b4eb-a8fe6aa6e442 |
| ***Astragalus columbianus*** | Joyal | 480 | NY | United States of America: Washington: Benton | 1984 | d67913a6-3e4b-43f5-a10b-a80a1625aa23 |
| ***Astragalus coluteocarpus*** | [Cyrillic] | 1158 | HUH | [Cyrillic] | 1965 | ffea8400-5a7a-42ca-b79c-24f8288da752 |
| ***Astragalus comosus*** | USDA-ARS | 384742 | NY | Islamic Republic of Iran | 1977 | aa314714-9dc5-45ba-8e1b-8f2af46c5c31 |
| ***Astragalus confertus*** | Ho | 1419 | HUH | People’s Republic of China: Qinghai | 1993 | ffd4d3ff-31a8-4acd-a374-fb87e1e29c09 |
| ***Astragalus congdonii*** | Howell | 40516 | NY | United States of America: California: Mariposa | 1964 | d66c1e4f-956a-445d-aae1-1566272def68 |
| ***Astragalus conjunctus*** | Halse | 6350 | NY | United States of America: Oregon: Wheeler | 2003 | cc8dba0a-47aa-4d0b-bc6b-a95da5c81c88 |
| ***Astragalus consanguineus*** | Schrenk | s.n. | NY | Kazakhstan [Songarei] | [None] | d20a6a6d-3701-4ac3-bc3e-3d0997a1aa96 |
| ***Astragalus consobrinus*** | Franklin | 6141 | NY | United States of America: Utah: Wayne | 1988 | d6657434-04b8-4bbf-a8e0-36d8bbb1373b |
| ***Astragalus contortuplicatus*** | Klinkova | s.n. | NY | Kazakhstan: West Kazakhstan: Bokey Orda | 1993 | d0ab6c7c-393e-4f97-898f-038103f3522e |
| ***Astragalus convallarius*** | Rogers | 652 | FLAS | United States of America: Washington: Ferry | 1940 | e9c56c49-ee98-452e-b246-54df6fc2d9b8 |
| ***Astragalus coquimbensis*** | Munoz | s.n. | NA | Chile: Atacama (III): Cha√±aral | 1991 | e4b125bd-2f45-4df3-9b1f-86098bc88562 |
| ***Astragalus cornutus*** | Gagnidze | 1026 | NY | Georgia | 2004 | d0bc3785-7b7c-460a-92e3-f1174e10c587 |

| | | | | | | |
|---|---|---|---|---|---|---|
| ***Astragalus corrugatus*** | Thomas | 2005 | NA | Algeria | 1983 | 153045b4-3607-47aa-ad62-f3c131afd643 |
| ***Astragalus cottonianus*** | Botschantzev | 6260 | HUH | Uzbekistan: Surxondaryo | 1930 | ff36fb0f-5ac1-4449-8b6c-df4545d78e1d |
| ***Astragalus cracca*** | Lopez M. | 7610 | NY | Peru: Recuay | 1970 | d29928c8-7289-4f94-8554-facdab47053a |
| ***Astragalus crassicarpus*** | Carter | 2171 | FLAS | United States of America: Iowa: Osceola | 1957 | e9cd683e-6cde-47c7-b868-368d7dfd7455 |
| ***Astragalus cremnophylax*** | Brian | 95-168 | NY | United States of America: Arizona: Coconino | 1995 | d66311da-acc1-45ec-91fc-d7d618ec0f9f |
| ***Astragalus crenatus*** | Podlech | 50839 | MO | Morocco: Er Rachidia | 1993 | 1532412e-cc64-4caf-9d2a-1e44df1f29d1 |
| ***Astragalus cretaceus*** | Samuelsson | 2714 | NY | Palestine, State of | 1933 | aa294139-4d6e-4493-903c-26ff06dc6881 |
| ***Astragalus creticus*** | Lambert | 591 | NY | Turkey | 1968 | aa2a478e-1dc3-407a-9d28-e163741bdbdf |
| ***Astragalus cronquistii*** | Barneby | 17803 | NY | United States of America: Colorado | 1982 | cc8c0252-4bac-4c6e-8b9e-e55d2a38542f |
| ***Astragalus crotalariae*** | Sphon | 254 | FLAS | United States of America: California: Imperial | 1959 | e9bb7aff-fcb6-4270-9e66-427685a1bc69 |
| ***Astragalus cruckshanksii*** | Kiesling | 8115 | OS | Argentina: San Juan: Calingasta | 1992 | e4a6cb79-f189-446a-9c36-5ca7673e24c7 |
| ***Astragalus crymophilus*** | Shapiro | | NY | Argentina | 1986 | d29096a7-77a9-4441-8773-9212d76b4f63 |
| ***Astragalus cryptanthus*** | Ricardi | 231 | NA | Chile | 1961 | e49285b5-4e7b-4a8a-98ec-b394008318e3 |
| ***Astragalus cryptobotrys*** | Ricardi | | NA | ChileParinacota | 1961 | e4904d58-d2b3-4a12-a31c-359574e1c25d |
| ***Astragalus curtipes*** | Cassen | 798 | FLAS | United States of America: California: San Bernardino | 1974 | e9b7ad8b-0e60-4e82-98e3-7ac37f0d8af0 |
| ***Astragalus curvicarpus*** | Isely | 10069 | NY | United States of America: Oregon: Deschutes | 1966 | d662c421-970c-4474-a9fe-b05dcf07f0f1 |
| ***Astragalus curvicaulis*** | Ellenberg | 4650 | NY | Chile: Valparaiso | 1971 | d2904729-d4fd-45f0-90a2-618b54731792 |
| ***Astragalus curviflorus*** | Ledingham | 4132 | NY | Islamic Republic of Iran | 1965 | abb94793-de88-4fc4-8bf3-b7007f151b39 |
| ***Astragalus curvirostris*** | USDA-ARS | 380710 | NY | Islamic Republic of Iran: Fars: Shiraz | 1977 | abb0eb5e-a4f2-4643-9988-79da5f73b7e2 |
| ***Astragalus cusickii*** | Bala | 11-71a | NY | United States of America: Idaho: Adams | 2011 | cc8b5edb-192e-4efc-91b2-ff7e1fc4b81a |
| ***Astragalus cyaneus*** | Higgins | 5163 | NY | United States of America: New Mexico: Santa Fe | 1972 | cc84a95a-8da9-4c49-aa7a-c54dd5e4248f |
| ***Astragalus cymboides*** | Holmgren | 9107 | NY | United States of America: Utah: Emery | 1979 | cc8445ce-3121-4a43-a6b2-2079055182a5 |
| ***Astragalus cyri*** | Lachashvili | 31 | NY | Georgia: Kartli: Mtskheta | 2005 | d0bf5322-395b-401f-bb28-517145f74a9a |

| | | | | | | |
|---|---|---|---|---|---|---|
| ***Astragalus dactylocarpus (Astragalus spinescens)*** | [Cyrillic] | [Illegible] | NY | Turkmenistan | 1978 | acf192aa-4c52-4d61-b7a2-94243dfb438c |
| ***Astragalus dactylocarpus*** | Volk | 2906 | NY | Afghanistan: Helmand | 1953 | abab945f-c513-4903-95f0-5a584e8147a5 |
| ***Astragalus dahuricus*** | Hu SY | 19730 | HUH | Mongolia | 1984 | fadace43-65ab-4303-9c15-6ec9905ccf71 |
| ***Astragalus daleae*** | Breedlove | 63018 | NA | Mexico: Durango | 1986 | 15726290-bdda-43b1-b96b-aff11ddc8f87 |
| ***Astragalus danicus*** | Jacobsen | 278 | NY | Denmark: Zealand | 1970 | d0c92fe7-333a-47b0-b710-e4a0283b21bf |
| ***Astragalus darumbium*** | Jiles | 4107 | NA | Chile | 1962 | e4714581-cf1c-4886-ad98-d29857a61565 |
| ***Astragalus dasyanthus*** | Akinfiev | 7244 | NY | Ukraine: Dnipropetrovsk | 1906 | d0cda681-47a0-4cda-bd87-c000c6ca5d31 |
| ***Astragalus debequaeus*** | Nelson | 79077 | NY | United States of America: Colorado: Mesa | 2010 | d65f9c07-3457-47ed-9eef-c426a38aad6f |
| ***Astragalus densifolius*** | Orhan | 1186 | MO | Turkey: Ankara: Kazan | 1993 | 0c1317c2-1b5c-4dfc-87ec-1cd503202fef |
| ***Astragalus denudatus*** | Karjagin | s.n. | NY | Azerbaijan: Shabran | 1937 | b1012b6b-eb41-4d39-839b-5e9f32333820 |
| ***Astragalus depressus*** | Stephani | s.n. | NY | Italy: Abruzzo: L'Aquila | 1927 | d17561d2-fac7-4b5f-919e-b91ec9ebccfc |
| ***Astragalus desereticus*** | Neese | 10391 | NY | United States of America: Utah: Utah | 1981 | d65f8edf-a741-4baa-bfb0-e96217a8297d |
| ***Astragalus desperatus*** | Atwood | 15021 | FLAS | United States of America: Utah: Garfield | 1991 | e95ff422-ad88-4038-afeb-e8d0b22dee36 |
| ***Astragalus deterior*** | Welsh | 2131 | NY | United States of America: Colorado: Montezuma | 1963 | cc82d713-452a-486b-98bb-fcc6db9bbdd4 |
| ***Astragalus detritalis*** | Huber | 5067 | NY | United States of America: Utah: Duchesne | 2011 | cc7da259-102b-41a8-bb93-35eff828e4f0 |
| ***Astragalus dianthus*** | Vvedensky | 6 | NY | Uzbekistan | 1922 | ccf0cf4e-b85d-4be6-a31f-70dbd511682b |
| ***Astragalus diaphanus*** | Wright | 1686 | NY | United States of America: Oregon: Grant | 1983 | cc7ada49-42ac-4d67-8728-178e594d0eb9 |
| ***Astragalus dictyolobus*** | Lamond | 3642 | NY | Islamic Republic of Iran: Azerbaijan | 1971 | ccf3aa90-ecd6-44c5-9959-3ce2609131d2 |
| ***Astragalus didymocarpus*** | Schramm | 1784 | NY | United States of America: California: Inyo | 1978 | d655f50b-16a0-40d0-bb4f-3c9841b561ac |
| ***Astragalus dilutus*** | Schischkin | 3137 | HUH | Russian Federation: Oirotia: Altai | 1931 | fad05c65-c8af-4433-b58c-db6bcb3d80f0 |
| ***Astragalus diminutivus*** | Deginani | 520 | NY | Argentina: Jujuy: Cochinoca | 1995 | d28aa9f3-fc66-4a7e-b493-c7e9d2064a59 |
| ***Astragalus dipelta*** | Mokeeva | 383 | NY | Unknown Country | 1928 | cd0c526b-f998-46a3-a742-bac91e255458 |
| ***Astragalus diphacus*** | Yen | 15646 | NY | Mexico: Nuevo Le√≥n: Galeana | 2015 | d3f94d5d-52c3-44f8-9b55-2e8031f46d44 |
| ***Astragalus diphtherites*** | Rechinger | 11548 | MO | Iraq: Dihok: Dohuk | 1957 | 0c18db31-c02d-4f4b-8d29-f65f32a94717 |

| | | | | | | |
|---|---|---|---|---|---|---|
| ***Astragalus discolor*** | Hu | 19700 | HUH | Mongolia | 1984 | fab91016-5bab-41c4-8e95-1002e5874eed |
| ***Astragalus distentus*** | Burkart | 5661 | NY | Argentina: Buenos Aires: Campana | 1933 | d3f8097f-02e2-47cb-a023-9fc71da04622 |
| ***Astragalus distortus*** | Redfearn Jr | 9815 | FLAS | United States of America: Missouri: Greene | 1962 | e96cb642-09c5-475c-bb1a-703089a28fe7 |
| ***Astragalus diversifolius*** | Caicco | 385 | NY | United States of America: Idaho: Custer | 1982 | d64f0fc2-5331-4c74-aa13-4040ae17c3fe |
| ***Astragalus dodtii*** | Munoz | s.n. | OS | Chile: Atacama (III): Copiap√≥ | 1991 | e48f77ab-a79c-4ba4-9d1e-1b79ef6a56e8 |
| ***Astragalus dolichocarpus*** | Granitov | 355 | NY | Uzbekistan? | 1926 | cd1ee8aa-b6d7-4837-99f5-e1df20d7b5db |
| ***Astragalus dolichophyllus*** | Klinkova | s.n. | NY | Russian Federation: Astrakhan: Akhtubinsky | 1990 | d1faca8c-0e5f-42c7-9560-36ca65162fc8 |
| ***Astragalus douglasii*** | NA | NA | NA | Mexico: Baja California: Ensenada | 1961 | 1622ccca-0c25-4463-8e43-6e702877f846 |
| ***Astragalus drabelliformis*** | Holmgren | 8374 | NY | United States of America: Wyoming: Sublette | 1977 | cc72ef86-77cd-4c59-b771-34eacf55385c |
| ***Astragalus drummondii*** | Foster | 10430 | FLAS | United States of America: Wyoming: Carbon | 1991 | e9626080-c4ec-4d0c-b8eb-ea8e20074f76 |
| ***Astragalus duchesnensis*** | Goodrich | 27952 | NY | United States of America: Utah: Uintah | 2011 | cc6ed3a2-2cad-463f-919c-64ed13d33758 |
| ***Astragalus echanensis*** | Dieterle | 1245 | NY | Afghanistan: Bamyan | 1971 | aba4d335-9fcb-426e-bc83-fa478a676da8 |
| ***Astragalus ePeople's Republic of Chinatus*** | Podlech | 43613 | MO | Morocco: Kenitra | 1987 | 151c4c7f-0014-40d9-adb0-b2670a28b264 |
| ***Astragalus echinops*** | Assadi | 2102 | NY | Islamic Republic of Iran: Kerman: Jiroft | 1977 | aba10a32-c891-4fe2-8be2-8334b4544247 |
| ***Astragalus edmondstonei*** | Gunckel | 39476 | NA | Chile: Valparaiso (V): Valparaiso | 1956 | e485ab4d-9494-4d98-9fb1-16ba17d249c3 |
| ***Astragalus edulis*** | Davis | 48892 | NY | Morocco: Souss-Massa: Taroudannt | 1969 | d05bdb12-bc68-4b95-a215-d85e7c1efca0 |
| ***Astragalus egglestonii*** | Spellenberg | 10871 | NY | United States of America: New Mexico: Socorro | 1991 | d649e8ad-d76c-4369-827a-7c9514344688 |
| ***Astragalus ellipsoideus*** | Goloskokov | 4326 | NY | Kazakhstan | 1959 | cfd315c3-5b04-494d-9c88-46c472175fde |
| ***Astragalus elymaiticus*** | Ledingham | 4068 | NY | Islamic Republic of Iran: Fars: Shiraz | 1965 | aba34fe2-d097-4cc5-b619-89ebef3e6b89 |
| ***Astragalus emarginatus*** | Liston | 496/10 | MO | Israel: HaZafon: Golan | 1986 | 0c67de25-81e3-4d7e-b15a-b69a83f99a3c |
| ***Astragalus emoryanus*** | Johnston | 10207 | CAS | Mexico: Nuevo Le√≥n | 1973 | 161d0726-efd8-4fc4-b63d-6ea755ed4175 |
| ***Astragalus endopterus*** | Demaree | 44420 | NY | United States of America: Arizona: Coconino | 1961 | cc59189d-ee7b-471e-9416-416a7fcfdf01 |
| ***Astragalus ensiformis*** | Coombs | 2440 | NY | United States of America: Arizona: Mohave | 1978 | cc5662f3-ef6f-4c83-900b-062c9141dbd2 |

| | | | | | | |
|---|---|---|---|---|---|---|
| ***Astragalus episcopus*** | Hufford | 1843 | NY | United States of America: Utah: Emery | 1997 | d7403ed3-fed1-4e87-a9ca-d6f06d0cab1e |
| ***Astragalus episcopus (Astragalus lancearius)*** | Welsh | 28054 | NY | United States of America: Utah: Kane | 2001 | cb9992b2-9936-47c9-adbc-296d01cd8d58 |
| ***Astragalus eremiticus*** | Ben Franklin | 7029 | FLAS | United States of America: Utah: Washington | 1990 | e95531a2-bd0a-4dc8-a228-6051daea3101 |
| ***Astragalus eremophilus*** | Adam | 13280 | MO | Mauritania: Atar | 1957 | 157d0c99-f083-419b-952e-4a646d16b713 |
| ***Astragalus eriocarpus*** | Szoviks | s.n. | NY | Armenia | [None] | b0f66b2e-11c6-4132-972c-9c90c359f9fb |
| ***Astragalus eriopodus*** | [Cyrillic] | | NY | [Cyrillic] | 1956 | cfc9b8b4-3f22-477f-8d77-24a8372cbb92 |
| ***Astragalus ernestii*** | D.E. Boufford | 39960 | HUH | People's Republic of China: Sichuan | 2007 | fa758999-c557-4732-a0e9-b9dfe9b4ca1a |
| ***Astragalus ervoides*** | NA | NA | NA | Mexico: Sinaloa: Concordia | 1983 | 167d1234-4119-4e84-a737-85af125d4178 |
| ***Astragalus erythrosemius*** | Volk | 71/38 | HUH | Afghanistan: Paktia | 1971 | f80a227e-ed97-4a1c-abf6-da72ec3f12da |
| ***Astragalus eucosmus*** | Dutilly | 39,425 | NY | Canada: Quebec | 1961 | cc4f5cb7-86db-45a0-ba61-dffd44c43135 |
| ***Astragalus eurekensis*** | Welsh | 1970 | NY | United States of America: Utah: Wasatch | 1963 | d72ecbec-eb68-4f48-a367-035ddbc27ec3 |
| ***Astragalus eurylobus*** | Nichols | 434 | NY | United States of America: Nevada: Lincoln | 1985 | cc4ac8c1-b4d3-4b62-8c92-ed6c09fbbdb4 |
| ***Astragalus exscapus*** | Lewalle | 12381 | MO | Morocco | 1989 | 154ec09b-1aed-470f-96c3-72d9efdb86ad |
| ***Astragalus fabaceus*** | Lachashvili | 384 | NY | Georgia: Kakheti: Dedophlistkaro | 2006 | d1f7025a-c0e1-49bb-ad40-d0792490160b |
| ***Astragalus falcatus*** | Weber | 15359 | NY | United States of America: Colorado: Boulder | 1978 | d73c5799-8bd0-472e-ac68-72930aa6ac6a |
| ***Astragalus falconeri*** | Webster | 5929 | HUH | Pakistan: Baltistan | 1955 | f7f48647-7188-47f3-8ca0-1deeda4ce040 |
| ***Astragalus farctissimus*** | Butkov | 6212 | HUH | Uzbekistan | 1931 | fa60b678-e9fd-45db-a3d4-80821bf4ee5a |
| ***Astragalus fasciculifolius*** | Ledingham | 4123 | NY | Islamic Republic of Iran: Shiraz | 1966 | aa20932b-56f1-4180-8cf8-a08de705f68f |
| ***Astragalus fastidius*** | Breedlove | 72822 | CAS | Mexico: Baja California | 1993 | 16790db1-b51a-4bce-84b2-421419b8d634 |
| ***Astragalus fedtschenkoanus*** | Skvortsov | s.n. | HUH | Kazakhstan | 1965 | fa4f5d06-b2e0-42ac-8551-0a881246ff42 |
| ***Astragalus feensis*** | Barneby | 17949 | NY | United States of America: New Mexico: Santa Fe | 1983 | d71118e5-5642-4d3a-92b8-0d261f3bdf8a |
| ***Astragalus filicaulis (Astragalus rytilobus)*** | Goloskokov | 4472 | NY | Kazakhstan | 1937 | b0274a7e-f050-44be-9556-a9208a6e1877 |
| ***Astragalus filicaulis*** | Podlech | 30206 | NY | Afghanistan: Takhar: Farkhar | 1977 | aa19ecab-227f-4849-b777-f0801e464fd9 |

| | | | | | | |
|---|---|---|---|---|---|---|
| ***Astragalus filipes*** | Moran | 28914 | FLAS | Mexico: Baja California | 1980 | e94949f8-c697-4526-bcfc-6fdd4868a4c9 |
| ***Astragalus flavus*** | Foster | 10415 | FLAS | United States of America: Wyoming: Uinta | 1991 | e942cc23-dae2-45b5-8278-030909575238 |
| ***Astragalus flexilipes*** | Ledingham | 4261 | NY | Islamic Republic of Iran | 1965 | aa14b66b-91a5-4647-aa56-159db76c4437 |
| ***Astragalus flexuosus*** | Atwood | 29690 | NY | United States of America: Colorado: Chaffee | 2003 | cc44afdf-d8ca-46b0-867c-d91d7631ac80 |
| ***Astragalus floridus*** | Boufford | 29979 | NY | People's Republic of China: Xizang (Tibet) | 2000 | cea8a748-14f7-4238-8483-33f337ecf7e2 |
| ***Astragalus follicularis*** | Cronquist | 12119 | NY | Russian Federation: Gorno-Altaysk | 1988 | cd56e556-9ef2-49c3-94cc-9c9433da8dd9 |
| ***Astragalus forrestii*** | Boufford | 28682 | HUH | People's Republic of China: Sichuan: Garz√™ | 1998 | fa2ce01f-bd8c-457b-a5e5-fd6d3e369d2b |
| ***Astragalus fraxinifolius*** | USDA-ARS | 383604 | NY | Turkey: Dagyolo | 1977 | aa07629f-6fb9-4acf-ae4f-c098f1db3501 |
| ***Astragalus frickii*** | Atha | 5875 | NY | Georgia: Adigeni | 2007 | d1fe2345-e06e-4ae8-8556-4c72d18287e5 |
| ***Astragalus frigidus*** | Delannay | 1195 | MO | Switzerland: Graubunden: Albula | 1973 | 09c4b1db-d19c-42ff-b505-f9dd9c53a89e |
| ***Astragalus frigidus*** | Gage | SG 2426 | NY | Russian Federation | 1996 | d1ec075d-d37e-4651-afbe-aa74faa35ba4 |
| ***Astragalus frigidus (Astragalus secundus)*** | Webster | 6047 | HUH | India | 1955 | f7a96c9a-208b-44eb-9c63-0307df3960fe |
| ***Astragalus froedinii*** | Jahandiez | 235 | NY | Morocco: Grand Atlas | 1931 | d04be173-87e7-4518-a645-39a104775a94 |
| ***Astragalus fruticosus*** | Ikonnikov-Galitzky | 2416 | HUH | Mongolia | 1929 | feb58bbb-fe77-40ae-b385-1b183bf99794 |
| ***Astragalus fucatus*** | Atwood | 21743 | NY | United States of America: Utah: San Juan | 1997 | cc31aa5f-3b8d-4ae5-8a1c-67a5b69326d2 |
| ***Astragalus funereus*** | Cochrane | 372 | NY | United States of America: Nevada: Nye | 1977 | d72d6ea9-c7a5-401b-8460-d4576fefa456 |
| ***Astragalus galegiformis*** | Atha | 3704 | NY | Georgia | 2003 | cf2262c2-99f6-48e2-87eb-9716d00bbe8a |
| ***Astragalus gambelianus*** | Jamaes | 1739 | NY | United States of America: California: Marin | 1946 | cb826c61-1395-46eb-9c19-5cb379544230 |
| ***Astragalus garbancillo*** | Anderson | 12347 | NA | Argentina: Jujuy: Humahuaca | 1982 | 16703338-ed70-4776-87a7-7357a24bcf04 |
| ***Astragalus geminiflorus*** | Peredes | 4579 | NY | Ecuador: Quito | 1966 | d1613cb0-6efb-4a8a-94fe-043dc9db22ce |
| ***Astragalus geniculatus*** | Jahandiez | 276 | NY | Morocco: F√®s-Mekn√®s: Taza | 1929 | d045bc91-02f9-4a39-a4d3-64f179d0090d |
| ***Astragalus gentryi*** | Van Devender | 95-380 | TEX | Mexico: Sonora: Y√©cora | 1995 | 2109def2-3589-4ce4-a2f7-14e5e42815d3 |
| ***Astragalus georgii*** | [Cyrillic] | 1364 | NY | Kazakhstan: Turkistan | 1977 | cd317342-0d05-4329-a851-fafc1a12edab |

| | | | | | | |
|---|---|---|---|---|---|---|
| ***Astragalus geyeri*** | Smith | 12528 | NY | United States of America: Idaho: Owyhee | 2015 | d70d4e4b-abdc-4d25-8e93-cf3cfc35c030 |
| ***Astragalus giganteus*** | Lebgue | 3526 | CAS | Mexico: Chihuahua: Gomez Farias | 1994 | 1602aae2-df0e-4108-9bc5-9a8133722131 |
| ***Astragalus gilensis*** | Landrum | 5253 | NY | United States of America: Arizona: Apache | 1986 | cb7f0c6e-3a75-4498-8138-f68f6ed1e941 |
| ***Astragalus gilmanii*** | Marrs-Smith | s.n. | NY | United States of America: Nevada: Lincoln | 1985 | d70858c5-b4aa-4e47-bc55-e423eb735ea8 |
| ***Astragalus glabrescens*** | [Cyrillic] | 1027 | NY | [Cyrillic] | 1960 | ceb5935a-e24c-4973-8732-7d978d17b2f7 |
| ***Astragalus glaux*** | Castroviejo | FC 5422 | MO | Morocco: Tetouan | 1981 | 153f0f1e-8613-4f04-afd7-6f7087855d1f |
| ***Astragalus globiceps*** | Butkov | 6256 | NY | Turkmenistan: Mary: Serhetabat | 1930 | cf12ba7e-8d4f-4529-a782-dfb7d0f0be01 |
| ***Astragalus globiceps (Astragalus agameticus)*** | Cuba | s.n. | NY | Russian Federation [Checheno-Ingush Autonomous Soviet Socialist Republic] | 1990 | ab9a2fd3-3d78-41fc-a573-c24a159c627a |
| ***Astragalus glumaceus*** | Ledingham | 4230 | NY | Islamic Republic of Iran: Kordestan: Sanandaj | 1965 | ab996d0d-10ca-443d-8574-a977a2d21f75 |
| ***Astragalus glycyphylloides*** | Karjagin | s.n. | NY | Azerbaijan | 1937 | cf2970ec-df7d-4af3-8534-f3542b76cf47 |
| ***Astragalus glycyphyllos*** | USDA-ARS | 234686 | NY | Denmark: Zealand: Kulhuse | 1971 | cb7e9efe-6c19-418f-ad9d-eb032fe918c2 |
| ***Astragalus goktschaicus*** | Sytin | 6532 | NY | Armenia: Gegharkunik | 1982 | cf2bb4a2-fc7d-4088-8c56-fd179f4cad32 |
| ***Astragalus goldmanii*** | McVaugh | 91 | CAS | Mexico: Aguascalientes | 1959 | 15f98320-9be4-4665-a824-2b29af731764 |
| ***Astragalus gombo*** | Herrero | AH3870 | MO | Tunisia: Gabes: Metlaoui | 2009 | 1536870e-a1f0-408b-9bfe-3a870c5c0ebb |
| ***Astragalus gracilis*** | Higgins | 5112 | NY | United States of America: Texas: Oldham | 1972 | d6f7a328-4f27-4616-94cd-82f9036ed4cf |
| ***Astragalus granatensis*** | Larsen | 36079 | MO | Italy: Sicily: Catania | 1977 | 2e7479b6-2a41-4964-8f7c-1abd2f233730 |
| ***Astragalus graveolens*** | Khan | 1098 | HUH | Pakistan: Islamabad | 1977 | f7eba16e-156b-4cb0-b1af-86a99ba2c64a |
| ***Astragalus grayi*** | Hartman | 10919 | NY | United States of America: Wyoming: Washakie | 1980 | d6eb77ee-4400-410a-823e-c5bfcb2def61 |
| ***Astragalus griffithii*** | Dieterle | 1272 | NY | Afghanistan: Bamyan | 1971 | ab7d85ae-157e-4da6-81b9-9423c003f098 |
| ***Astragalus gruinus*** | Wiggins | 16627 | CAS | Mexico: Baja California: Ensenada | 1961 | 15f37c46-b813-49cb-a3b8-609620666bd3 |
| ***Astragalus guatemalensis*** | Ruiz | 618 | CAS | Mexico: Chiapas: Chamula | 1988 | 157c1046-3cd9-4942-a39e-05d9d3dbc5cb |
| ***Astragalus gummifer*** | Rechinger | 11292 | MO | Iraq: Erbil: Choman | 1957 | 0c6743e4-0d25-4b30-aa0c-d0ebd8c92637 |
| ***Astragalus guttatus*** | Korovin | 20 | NY | Kazakhstan: Turkistan: Saryaghash | 1924 | cd451549-4999-41ed-9677-a5b447b7b232 |

| | | | | | | |
|---|---|---|---|---|---|---|
| ***Astragalus gypsodes*** | Holmgren | 6956 | NY | United States of America: New Mexico: Eddy | 1973 | d6e94ca3-9448-436c-85b6-5c6c6fdc010f |
| ***Astragalus hajastanus*** | Sytin | 6533 | NY | Armenia | 1981 | cf3fd5a2-31cf-45f7-b03c-c8bc57330543 |
| ***Astragalus hallii*** | Grimes | 2029 | NY | United States of America: Utah: Kane | 1981 | cb6b42df-4422-4310-826a-3eef4348ff77 |
| ***Astragalus hamiltonii*** | Goodrich | 27817 | NY | United States of America: Utah: Uintah | 2010 | cb613bdb-7b88-4659-b585-ad10d2cbb187 |
| ***Astragalus hamosus*** | Miller | 552 | MO | Morocco: SW | 1974 | 1533078d-1e49-45cb-8088-cb1bbd0d843a |
| ***Astragalus harbisonii*** | Breedlove | 71572 | CAS | Mexico: Baja California | 1991 | 15f239ba-7664-4b6a-8623-42034879183d |
| ***Astragalus harrisonii*** | Welsh | 21180 | NY | United States of America: Utah: Wayne | 1982 | d6e872e1-ee19-4e3d-9139-81406486e4ef |
| ***Astragalus hartwegii*** | Ripley | 14154 | CAS | Mexico: Zacatecas: Sombrerete | 1965 | 156a5fc3-aebd-4cd5-baa0-a077b6915acb |
| ***Astragalus havianus*** | Boufford | 39058 | HUH | People’s Republic of China | 2007 | fa75b2c0-ec09-4199-b3ee-c8087e138b3a |
| ***Astragalus hemsleyi*** | Freitag | 5772 | NY | Afghanistan: Logar | 1969 | ab66f5c1-6221-463a-a53c-e138dba5e4fb |
| ***Astragalus heptapotamicus*** | Goloskokov | 5771 | HUH | Kazakhstan: Almaty | 1971 | fa755eb9-7ca4-48b4-b017-bb4c322f48f7 |
| ***Astragalus heterodontus*** | Gusev | 4582 | NY | Tajikistan: Pamir | 1958 | cea31600-9245-4ab5-8750-8ca85aaeef5a |
| ***Astragalus hidalgensis*** | Delgado | 1117 | CAS | Mexico: Hidalgo: Zimapan | 1979 | 160ebf3c-7a60-4564-9345-41a8ac853704 |
| ***Astragalus himalayanus*** | Suzuki | 9191431 | HUH | Nepal: Seti Zone: Bajhang Distr. | 1991 | f8181a2b-b73b-4b15-b1e6-600f0f356280 |
| ***Astragalus hispidulus*** | Pitard | 87 | NA | NA: NA: NA | NA | 157dce89-f110-454c-a498-daf58d1c34ae |
| ***Astragalus hissaricus*** | [Cyrillic] | 1646 | HUH | [Cyrillic] | 1960 | fa46b711-95e7-4cff-850c-1dcbee23afcb |
| ***Astragalus hoantchy*** | Morefield | 5141 | NY | People’s Republic of China: Xinjiang | 1989 | cea20bae-5855-4b67-987d-4b53af3f5b29 |
| ***Astragalus hoffmeisteri*** | Anders | 5092 | NY | Afghanistan: Kunar | 1970 | a9fb4d80-0213-4941-9abc-a743cfa71052 |
| ***Astragalus holmgreniorum*** | Atwood | 10742A | NY | United States of America: Utah: Washington | 1985 | d6e86156-1e63-4f8f-b07d-2602e902efea |
| ***Astragalus hololeios*** | Podlech | 30604 | HUH | Afghanistan: Kabul | 1978 | f8009163-1488-4fbf-8141-f313100d132a |
| ***Astragalus hoodianus*** | Halse | 3397 | NY | United States of America: Washington: Klickitat | 1987 | d6e770db-7dfd-4a20-a194-8c16e121f97c |
| ***Astragalus hornii*** | Harbison | s.n. | CAS | Mexico: Baja California | 1953 | 160bc74d-3091-41b8-9dc8-5db0dba4206f |
| ***Astragalus howellii*** | Ertter | 7581 | NY | United States of America: Oregon: Wasco | 1988 | cb5363b0-03a2-468a-a111-d96c331b098c |
| ***Astragalus humillimus*** | Neese | 13080 | NY | United States of America: Utah: Tooele | 1983 | cb4c0096-5edd-497c-9559-0c910ede3c58 |

| | | | | | | |
|---|---|---|---|---|---|---|
| ***Astragalus humistratus*** | Atwood | 25901 | NY | United States of America: New Mexico: Mckinley | 2000 | cb49ea87-eccc-4137-b975-c04e6295e2ae |
| ***Astragalus hyalolepis*** | Akhlerdov | s.n. | NY | Armenia | 1960 | cea15494-b378-4294-b11c-3895722aa6cb |
| ***Astragalus hypoglottis*** | Krendl | s.n. | NY | Italy: Tuscany: Lucca | 1981 | cf79c9f5-2fa9-4e98-8b50-ae6c085ebb6c |
| ***Astragalus hypoleucus*** | NA | NA | NA | Mexico: Tlaxcala: Tlaxcala | 1981 | 160637cb-b865-4456-8ae2-336cd013cf06 |
| ***Astragalus hypoxylus*** | NA | NA | NA | Mexico: Sonora: Y√©cora | 2004 | 14e11017-17cf-4e4a-a1a1-7a4c722c9c19 |
| ***Astragalus hypsogenus*** | Budin | LIL 71105 | NY | Argentina: Tucum√°n: Burruyac√∫ | 1930 | d1557db5-7e0d-4f14-a40c-6b0c99395cd0 |
| ***Astragalus hyrcanus*** | Tzelev | 3570151 | NY | Russian Federation: Daghestan: Caucasus | 1961 | cf5c4b06-03dd-4b66-a7c2-be0f362840d0 |
| ***Astragalus ibicinus*** | Ledingham | 4086 | NY | Islamic Republic of Iran: Fars | 1965 | a9f3ca97-18aa-4721-b3c3-21efbaa9fa69 |
| ***Astragalus ibrahimianus*** | Reading Univ. | 750 | MO | Morocco: GA: Oukaimeden | NA | 1524c81a-b786-4521-a28e-83b4bc3c05f1 |
| ***Astragalus idrietorum*** | Breedlove | 60775 | CAS | Mexico: Baja California: Ensenada | 1984 | 1603e409-eda8-47db-b11f-8a2d900cf308 |
| ***Astragalus igniarius*** | Hajiyev | 3 | NY | Azerbaijan: Baku | 2006 | d1e8915e-d3ec-4071-97dc-23241cdc3f5a |
| ***Astragalus illinii*** | Leon | 3453 | NY | Argentina: Chubut: Gastre | 1983 | d1556547-043d-4067-a698-cb3dcb450ff8 |
| ***Astragalus incanus*** | Miller | 703 | MO | Morocco: AA | 1974 | 1522c83e-4fdc-42fd-ac01-2b2c5d1ef9cb |
| ***Astragalus incertus*** | [Cyrillic] | | NY | Armenia | 1965 | cea147f7-0a03-486a-855e-38ecd2745a69 |
| ***Astragalus indurescens*** | Vassiljeva | | NY | Uzbekistan | 1962 | ce9f51c7-7465-42b1-9fc2-703128a3eedb |
| ***Astragalus inflexus*** | Fishbein | 3328 | NY | United States of America: Washington: Asotin | 1998 | cb3a480d-2805-42a6-8857-d49c7e04b8a3 |
| ***Astragalus insularis*** | Breedlove | 62490 | CAS | Mexico: Baja California | 1986 | 15e60a7f-f802-4e22-9dfc-6a0370a9e97b |
| ***Astragalus inversus*** | Schoolcraft | 2093 | NY | United States of America: California: Lassen | 1990 | d6c77a8c-6d19-48b8-9e4a-86aa9a609ece |
| ***Astragalus inyoensis*** | Twisselmann | 15545 | NY | United States of America: California: Inyo | 1969 | cb35a1e8-3b0f-4004-a530-1baeac99cc53 |
| ***Astragalus iodopetalus*** | Barneby | 17879 | NY | United States of America: New Mexico: Rio Arriba | 1983 | cba14a51-6b56-4c14-a620-d2bea8a262b3 |
| ***Astragalus irolanus*** | Samuelsson | 4390 | MO | Syrian Arab Republic: Rif Dimashq | 1933 | 0c2e1b8e-62b8-444e-84a6-2bfdbc6102a9 |
| ***Astragalus iselyi*** | Tuhy | 3880 | NY | United States of America: Utah: San Juan | 2006 | cba10d3c-ef65-4bb4-bcd1-0874dc650a4e |
| ***Astragalus iskanderi*** | Ledingham | 3303 | NY | Canada: Saskatchewan | 1962 | cd52e99d-953d-40ec-952d-aa239b2d52ef |
| ***Astragalus jaegerianus*** | Huggins | 13-132 | NY | United States of America: California: San Bernardino | 2013 | cb9f37b2-7c53-4c13-8a20-0e1d365c91c2 |
| ***Astragalus jaliscensis*** | McVaugh | 18468 | CAS | Mexico: Aguascalientes | 1960 | 16316cd4-9906-4a1e-b0c0-bc9026fb7b89 |

| | | | | | | |
|---|---|---|---|---|---|---|
| ***Astragalus jejunus*** | Tiehm | 9665 | NY | United States of America: Nevada: Elko | 1985 | cb9e7b4d-6ab2-41f3-b46c-2db75027f746 |
| ***Astragalus johannis*** | Ledingham | 4114 | NY | Islamic Republic of Iran: Fars | 1965 | ab666b01-925c-428c-be85-e38976c97ecb |
| ***Astragalus johannis-howellii*** | Honer | 316 | NY | United States of America: California: Mono | 2000 | d6b92c54-e776-49dc-ac64-c42a8e7b9622 |
| ***Astragalus jolderensis*** | Popov | 6259 | HUH | Turkmenistan: Kopetdag | 2018 | ffc1f985-384b-4112-a5b7-a83b17aeb489 |
| ***Astragalus josephi*** | Rock | 17937 | HUH | People's Republic of China: Southwestern Szechwan: Muli (or Mili) Kingdom | 1929 | 0067d468-4031-4c60-b5dd-a7bcd1c8663a |
| ***Astragalus juratzkanus*** | [Cyrillic] | s.n. | HUH | [Cyrillic] | 1925 | 00bdfc36-a0b1-418c-a5b7-94afee749c88 |
| ***Astragalus kabadianus*** | Vvedensky | 225 | HUH | Uzbekistan: Khadzha-Ipak: Pamir-Alay | 1930 | 0046e608-c288-4662-94b4-26ea027cb690 |
| ***Astragalus kadschorensis*** | Lachashvili | 142 | NY | Georgia | 2005 | d0bd5d31-873b-4a9e-976f-2411b11a43df |
| ***Astragalus kahiricus*** | Grant | 15,356 | NY | Islamic Republic of Iran: Sistan and Baluchestan: Zahedan | 1964 | ab664be0-9827-4d06-b4a1-3d8602719271 |
| ***Astragalus karabaghensis*** | [Cyrillic] | s.n. | NY | [Cyrillic] | 1955 | cd5765f9-d90f-4df8-8f1f-76c00c3eb406 |
| ***Astragalus karakugensis*** | Tzvelev | 7134 | NY | Russian Federation: Dagestan | 1961 | cd54666e-4826-4263-a0a9-84dc760b45db |
| ***Astragalus karakuschensis*** | [Cyrillic] | s.n. | NY | Armenia | 1955 | cd59d4ed-cf7b-48d2-8a17-661059938a11 |
| ***Astragalus kasachstanicus*** | Ragkovska | 8157 | NY | Kazakhstan: Karaganda | 1966 | cd5ca40c-f18b-493a-9ae9-b78f202a6d95 |
| ***Astragalus kaschkadarjensis*** | Vvedensky | 6217 | HUH | Uzbekistan | 1929 | 0042832f-3b6f-4f9c-835d-e2e9f1d464fa |
| ***Astragalus kashmarensis*** | Singh | 2771 | NY | India: Ladakh | 1971 | ab5b90c8-ae28-47de-9065-b546426154e3 |
| ***Astragalus kazbeki*** | Abdaladze | 241 | NY | Georgia: Mtskheta-Mtianeti: Kazbegi | 2004 | d0d553dc-d0ac-4bef-9404-a83702a29b0d |
| ***Astragalus kelifii*** | Vvedensky | 387 | NY | Uzbekistan: Surxondaryo | 1927 | cd5dcd50-24d8-4b48-8211-3c4273e79f0f |
| ***Astragalus kentrophyta*** | Smith | 5413 | NY | United States of America: Idaho: Owyhee | 2005 | cb9df1ff-502f-42ea-a53b-0ee9e45fdd31 |
| ***Astragalus kialensis*** | D.E. Boufford | 29915 | HUH | People's Republic of China | 2000 | 003604b7-3fc3-4d9d-8246-cfb168facc7a |
| ***Astragalus kifonsanicus*** | [None] | | HUH | People's Republic of China: Sung.: Tsinan | 1924 | 00344364-90d4-4d32-83ae-3452b96ad573 |
| ***Astragalus kirrindicus*** | Grant | 15,77 | NA | Islamic Republic of Iran: Fars: Shiraz | 1964 | e5b65a2a-fe55-4747-9596-e271125243e5 |
| ***Astragalus knightii*** | Knight | 1922 | NY | United States of America: New Mexico: Sandoval | 1982 | cb99c9ad-6b5b-4fdf-a78e-8819018579d6 |

| | | | | | | |
|---|---|---|---|---|---|---|
| ***Astragalus kolymensis*** | Korobkov | s.n. | NY | Russian Federation: Chukotka: Anadyrsky | 1979 | ce99b062-a55e-4865-9b41-d1d82cf0d1b6 |
| ***Astragalus krauseanus*** | Baranov | s.n. | HUH | Uzbekistan | 1923 | 00211a6f-dbc5-4323-bdb5-20631e1280c0 |
| ***Astragalus kudrjaschovii*** | Borissova | 37 | HUH | Russian Federation | 1960 | 009dd2d9-7e7d-40cb-baa9-d51d183db794 |
| ***Astragalus kulabensis*** | [Cyrillic] | 1120 | HUH | Tajikistan | 1960 | 00b28180-39bf-46fe-b2a1-a1ee4a889123 |
| ***Astragalus kuramensis*** | Volk | 71/844 | NY | Afghanistan: Paktia | 1971 | ab480268-1737-477e-a026-7abbd2798914 |
| ***Astragalus kurdaicus*** | G. F. Ledingham | 3477 | NY | Canada: Saskatchewan | 1963 | ce943280-1f8f-4042-87f8-efc00c7f8f4f |
| ***Astragalus kuschakewiczii*** | Boufford | 27861 | HUH | People's Republic of China: Sichuan | 1997 | 00cb1843-4ba5-44eb-94f4-841bf04933bd |
| ***Astragalus kuschakewiczii (Astragalus mongutensis)*** | Morefield | 5243 | NY | People's Republic of China: Xinjiang Uygur Zizhiqu | 1989 | b0eaf9cd-e712-4d27-b1e4-518bb765275b |
| ***Astragalus kuschkensis*** | Butkov | 6218 | HUH | Turkmenistan | 1946 | 00c2473d-f954-4d92-a23f-0cc1ce92e961 |
| ***Astragalus laetus*** | Volk | 147 | NY | Afghanistan: Kabul | 1950 | ab437cf7-91a5-46e3-ace9-c7180a6759b9 |
| ***Astragalus lagopoides*** | Lamond | 4098 | NY | Islamic Republic of Iran: West Azerbaijan | 1971 | ab29bc5a-07be-4c39-a0c6-eecd69358aee |
| ***Astragalus laguriformis*** | Ledingham | 4212 | NY | Islamic Republic of Iraniran | 1965 | ab297796-5773-4e65-84ec-216b7149d65e |
| ***Astragalus laguroides*** | [Cyrillic] | 90 | HUH | Russian Federation | 1979 | fa206d78-8fbb-4d2c-8ed4-f3685a6aeecb |
| ***Astragalus lanuginosus*** | Goloskokov | 5772 | NY | Kazakhstan | 1971 | ce8350c4-4172-4529-a11f-34ec466b0d38 |
| ***Astragalus lasiosemius*** | Anders | 7234 | NY | Afghanistan: Badakhshan | 1971 | ab262957-a407-41cd-86a9-747f1d3136a7 |
| ***Astragalus lasiostylus*** | Popov | 6261 | HUH | Uzbekistan: Samarqand: Urgut | 1928 | 0062e39a-5693-4b7f-be19-d45a80c6d33d |
| ***Astragalus laxmannii (Astragalus adsurgens)*** | Boufford | 27113 | HUH | People's Republic of China: Beijing | 1996 | 001e1e76-135d-424b-bec1-e45967b897ff |
| ***Astragalus laxmannii*** | Boufford | 40308 | HUH | People's Republic of China: Sichuan: Ruoergai | 2007 | 0059ba0b-8ef5-4832-bf03-14b48ab61fc0 |
| ***Astragalus layneae*** | Higgins | 8367 | NY | United States of America: California: San Bernardino | 1974 | cb83f291-bf88-4a32-84d3-1c31f018dd5f |
| ***Astragalus leibergii*** | Spellenberg | 1665 | NY | United States of America: Washington: Kittitas | 1967 | d6a4e5b5-3d91-4dc6-aeda-c190467edd02 |
| ***Astragalus lemmonii*** | Ertter | 6722 | NY | United States of America: California: Mono | 1986 | d74c4143-b0a3-40ef-9a86-5796f088d796 |
| ***Astragalus lentiformis*** | Quesenberry | 32.1 | FLAS | United States of America: California: Plumas | 2006 | e941b3ca-c0ad-4514-b605-d7684001bd70 |

| | | | | | | |
|---|---|---|---|---|---|---|
| ***Astragalus lentiginosus*** | Lehto | L21308 | FLAS | United States of America: Arizona: Not shown | 1977 | e9ab5881-86ad-4d25-8b05-4e7dab85d40f |
| ***Astragalus lentiginosus (Astragalus iodanthus)*** | Holmgren | 4130 | TEX | United States of America: Nevada: Pershing | 1970 | 21607353-8d63-4b7c-a97f-4d6ce0dfb61f |
| ***Astragalus leontinus*** | Pistarino | s.n. | OS | Italy: Valle d'Aosta | 1986 | e59f67fa-416d-4651-985c-375a5a0cd7f9 |
| ***Astragalus lepsensis*** | Roldugin | 4819 | HUH | Kazakhstan | 1964 | 0034c885-0543-425a-a6d8-592b7795bbad |
| ***Astragalus leptocarpus*** | Thomas | 103,645 | FLAS | United States of America: Arkansas: Ouachita | 1988 | e9a49f11-2742-4c3a-8c48-1ed9155c3377 |
| ***Astragalus leptocaulis*** | Schischkin | 3717 | HUH | Kazakhstan: East Kazakhstan: Oskemen | 1931 | 0016f6ce-c875-41c8-b0e0-06f00661b527 |
| ***Astragalus leucocephalus*** | Shahzad | 134 | HUH | Pakistan: Punjab: Rawalpindi | 1976 | f7f3de28-f614-4337-b50e-e60df68e95c5 |
| ***Astragalus leucolobus*** | Thorne | 51946 | NY | United States of America: California: San Bernardino | 1978 | d57dac91-feff-4689-84d6-d58306b9959d |
| ***Astragalus lilacinus*** | W6 AMD | 126 | NY | Islamic Republic of Iran: North khorasan: Maneh and Samalqan | 1977 | aafe5a64-67b5-4eec-b9c8-fc1b3d073473 |
| ***Astragalus limariensis*** | Jiles | 2831 | OS | Chile: Coquimbo (IV): Choapa | 1955 | e457d48c-d384-4c42-b876-d7a01e186c2c |
| ***Astragalus limnocharis*** | Isely | 11475 | NY | United States of America: Utah: Kane | 1971 | d78d267b-be70-45d6-a119-1e244374a825 |
| ***Astragalus lindheimeri*** | Barneby | 18280 | NY | United States of America: Texas: Stonewall | 1989 | cc0008c0-ec0a-4545-918e-4bf8887906be |
| ***Astragalus lineatus (Astragalus gezeldarensis)*** | [Cyrillic] | s.n. | NY | Armenia | 1955 | cd436bf4-a72e-43cf-918a-c6988dd20779 |
| ***Astragalus lineatus (Astragalus grammocalyx)*** | Ledingham | 4167 | NY | Islamic Republic of Iran | 1965 | ab872349-0994-4645-896e-5e70dd1de4fc |
| ***Astragalus linifolius*** | Atwood | 8742 | NY | United States of America: Colorado: Mesa | 1982 | cb833c0c-068a-4bbf-ac89-fb367f4c6bbc |
| ***Astragalus lipskyi*** | Kamelin | 6220 | HUH | Uzbekistan | 1930 | 00400878-9697-4bc4-9024-0b031cb4d8fd |
| ***Astragalus litwinowii*** | Ledingham | 4824 | NY | Canada: Saskatchewan | 1966 | ce7aaaec-2a82-4350-8b09-4185a71d9b38 |
| ***Astragalus loanus*** | Welsh | 26797 | NY | United States of America: Utah: Sevier | 1997 | cc007253-cd08-4576-880a-270e29e0355e |
| ***Astragalus lonchocarpus*** | Atwood D | 8732 | NY | United States of America: Nevada: Lincoln | 1982 | d786c784-6dce-4991-adce-3fd7b125630e |
| ***Astragalus longipetalus*** | Skvortsov | | NY | Russian Federation: Volgograd: Dubovsky | 1967 | cf5de8f0-82e4-4938-9dfc-d8dc2c6782ab |
| ***Astragalus longipetiolatus*** | Klinkova | s.n. | NY | Russian Federation: Astrakhan | 1990 | cd74498f-550d-4cc8-b910-62aca8862062 |

| | | | | | | |
|---|---|---|---|---|---|---|
| ***Astragalus longissimus*** | Reina | 97-618 | CAS | Mexico: Sonora: Yecora | 1997 | 14db4077-376f-486b-aee3-ade6c0114e54 |
| ***Astragalus looseri*** | Kiesling | 7696 | NY | Argentina: San Juan: Calingasta | 1991 | d12b1581-e70e-4e9b-acc5-92d5fd203fa2 |
| ***Astragalus lotiflorus*** | Boivin | 9984 | NY | Canada: Saskatchewan: Swift Current | 1952 | d7864769-e11c-471d-9ef8-824427365056 |
| ***Astragalus lumsdenianus*** | Podlech | 29566 | NY | Afghanistan: Herat | 1977 | aaf3adc5-852c-435e-a6b9-430766dfb424 |
| ***Astragalus lupulinus*** | Popov | 3823 | NY | Russian Federation | 1953 | adae4e0b-7d34-46ef-9287-fc8293b3e386 |
| ***Astragalus lutosus*** | Franklin | 7298 | FLAS | United States of America: Utah: Wasatch | 1990 | e9a2b0d0-504d-49ec-bbee-201909aa23e3 |
| ***Astragalus lyallii*** | Fishbein | 3367 | NY | United States of America: Washington: Not shown | 1998 | d7831d5e-f4e0-449d-af11-736892fca126 |
| ***Astragalus macrobotrys*** | Vasak | s.n. | NY | Turkmenistan | 1984 | adac4ddb-2e8f-44ba-adb7-64cbbb7d3d8c |
| ***Astragalus macrocephalus*** | Astragalus Macropetalus | 6222 | HUH | Not shown | 1923 | fefc62f9-45a6-4e15-b780-12e926904018 |
| ***Astragalus macrocladus*** | Popov | 6221 | HUH | Tajikistan: Kaznak | 1924 | 0032ece3-50b0-460b-acbd-f2395cec37e2 |
| ***Astragalus macrodon*** | Ledingham | 4040 | NY | United States of America: California: Kern | 1964 | d78113b6-9eee-4f25-9a2d-03e6d46eb248 |
| ***Astragalus macropelmatus*** | Ledingham | 4198 | NY | Islamic Republic of Iran: Kermanshah: Harsin | 1965 | aaf31dad-500d-4958-9f33-36b94d3af6c2 |
| ***Astragalus macropodium*** | [Cyrillic] | 1624 | HUH | Russian Federation | 1960 | 00a424e7-9aeb-4e37-a05d-187be42e8196 |
| ***Astragalus macropterus*** | Orazova | 5377 | HUH | Kazakhstan: Shymkent: Shymkent | 1968 | 00fa1b32-39b8-4c36-a7d1-5407169abd30 |
| ***Astragalus macropus*** | Sagalaev | | NY | Russian Federation: Saratov: Pereljub | 1993 | a9dcd1cd-2034-4509-bbf6-c931feb855d9 |
| ***Astragalus macrostachys*** | Ledingham | 5200 | NY | Canada: Saskatchewan | 1969 | b020cdeb-acb0-4836-856e-57837ed820f3 |
| ***Astragalus macrotropis*** | Blak | 3319 | NY | Kazakhstan: Zhambyl | 1909 | adae75d7-3ff6-4a72-b58e-2b63a2666c06 |
| ***Astragalus magdalenae*** | Felger | 12450 | CAS | Mexico: Sonora: Hermosillo | 1965 | 166cafdb-4327-458f-a62d-0ec50592628d |
| ***Astragalus mahoschanicus*** | Ho | 1466 | HUH | People’s Republic of China | 1993 | 00d221dd-cbc6-4e59-af08-a2b07e5c3235 |
| ***Astragalus malacoides*** | Welsh | 21188 | NY | United States of America: Utah: Garfield | 1982 | d7794608-3380-46b9-8201-cb93c584fda7 |
| ***Astragalus malacus*** | Otting | 511 | NY | United States of America: Oregon: Malheur | 2003 | cbf0adcd-c1c8-4da7-a01e-42fd8b0436cf |
| ***Astragalus managildensis*** | Sovetkina | 389 | NY | Kazakhstan | 1925 | b0200651-36d1-4398-9683-fa1baca1ab0b |
| ***Astragalus mareoticus*** | Podlech | 35210 | NY | Algeria: Adrar: Timimoun | 1981 | aae97c8e-6a60-4ec4-b284-dea2da0ad0b6 |
| ***Astragalus maroccanus*** | M'sabih Talaa | 102 | NY | Morocco: montana: not shown | 1986 | d03fec64-63a2-4655-b7f7-780d126a6a3c |

| | | | | | | |
|---|---|---|---|---|---|---|
| ***Astragalus masenderanus*** | Bornmuller | 6855 | NY | Islamic Republic of Iran | 1902 | b076641e-4835-445d-a406-e8b8f30f2883 |
| ***Astragalus mattam*** | M. Gilbert | 1286 | HUH | People's Republic of China | 1993 | 00cf8891-0e51-421a-b6ce-8b65e2d46637 |
| ***Astragalus maurorum*** | Lippert | 21917 | NY | Morocco: Taza | 1987 | d03f68bb-3d34-4d6b-904f-b149a3cf6088 |
| ***Astragalus maximowiczii*** | Butkov | 6224 | HUH | Turkmenistan | 1930 | 00ba1fb6-feba-4bdb-83f9-9af440cd07b5 |
| ***Astragalus maxwellii*** | Kanai | 852281 | HUH | Japan | 1972 | f81dbfe0-92c0-4b04-860d-f7285cee1b3e |
| ***Astragalus medius*** | Androssov | | NY | Kazakhstan | 1909 | b103229b-6984-40c0-acf5-6aa6f7f0a309 |
| ***Astragalus megacarpus*** | Tiehm | 14272 | NY | United States of America: Nevada: Elko | 2003 | cbeac4a8-32e3-46f9-882d-96e54e41c283 |
| ***Astragalus megalanthus*** | Schischkin | s.n. | NY | Kazakhstan: East Kazakhstan: Oskemen | 1931 | b0f70e84-3b7f-44bc-8d86-cc28822b90c8 |
| ***Astragalus megalomerus*** | Pojarkova | 1177/216 | HUH | Tajikistan | 1944 | 00d3f1d7-ba08-4b84-b0b0-3d30e520e201 |
| ***Astragalus melanostachys*** | Koelz | 2910 | NY | India | 1931 | aae976f3-98cc-4f80-ad11-b137f2c207a7 |
| ***Astragalus melilotoides*** | Bartholomew B. | 2075 | HUH | People's Republic of China: Beijing | 1984 | 00b483eb-8a6c-4767-8c3d-2c1261984c74 |
| ***Astragalus mesites*** | Abemuceu | | NY | [Cyrillic] | 1956 | adaaa596-14fe-4edb-a317-0bf947ee1646 |
| ***Astragalus michauxii*** | William B Fox | 4639 | FLAS | United States of America: North Carolina: Cumberland | 1951 | ea2d1432-591a-4304-9022-00c51132cd23 |
| ***Astragalus micranthellus*** | Ricardi | 30266 | OS | Chile: Tarapaca: Iquique | 1961 | e44840bc-2b6a-4271-9c0c-026401aec941 |
| ***Astragalus micranthus*** | Hernandez | s.n. | CAS | Mexico: Mexico | 1967 | 165e73d7-9d01-448d-abfe-14b7c2df5380 |
| ***Astragalus microcalycinus*** | Lamond | 2029 | NY | Afghanistan | 1965 | aad78ff4-f5c2-4362-aed6-5da131741bda |
| ***Astragalus microcephalus*** | Hageskuna | 152 | NY | Russian Federation: Leningrad: Northwestern | 1953 | af5b069d-ac93-489f-80a7-3aae3ba4a1b9 |
| ***Astragalus microcystis*** | Lesica | 4816 | NY | United States of America: Montana: Sanders | 1989 | cbe42b45-89e7-42c5-8a95-56f7bed06a40 |
| ***Astragalus micromerius*** | Atwood | 30432 | NY | United States of America: New Mexico: Mckinley | 2004 | d775a89e-6129-484d-86ea-e5bece8819e6 |
| ***Astragalus microphysa*** | Assadi | 1838 | NY | Islamic Republic of Iran: KERMAN: Unknown | 1977 | cd613e04-1576-4605-8222-4f6cd18d0ff1 |
| ***Astragalus micropterus*** | Donmez | 2776 | NY | Turkey | 1990 | aa7fc535-6266-4ef1-a0f0-ef5d4aa4e280 |
| ***Astragalus miguelensis*** | R.M Beauchamp | 22703 | FLAS | United States of America: California | 1975 | e9a0dcb6-2e63-49ba-9cb8-b040b72ccf38 |
| ***Astragalus miniatus*** | [Cyrillic] | [?]738 | NY | Mongolia | 1941 | ada40d54-201a-4d41-93cb-5463c05c0b3d |
| ***Astragalus minimus*** | Asplund | 2672 | NY | Bolivia (Plurinational State of): La Paz: Pacajes | 1921 | d12a799b-568f-4e9f-9b5d-2860f4bb038a |

| | | | | | | |
|---|---|---|---|---|---|---|
| ***Astragalus minthorniae*** | Clokey | 7575 | FLAS | United States of America: Nevada: Clark | 1937 | e9a09e30-60dd-49ad-a68b-d4c238dd5996 |
| ***Astragalus minutidentatus*** | Iokawa | 20310064 | HUH | Nepal | 2003 | f7daa87d-9c94-4450-b671-5e62f1c85d35 |
| ***Astragalus minutissimus*** | Marticorena | 30228 | OS | Chile: Tarapaca | 1964 | e5303619-f497-41fa-b110-7fef53167072 |
| ***Astragalus mirabilis*** | Rassulova | 62 | NY | Tajikistan | 1976 | aeb01698-7b96-4ed9-ac81-0e1f802f1e8d |
| ***Astragalus misellus*** | Camp | 904 | NY | United States of America: Washington: Klickitat | 1982 | d769abcb-df0d-448e-8293-86df9d3c5b80 |
| ***Astragalus miser*** | Foster | 12038 | NY | United States of America: Idaho: Power | 2001 | cbe11c1e-1bb7-44f6-998f-3244f1735d06 |
| ***Astragalus missouriensis*** | Thorne | 18488 | FLAS | United States of America: Iowa: Woodbury | 1957 | e99bf887-c6bb-4971-9cd6-0d110006af97 |
| ***Astragalus moencoppensis*** | Cronquist | 9198 | NY | United States of America: Utah: Emery | 1961 | d7616b4c-9eed-445f-ab1c-a7b130e5052f |
| ***Astragalus mohavensis*** | Leary | 5952 | NY | United States of America: Nevada: Nye | 1999 | cbaf813a-a735-47de-9cb5-275ba4aff8b2 |
| ***Astragalus mokiacensis*** | Atwood | 12165 | NY | United States of America: Utah: Washington | 1986 | cba34a4d-3f20-4c18-9e18-227ca9b55a14 |
| ***Astragalus mollissimus*** | Lyonnet | 540700001 | CAS | Mexico: Mexico | 1954 | 14dd7129-aab8-457b-9ca5-91e0d971ff20 |
| ***Astragalus molybdenus*** | Wilken | 14124 | NY | United States of America: Colorado: Gunnison | 1984 | cc2c08dd-0f0f-4012-8a99-9291e1107a18 |
| ***Astragalus monadelphus*** | Ching | 427 | NY | People’s Republic of China: Kansu | 1923 | b0ec3328-36ba-4cf2-af86-cd2570c5e0a1 |
| ***Astragalus monanthemus*** | Ledingham | 4095 | NY | Islamic Republic of Iran: Gardaneh Nabaty | 1965 | ab1cd250-5c7c-46fb-9d19-f5f2ac6d69e7 |
| ***Astragalus monbeigii*** | Boufford | 27351 | HUH | People’s Republic of China: Sichuan | 1997 | 01778c1b-e56f-467e-94fa-e30498681fab |
| ***Astragalus mongholicus*** | Boufford | 33843 | HUH | People’s Republic of China | 2005 | 015d9362-f48f-4653-8b84-7f4c8978bb50 |
| ***Astragalus mongholicus (Astragalus penduliflorus)*** | Elias | 7223 | MO | Russian Federation: Altai: Gorno-Altaysk | 1983 | 0bce7b14-583b-4d27-bf54-d3af73de58df |
| ***Astragalus mongholicus (Astragalus propinquus)*** | Ho | 1984 | HUH | People’s Republic of China: Qinghai | 1996 | 017e14ae-9939-4a3e-88e0-1e95de46affb |
| ***Astragalus mongholicus (Astragalus propinquus)*** | Ikonnikov-Galitzky | 2573 | HUH | Mongolia | 1929 | 010007ef-30bd-4fac-b67e-134794a7b2e3 |
| ***Astragalus mongholicus (Astragalus penduliflorus)*** | Schastin | | HUH | Mongolia | [None] | 013b6f7c-b9a1-42ea-94f3-4b2c455b5015 |
| ***Astragalus monoensis*** | Spellenberg | 2877 | NY | United States of America: California: Mono | 1972 | cc23462b-8ba6-405c-b221-d284736b2234 |
| ***Astragalus monspessulanus*** | Bornmuller | 3878 | NY | Macedon | 1918 | d2200f72-0e7b-4e8e-a76f-fd80f115fda7 |

| | | | | | | |
|---|---|---|---|---|---|---|
| ***Astragalus monticola*** | Kiesling | 8073 | OS | Argentina: San Juan: Calingasta | 1992 | e52c9920-e5c5-4e70-aa6f-0226ea026ffe |
| ***Astragalus monumentalis*** | Knight | 1397 | NY | United States of America: New Mexico: San Juan | 1981 | d75cca14-6f86-4d5b-af24-e9edf2ff5ad2 |
| ***Astragalus moranii*** | Moran | 23268 | TEX | Mexico: Baja California: Ensenada | 1976 | 20f4e16d-0e0e-4be3-b39f-3b3ee1c9a021 |
| ***Astragalus mucidus*** | Vvedensky | 13 | NY | Uzbekistan: Taschkent | 1922 | b0e570e3-86c2-4b53-911a-022083c37074 |
| ***Astragalus mucronifolius*** | Bokhari | 2091 | NY | Islamic Republic of Iran: Fars | 1977 | ab10775c-01bd-48dc-af7b-6fe2cf497f32 |
| ***Astragalus mugosaricus*** | Rusanov | 533 | NY | Kazakhstan: Aktyubinskaya | 1927 | b0e10e5d-84e2-496e-9b1e-8d8390b68e9a |
| ***Astragalus mulfordiae*** | Rosentreter | 4890 | NY | United States of America: Oregon: Malheur | 1988 | d75d9a78-f94a-44cf-b87c-5032928debb2 |
| ***Astragalus muliensis*** | Boufford | 29809 | HUH | People's Republic of China | 2000 | 0155f375-5224-4e7a-a351-3a6d9a07d17d |
| ***Astragalus munroi*** | G .Singh | 4443 | NY | India: Kashmir: Ladakh | 1971 | ab0c39c4-eaf2-43ae-a20f-bffbab6ed559 |
| ***Astragalus murinus*** | G. Ledingham | 4149 | NY | Islamic Republic of Iran | 1965 | cd8dee03-0837-40a9-9e94-d06c1bc126cb |
| ***Astragalus muschketovii*** | Puczkova | 6265 | HUH | Not shown: Not shown: Not shown | 1968 | 01472c90-602c-44bc-8831-a2a53471c23c |
| ***Astragalus musiniensis*** | Higgins | 1253 | NY | United States of America: Utah: Emery | 1968 | cc1491af-2880-419c-b46f-e37d184e2fe9 |
| ***Astragalus namanganicus*** | Popov | 391 | NY | Uzbekistan | 1926 | b0df7dd2-ff05-49e3-8f11-0933962bc39f |
| ***Astragalus naturitensis*** | Hartman | 89334 | NY | United States of America: Colorado: Mesa | 2010 | d74d4cca-5c72-48ee-9f27-cce4d3375673 |
| ***Astragalus neglectus*** | Boivin | 12907 | NY | Canada: Manitoba | 1958 | d747ad41-f6a8-4c4e-9f2e-c1d94a657ebe |
| ***Astragalus nelsonianus*** | Huber | 5428 | NY | United States of America: Wyoming: Sweetwater | 2016 | d5061049-5e55-45da-b34d-8c02149f5e93 |
| ***Astragalus nematodes*** | Goloskokov | 4473 | NY | Kazakhstan: Zhambyl | 1963 | b0db40f3-1d7b-4687-a4ae-43ef0a95353e |
| ***Astragalus neolipskyanus*** | Goloskokov | 4474 | NY | Kazakhstan: Zhambyl | 1963 | b0d00aab-913e-4d0e-b7bd-20a41bbadedb |
| ***Astragalus neomexicanus*** | Spellenberg | 2091 | NY | United States of America: New Mexico: Otero | 1969 | d50474b6-d1dc-40a5-b7ce-04d4c1a0823f |
| ***Astragalus neomonadelphus*** | Boufford | 30414 | HUH | People's Republic of China: Sichuan: Xiangcheng | 2004 | 013c2314-9bbc-4557-9fd3-609ba222f37e |
| ***Astragalus neuquenensis*** | Selander | 88-A-24 | NY | Argentina: Chubut: Paso de Indios | 1987 | d1088018-b950-4cbe-bc9d-edf9743b38cf |
| ***Astragalus neurophyllus*** | Borissova | 1631 | HUH | [Cyrillic] | 1960 | 01388adf-b87e-4a54-b630-6c8ec31386dc |
| ***Astragalus nevinii*** | Moran | 22704 | NY | United States of America: California: Los Angeles | 1975 | d5008e71-641f-43ae-a7ae-cdef81471e9a |

| | | | | | | |
|---|---|---|---|---|---|---|
| ***Astragalus newberryi*** | Pinzl | 9496 | NY | United States of America: Nevada: White Pine | 1991 | cc0a2ee1-b03a-4b01-a20e-ed0fe22d2c19 |
| ***Astragalus nicolai*** | Goloskokov | 4327 | NY | Kazakhstan | 1959 | b0cb426f-a28f-47ca-a838-87504f6a4b42 |
| ***Astragalus nidularius*** | Welsh | 9889 | NY | United States of America: Utah: Wayne | 1970 | d5d77caf-607d-4a9e-9164-e1fbfa164443 |
| ***Astragalus nigricans*** | Linczevski | 6226 | HUH | Turkmenistan: Badhyz | 1930 | 0134cf13-078a-4e6c-bd47-b632003a219a |
| ***Astragalus ninae*** | Androssov | 3321 | NY | Kazakhstan | 1908 | b0c43d88-2bd5-4c06-86e8-6dcd80391813 |
| ***Astragalus nivalis*** | Ho | 1136 | HUH | People's Republic of China: Qinghai | 1993 | 0113b67a-22d5-4b70-bef0-ce1256057a3b |
| ***Astragalus nivicola*** | Guerrido | 443 | NY | Argentina: Santa Cruz | 2001 | d1165230-4568-4f64-aefd-dff4de319505 |
| ***Astragalus nobilis*** | Zakrzewski | 6227 | HUH | Uzbekistan | 1930 | 010a7d50-a3e4-4775-b585-5ae6e36aef9c |
| ***Astragalus norvegicus*** | Moldenke | 21002 | NY | Sweden: Norrbotten: Kiruna | 1950 | d0b86805-bdcd-4822-aac9-352a90dc7723 |
| ***Astragalus nothoxys*** | van Devender | 97-387 | CAS | Mexico: Sonora: Yecora | 1997 | 16b6c824-dd61-4282-8f1d-5a953e1c4c3b |
| ***Astragalus noziensis*** | Volk | 71/380 | NY | Afghanistan: Paktia | 1971 | cd8c8704-786b-4f09-b0af-008e91761d04 |
| ***Astragalus nudisiliquus*** | Joyal | 503 | NY | United States of America: Oregon: Malheur | 1984 | d4f2d3d2-95b0-43f7-97e9-84201b4d343f |
| ***Astragalus nudus*** | Marticorena | 1465 | OS | Chile: Coquimbo (IV): Ovalle | 1971 | e7593fd3-960a-4bb9-a8bf-22c0c174dcf1 |
| ***Astragalus nutans*** | Thorne | 49,247 | NY | United States of America: California: San Bernardino | 1977 | d4e62e7d-12ed-4252-ad64-86708e613bd0 |
| ***Astragalus nutriosensis*** | Beasley | 948 | NY | United States of America: Arizona: Apache | 1991 | d4d7e070-6a01-415a-b769-1f6906a44ea4 |
| ***Astragalus nuttallianus*** | Reina | 2003-514 | CAS | Mexico: Sonora: Nacozari de Garc√≠a | 2003 | 16b54691-ad06-4fae-97f7-7428ec6c1610 |
| ***Astragalus nuttallii*** | Tilforth | 656 | NY | United States of America: California: Monterey | 1972 | d5d4301e-8248-487c-baa7-34c69cc24d20 |
| ***Astragalus nutzotinensis*** | Cody | 30320 | NY | Canada: Yukon Territory | 1982 | d4cc5b57-95d1-41c7-a489-708ec18f4f46 |
| ***Astragalus nyensis*** | Cochrane | 986 | NY | United States of America: Nevada: Nye | 1978 | d4c44052-6cc8-4c1b-8d2f-bef026cd86a1 |
| ***Astragalus obcordatus*** | Herring | 680 | FLAS | United States of America: Florida: Suwannee | 1992 | e912d82d-de22-4bce-b7b7-1f972fd4e983 |
| ***Astragalus obscurus*** | Foster | 11423 | NY | United States of America: Idaho: Owyhee | 1997 | d5d215ed-5590-48cc-b1c8-21eb6c216bd6 |
| ***Astragalus ochrochlorus*** | Ledingham | 4170 | NY | Islamic Republic of Iran | 1965 | cd8af527-5826-461e-b2ce-9a295c14c0af |
| ***Astragalus odoratus*** | USDA-ARS | 383593 | NY | Turkey: Erzurum: Yakutiye | 1977 | cd88d776-a602-421e-9a6f-6b2c800c4907 |

| | | | | | | |
|---|---|---|---|---|---|---|
| ***Astragalus olchonensis*** | Bannets | 337 | NY | Russian Federation: Siberia | 1979 | b0b91ceb-4f64-4df1-a482-9bc304581afd |
| ***Astragalus oldenburgii*** | B. Fedtsch | 6228 | HUH | Uzbekistan | 1930 | 0179a9e9-aa60-4409-a8eb-af6e864b3d8d |
| ***Astragalus oleaefolius*** | Manakyan | s.n. | MO | Armenia: Ararat | 1997 | 0c36fa66-6ae9-4f58-a459-92be36b09fd2 |
| ***Astragalus oleaefolius (Astracantha deinacantha)*** | Zohary | 344 | MO | Not shown | 1935 | 0c1718df-2cab-4a8c-a006-029bed2cf177 |
| ***Astragalus olgae*** | Vassiljeva | 5188 | NY | Uzbekistan | 1962 | cffba89a-873e-40fc-b09b-2cea19baa069 |
| ***Astragalus oniciformis*** | DeBolt | 1111 | NY | United States of America: Idaho: Lincoln | 1989 | d5375268-e432-463c-b970-233fca8cce66 |
| ***Astragalus onobrychioides*** | Tzvelev | 7135 | NY | Russian Federation: Daghestan: Gunib | 1961 | aea9e6cf-8e8c-4921-8eae-e01090c9d1b7 |
| ***Astragalus onobrychis*** | Krendl | s.n. | NY | Yugoslavia: Makedonija: Debar | 1976 | cf676976-7f9b-4dd7-b483-162e9f7be2f3 |
| ***Astragalus oocalycis*** | Atwood | 25653 | NY | United States of America: Colorado: La Plata | 2000 | d4b00b6a-94eb-48cb-b6a2-f01e1898748a |
| ***Astragalus oocarpus*** | Moran | 8340 | NY | United States of America: California: San Diego | 1960 | d4b5e78e-3f94-41d5-9bbb-13e0db13edd5 |
| ***Astragalus oophorus*** | Atwood | 15474 | FLAS | United States of America: Utah: Juab | 1991 | e991d56f-ca60-4dda-ad93-4cb0fb8d3cee |
| ***Astragalus orbiculatus*** | Hikitin | s.n. | HUH | Kazakhstan: Almaty | 1930 | 01716fe2-7974-48b4-b7c6-59591f895ac3 |
| ***Astragalus orcuttianus*** | Harbison | s.n. | CAS | Mexico: Baja California | 1951 | 16b14aea-25cb-42ff-b6c1-b8a4d673c60d |
| ***Astragalus oreganus*** | Lesica | 5369 | NY | United States of America: Montana: Carbon | 1991 | d530324e-efc5-4df9-9565-3c9279684a1e |
| ***Astragalus ornithopodioides*** | Mona | s.n. | NY | Armenia: Yerevan | 1960 | aea878f9-b9d0-4c55-b267-e49077195ddb |
| ***Astragalus ornithorrhynchus*** | Goloskokov | 4418 | NY | Kazakhstan | 1947 | aea0a78c-b9af-4f48-80b9-90ed5d312d5f |
| ***Astragalus osterhoutii*** | Neese | 17181 | NY | United States of America: Colorado: Grand | 1985 | d528fb48-0ef5-4608-a258-04b0054217fb |
| ***Astragalus oxyglottis*** | Lachashvili | 447 | NY | Georgia: East Georgia | 2006 | d0aaf4d9-1447-468e-9666-70597a727aad |
| ***Astragalus oxyphysopsis*** | Moran | 23095 | TEX | Mexico: Baja California | 1976 | 02551a79-54b1-4005-8832-fd3072e967fc |
| ***Astragalus oxyphysus*** | Moran | 28353 | CAS | Mexico: Baja California: Ensenada | 1980 | 16b00c1e-46f1-4d27-8c09-88146b41712f |
| ***Astragalus pachypus*** | Mistretta | 1521 | NY | United States of America: California: Los Angeles | 1995 | d523607c-3ac0-4b76-87c4-d6c1d860b2f6 |
| ***Astragalus pachyrachis*** | Volk | 20 | NY | Afghanistan: Kabul | 1950 | aad5e993-2f37-41bb-8194-037be123645f |
| ***Astragalus palenae*** | O'Donell | 2384 | NY | Argentina: Neuquen | 1945 | d13a562a-61ec-4a09-a33d-10a30907169f |
| ***Astragalus palmeri*** | Beauchamp | 2036 | FLAS | United States of America: California: San Diego | 1971 | e97ff386-dd07-4bd5-9b33-be43948ed26c |

| | | | | | | |
|---|---|---|---|---|---|---|
| ***Astragalus pamirensis*** | Stanjukovicz | 6266 | HUH | Tajikistan | 1936 | 0154a316-124c-4a8e-880d-587916a21adf |
| ***Astragalus panamintensis*** | Niles | 5591 | NY | United States of America: Nevada: Nye | 1998 | d4ae36d6-1fd4-41de-82c2-8ca8243bf882 |
| ***Astragalus paposanus*** | Jiles | 5394 | OS | Chile: Antofagasta (II): Antofagasta | 1969 | e73deb1b-ac93-4197-ab05-4992669943b7 |
| ***Astragalus pardalinus*** | Welsh | 9592 | NY | United States of America: Utah: Wayne | 1970 | d519a02a-0105-41b4-b013-6a5d325a6ffb |
| ***Astragalus parnassi*** | de Fellenberg | s.n. | NA | Greece: Attika | 1863 | e7278a3e-5f74-4c97-a89e-787be6aed369 |
| ***Astragalus parodii*** | Sayago | 815 | NY | Argentina: Cordoba: El Batan | 1952 | d13a1878-3a05-4fa7-850e-ee7750dfcd37 |
| ***Astragalus parryi*** | Isely | 9800 | NY | United States of America: Wyoming: Albany | 1965 | d515f796-ac19-478f-baea-03f8a0afcee3 |
| ***Astragalus parvus*** | McVaugh | 16982 | CAS | Mexico: Jalisco | 1958 | 14d7ec68-3ed3-4662-932e-fcee2f367d6e |
| ***Astragalus patagonicus*** | [None] | | NY | Argentina: Santa Cruz: Lago Buenos Aires | 2017 | d1377632-f24c-4d9c-a28c-f1f039451694 |
| ***Astragalus pattersonii*** | Patterson | | FLAS | United States of America: Colorado | 1936 | e97b1679-e99b-4404-b1ef-0e1fe20a6657 |
| ***Astragalus paucijugus*** | Rusanovich | | NY | Kazakhstan: Alma-Ata: Taukum Sands | 1984 | d0a269dd-8ec9-42d7-ad48-34c0daca0eb7 |
| ***Astragalus pauperculus*** | Barneby | 11491 | NY | United States of America: California: Tehama | 1954 | d4a56123-a80e-42ce-b6ec-3477e62d98f8 |
| ***Astragalus pavlovii*** | [Cyrillic] | 6589 | NY | Mongolia | 1948 | ae96d7e2-d564-40ce-8c48-255038e61535 |
| ***Astragalus paysonii*** | Holmgren | 16590 | NY | United States of America: Wyoming: Sublette | 1978 | d49cf7c1-cad9-4e30-bc37-6c8ec00f809b |
| ***Astragalus peckii*** | Peck | 21931 | NY | United States of America: Oregon: Klamath | 1943 | d4948adc-548c-4811-acb5-0ab3a4542288 |
| ***Astragalus pectinatus*** | Moran | 426 | FLAS | United States of America: North Dakota: Billings | 1939 | e9410742-2815-4251-9a03-48b86c555896 |
| ***Astragalus peduncularis*** | Anders | 7232 | HUH | Afghanistan: Badakhshan | 1971 | f7bc2efa-1512-46c1-8104-d2de8c869a4b |
| ***Astragalus peduncularis (Astragalus corydalinus)*** | Butschantzev V; Butkov. A | 1936 VII 7 | HUH | Russian Federation | 1936 | ff9f67e8-5008-4118-a690-878438e76436 |
| ***Astragalus pehuenches*** | Marticorena | 972 | NA | Chile: Curico: Curico | 1967 | e748e253-55f5-4401-b7ad-2196311396f7 |
| ***Astragalus pelecinus*** | Raus | 9257 | NY | Greece | 1984 | d0a210d0-6de7-4ce4-ae19-cbea9aad8098 |
| ***Astragalus pennellianus*** | Breedlove | 63019 | CAS | Mexico: Durango | 1986 | 167e5316-d31f-43ff-b128-f7cba467f996 |
| ***Astragalus peregrinus*** | Tackholm | s.n. | MO | Egypt | 1965 | 164eb72e-0e07-4030-919b-8dceb6871476 |
| ***Astragalus perianus*** | Madsen | 1484 | NY | United States of America: Utah: Piute | 2002 | d472304a-6fde-46ae-9665-cea46d031ae9 |

| | | | | | | |
|---|---|---|---|---|---|---|
| ***Astragalus persepolitanus*** | Volk | 1690 | NY | Afghanistan: Kabul | 1950 | aad5639b-4bd4-414c-ae17-552349223f70 |
| ***Astragalus peruvianus*** | Casas | FC 66 40 | NY | Bolivia (Plurinational State of): Murillo: La Paz | 1982 | d10359d1-bc7e-4430-81a5-e975698fad86 |
| ***Astragalus peterfii*** | Peterfi | s.n. | NY | Romania | 1916 | cf683a0b-10b0-465f-944f-4077d597f32e |
| ***Astragalus petropylensis*** | Schischkin | 3323 | NY | Kazakhstan: East Kazakhstan: Oskemen | 1931 | ae8219b6-3eae-4c49-a29e-300b37b99cd0 |
| ***Astragalus phoenix*** | Beatley | none | NY | United States of America | 1970 | d50d564a-7ab5-4b9d-8322-118c420fe380 |
| ***Astragalus physocarpus*** | USDA-ARS | 420687 | NY | Kazakhstan | 1980 | ae7886ba-1ded-4851-9986-bef679e5f998 |
| ***Astragalus physodes*** | Skvortsov | | NY | Russian Federation: Volgograd | 1968 | cf6c7976-bab5-41ec-a0ff-8e4d9a7df04b |
| ***Astragalus pickeringii*** | Weigend | 5471 | NY | Peru | 2001 | d0fce35d-b817-4e89-9954-28c3bd247218 |
| ***Astragalus pictiformis*** | Barneby | 18076 | NY | United States of America: New Mexico: Lincoln | 1986 | d480d94b-b78f-4b11-ae1f-1125e4fdc40d |
| ***Astragalus piletocladus*** | [Cyrillic] | s.n. | NY | Turkmenistan | 1954 | ae7f9296-3dab-4dd4-af6d-6416ff99e125 |
| ***Astragalus pinetorum (Tragacantha declinata)*** | [Cyrillic] | s.n. | MO | Armenia | 1962 | 0c69abbc-8c6c-4594-8b8b-51dbf4d6d45a |
| ***Astragalus pinetorum*** | Samuelsson | 2256 | NY | Lebanon | 1932 | aad0a9ce-b2ea-4d61-857e-d7b0fb40b5bf |
| ***Astragalus pinonis*** | Atwood | 20824 | NY | United States of America: Arizona: Coconino | 1996 | d49278c5-6412-4357-83f9-84b8e7712085 |
| ***Astragalus piscator*** | Barneby | 17808 | NY | United States of America: Utah: Grand | 1982 | d4716456-ac22-4429-b0e8-c15ba3285df5 |
| ***Astragalus piscinus*** | Raven | 12564 | CAS | Mexico: Baja California: Ensenada | 1958 | 16ab185c-516b-4211-bdfa-5e21dbc8000b |
| ***Astragalus pissisi*** | Ricardi | | NA | Chile: Coquimbo (IV): Choapa | 1952 | e742cecb-e006-4908-940a-8d441ca57f70 |
| ***Astragalus piutensis*** | Tiehm | 9474 | NY | United States of America: Nevada: White Pine | 1985 | d4912524-49ed-4d20-ab17-d3ea2e0c255f |
| ***Astragalus plattensis*** | Wheeless | 15 | FLAS | United States of America: Texas: Eastland | 1966 | e93fce93-959a-4f20-9888-a7b05ae2a7b7 |
| ***Astragalus platyphyllus*** | Ishmayaova | 1746 | NY | Kazakhstan | 1970 | ae7c2716-4bb9-4cdd-baec-20db4293e093 |
| ***Astragalus platysematus*** | USDA-ARS | 380721 | NY | Islamic Republic of Iran: Zadabeh: Unknown | 1979 | aacb0391-632f-4ff3-8015-0a7111fc6145 |
| ***Astragalus platytropis*** | Packard | 79-215 | NY | United States of America: Oregon: Malheur | 1979 | d481b244-eba8-4935-8847-2582332b85f9 |
| ***Astragalus plumosus*** | Lambert | 621 | NY | Turkey | 1968 | aac3f097-1c40-41fd-8b36-4da68b04ff91 |
| ***Astragalus podolobus*** | Ladingham | 4098 | NY | Islamic Republic of Iran | 1965 | aac334fe-1694-4d82-b756-e64214762a2d |

| | | | | | | |
|---|---|---|---|---|---|---|
| ***Astragalus polaris*** | Astragalus Polaris Benth | 535 | NY | United States of America: Not shown | 1975 | ae747978-4d46-49be-86a6-73278be85e08 |
| ***Astragalus polyacanthus*** | Webster | 6087 | HUH | India: Kashmir | 1955 | f7b32493-d705-4ccb-ae6c-4dda36ae533d |
| ***Astragalus polybotrys*** | Ledingham | 5277 | NY | Canada: Saskatchewan | 1970 | aa9d2e93-f052-4e05-80ed-a58967643530 |
| ***Astragalus pomonensis*** | Howell | 31085 | CAS | Mexico: Baja California | 1956 | 16a96aa9-20c4-4883-b8b5-243ca4f7ce2d |
| ***Astragalus ponticus*** | Akinfiev | 1800 | NY | Ukraine: Dnepropetrovsk | 1906 | cf6daeed-e72e-4b99-a49c-814c76f7b030 |
| ***Astragalus porrectus*** | Tiehm | 4281 | NY | United States of America: Nevada: Pershing | 1978 | d46fdec5-4343-422d-8371-795cd7a54d03 |
| ***Astragalus praelongus*** | Atwood | 23622 | NY | United States of America: Colorado: San Miguel | 1998 | d56eee88-264d-4b09-a656-fa2d2ae151fa |
| ***Astragalus preussii*** | Neese | 16778 | NY | United States of America: Arizona: Coconino | 1985 | d462e850-8ed1-42f3-ba67-154e98617503 |
| ***Astragalus prorifer*** | Boyd | 2793 | NA | Mexico: Baja California: Sierra San Pedro M√°rtir | 1988 | 16a95a1b-69e5-4ad9-b064-fc262b422314 |
| ***Astragalus proximus*** | Barneby | 17795 | NY | United States of America: New Mexico: San Juan | 1982 | d45babaf-8762-4492-b7e3-235353c14feb |
| ***Astragalus przewalskii*** | Boufford | 36686 | HUH | People’s Republic of China: Sichuan | 2006 | 009cf286-56e7-4f0b-b595-897d50127858 |
| ***Astragalus pseudoadsurgens*** | Maltzev | 7333 | HUH | Russian Federation: Irkutsk: Ust-Uda | 1964 | 00699996-f908-4284-96a2-b5db4c644b88 |
| ***Astragalus pseudocyclophyllus*** | Rechinger | 309 | NY | Russian Federation: Stavropol Krai | 1949 | aac03dcf-f334-4eda-a3fe-1a6e66b52972 |
| ***Astragalus pseudocytisoides*** | Bajtenov | 5378 | NY | Kazakhstan | 1967 | ae62f62f-0856-4408-8217-9f2352de1723 |
| ***Astragalus pseudomegalomerus*** | Botschantzev | | HUH | Uzbekistan | 1936 | 00f724b2-4f02-40e3-b59d-5c3cfa333c6e |
| ***Astragalus pseudonobilis*** | Rajkova | 6232a | HUH | Uzbekistan | 1926 | 00efe10e-c18e-4341-aba8-9df3cb985fc0 |
| ***Astragalus pseudoparrowianus*** | Rechinger fil. | 988 | NY | Islamic Republic of Iran | 1937 | aab0393a-37ed-443a-9325-0a030efacf80 |
| ***Astragalus pseudoutriger*** | Petunnikovy | 3720 | NY | Azerbaijan | 1910 | cf7109f5-8bb3-4a8e-a639-fa8d91b979a8 |
| ***Astragalus psilacanthus*** | Lamond | 2442 | NY | Afghanistan: Paktia | 1965 | aaa230d9-87ce-41d4-b197-7e8e1476aad2 |
| ***Astragalus psilocentros*** | Stewart | 23607 | NY | Pakistan: Punjab | 1949 | aaa96a17-b7c5-459b-9844-06b0df2e9e05 |
| ***Astragalus pterocarpus*** | Tiehm | 16050 | NY | United States of America: Nevada: Humboldt | 2010 | d56ee106-5c36-4d5c-86a0-03dbab666e6a |
| ***Astragalus pterocephalus*** | Belolipov | UPL_00396 | MO | Uzbekistan: Djizak: Bahmal | 2005 | 0c5ccf8d-890b-490c-8201-6db4ce9247b7 |
| ***Astragalus pubentissimus*** | Goodrich | 1473 | NY | United States of America: Wyoming: Sweetwater | 1973 | d459b698-c212-47fd-bcf7-895b729ce9dc |
| ***Astragalus puberulus*** | Cronquist | 12151 | NY | Russian Federation: Altai | 1988 | ae616acd-e14f-4feb-aa5b-d1a41cb117c9 |

| | | | | | | |
|---|---|---|---|---|---|---|
| ***Astragalus pullus*** | Boufford | 28432 | HUH | People's Republic of China | 1998 | 00d5c748-f651-496e-9a60-fb578922e0c0 |
| ***Astragalus pulsiferae*** | Schoolcraft | 385 | NY | United States of America: California: Modoc | 1981 | d56e81c0-2d1b-4c97-8867-50d4bb0cc9e5 |
| ***Astragalus punae*** | M√∫lgura | 1272 | NY | Argentina: Jujuy: Susques | 1994 | d1301124-9cf5-44e9-a442-aff58e5064b4 |
| ***Astragalus punctatus*** | Grossheim | 233 | NY | Islamic Republic of Iran: East Azerbaijan | 1924 | af6b4283-fda8-4413-86cd-720d974371d7 |
| ***Astragalus puniceus*** | Spellenberg | 4976 | NY | United States of America: New Mexico: Taos | 1978 | d56367f1-f3aa-4493-b3f8-535cce698214 |
| ***Astragalus purshii*** | Neely | 4190 | NY | United States of America: Colorado: Moffat | 1987 | d454bd53-46e5-47cd-a88a-113b25888dfb |
| ***Astragalus pusillus*** | Solomon | 16211 | NY | Bolivia (Plurinational State of) | 1987 | d12c1297-1b05-4d53-84c9-60c5b2128da5 |
| ***Astragalus pycnostachyus*** | Constance | 2523 | NY | United States of America: California: Marin | 1939 | d5600e08-7de0-45aa-867e-00237bfb2704 |
| ***Astragalus quisqualis*** | [Cyrillic] | | HUH | [Cyrillic] | 1965 | 009612ca-8605-4c41-bbc0-71b7ff8b5e01 |
| ***Astragalus racemosus*** | Goodrich | 23331 | FLAS | United States of America: Utah: Uintah | 1991 | e93b0012-c2e3-4418-b16b-220d4d9f4989 |
| ***Astragalus raddei*** | Litvinov | 1231 | NY | Turkmenistan | 1898 | af584e6d-2926-4e34-877b-9af3a2fbd6d7 |
| ***Astragalus rafaelensis*** | Shultz | 2502 | NY | United States of America: Utah: Emery | 1978 | d44d808b-e9b9-440a-a8ac-5488082f5a07 |
| ***Astragalus rattanii*** | Halse | 8764 | NY | United States of America: California: Colusa | 2013 | d4415fe8-5dc9-44e4-bfb1-dd76eb4f4bb5 |
| ***Astragalus recurvus*** | Imdorf | 1221 | NY | United States of America: Arizona: Gila | 1993 | d44107cf-c00d-4d50-86eb-3120e1ee1ed0 |
| ***Astragalus reduncus*** | Sagalaev | s.n. | NY | Russian Federation: Volgograd: Svetloyarsky | 1993 | cf81a944-c83f-4743-9b66-01a1e62873b2 |
| ***Astragalus reflexistipulus*** | Yokoyama | 9872501 | HUH | Japan: Yamagata Prefecture | 1998 | 00951840-8ffd-434b-84f8-60919cabf9e8 |
| ***Astragalus reflexus*** | Simon E. Wolff | 2877 | NY | United States of America: Texas: Bell | 1931 | d431af25-613e-4c87-b124-119805c1d21a |
| ***Astragalus refractus*** | USDA-ARS | 384801 | NY | Islamic Republic of Iran | 1977 | aa9a5ff9-f58f-4d3c-bde6-f012aec27350 |
| ***Astragalus remotiflorus*** | Assadi | 1775 | NY | Islamic Republic of Iran: Kerman: Baft | 1977 | af54ec47-2d86-4845-b506-4e957535bb19 |
| ***Astragalus remotus*** | Pinzl | 11228 | NY | United States of America: Nevada: Clark | 1995 | d55d205c-d416-4add-9625-51b2aec9c758 |
| ***Astragalus retamocarpus*** | Michelson | 3324 | NY | Uzbekistan: Samarqand | 1913 | af52aabf-ee34-45ad-ada2-87bbf6406c20 |
| ***Astragalus reticulatus*** | Roshewitz | 382 | NY | Kazakhstan | 1926 | b02e67cb-801d-46c9-a897-d969cb4b513c |

| | | | | | | |
|---|---|---|---|---|---|---|
| ***Astragalus reuterianus*** | Assadi | 1698 | NY | Islamic Republic of Iran: Kerman: Baft | 1977 | aa94d172-4003-4921-9de4-e43c5ccc2dd8 |
| ***Astragalus reventiformis*** | Chamberlain | 1265765 | NY | United States of America: Washington: Klickitat | 1980 | d555e9b8-e953-487a-8471-b1d72eb76986 |
| ***Astragalus reventus*** | Larry Hufford | 1192 | NY | United States of America: Washington: Garfield | 1996 | d5522bbf-cfb3-4ac3-8b1d-57a9da056e14 |
| ***Astragalus rhacodes*** | [Cyrillic] | 714 | HUH | [Cyrillic] | 1960 | 0090d966-d9fd-4fbb-a015-1b0a4551c308 |
| ***Astragalus rhizanthus*** | Webster | 6349 | HUH | Pakistan: Kashmir: Baltistan | 1955 | fc40d59f-a4f6-449f-a612-3c1dea13c5a9 |
| ***Astragalus rhizocephalus*** | Volk | 2752 | NY | Afghanistan: Bamian | 1952 | cd7662de-73dd-445b-99b6-c5ba3090e234 |
| ***Astragalus richii*** | Leiva | 1155 | NY | Peru: Otuzco: La Libertad | 1994 | d23bbac6-3363-4720-bd6d-7a4b7283c873 |
| ***Astragalus ripleyi*** | Weber | 7788 | FLAS | United States of America: Colorado: Conejos | 1952 | e9382f96-2850-4701-9840-4ddc382107e6 |
| ***Astragalus robbinsii*** | Kierstead | 84-28 | NY | United States of America: Oregon: Wallowa | 1984 | d42d5002-4f63-493f-a3b2-665975825fe3 |
| ***Astragalus robustus*** | Takhtajan | s.n. | NY | Armenia | 1960 | b02a95bd-3b63-4689-8cf4-0bfed6019b42 |
| ***Astragalus roemeri*** | Ledingham | 5264 | NY | Czech Republic | 1970 | cf9112f3-27e4-4ff3-9db7-a409dd6970d0 |
| ***Astragalus romasanus*** | Hoffmann | 170 | TEX | Peru | 1960 | 20cd213b-095d-4843-9825-e0fd0edebcac |
| ***Astragalus roseus*** | Schischkin | s.n. | NY | Kazakhstan: East Kazakhstan: Oskemen | 1931 | b02a8e16-8144-426e-bda2-8f2da88b5065 |
| ***Astragalus rubrifolius*** | Butkov | 6236 | NY | Turkmenistan | 1930 | b0236ce0-2754-4daa-95e3-976eb1e2848a |
| ***Astragalus rubromarginatus*** | Korovin | 365 | NY | Turkmenistan | 1925 | b0295514-a83a-482e-ad4b-488299548448 |
| ***Astragalus rubtzovii*** | Aristangaliev | 4820 | HUH | Kazakhstan | 1963 | 008a54c8-4051-4b77-afd2-e0be2519a7c4 |
| ***Astragalus ruiz-lealii*** | Covas | 3059 | NY | Argentina: Mendoza: Las Heras | 1944 | d23b0c40-f18b-4741-9b43-75d12b8bab09 |
| ***Astragalus rupifragus*** | Skvortsov | s.n. | NY | Russian Federation: Saratov: Hvalynsk | 1993 | cf777972-6ed2-4d13-9230-608f79bee691 |
| ***Astragalus rusbyi*** | Isely | 10815 | NY | United States of America: Arizona: Coconino | 1969 | d5511611-0bc2-443b-8316-b927020c1c62 |
| ***Astragalus ruscifolius*** | USDA-ARS | 384744 | NY | Islamic Republic of Iran | 1977 | ce7a63c7-fcef-452e-8d4d-88008d47a972 |
| ***Astragalus sabulonum*** | Felger | 92-221 | CAS | Mexico: Sonora: San Luis R√≠o Colorado | 1992 | 169fa003-281c-4e59-a836-5e949faf7a52 |
| ***Astragalus sabulosus*** | Welsh | 4021 | NY | United States of America: Utah: Grand | 1965 | d42b9f80-a4b0-479d-be76-5748c6e8cc40 |

| | | | | | | |
|---|---|---|---|---|---|---|
| ***Astragalus sachalinensis*** | Pavlova | 5970 | NY | Russian Federation: Sakhalin: Alexandrovsk-Sakhalinsky | 1978 | ae121256-56f2-4840-971d-dce335482eaf |
| ***Astragalus salmonis*** | Tiehm | 12852 | NY | United States of America: Nevada | 1999 | d42b6086-494b-4589-b7bd-b55738d62f70 |
| ***Astragalus sanctae-crucis*** | Hicken | SI 7524 | NY | Argentina: Buenos Aires: Partido del General Pueyrred√≥n | 1932 | d0e07e87-1ce8-4895-97fa-92a0733768ce |
| ***Astragalus sanctorum*** | Moran | 15929 | TEX | Mexico: Baja California | 1969 | 20e9ca4f-a182-4adf-919a-eb6c671e22b5 |
| ***Astragalus sanguineus*** | Thompson | 524 | CAS | Mexico: Coahuila | 1983 | 1643c244-1682-4956-8d54-f4f909fe9348 |
| ***Astragalus saurinus*** | Atwood | 31216 | NY | United States of America: Utah: Uintah | 2005 | d425846d-ef0b-425b-acf5-8a59b74d2c02 |
| ***Astragalus saxifractor*** | Podlech | 17775 | NY | Afghanistan: Kabul | 1970 | ce79fde7-d587-4ab7-b4f8-73437f5090cc |
| ***Astragalus scaberrimus*** | Guo | 1504049 | HUH | People’s Republic of China: Shandong: Qufu | 2004 | 006b049a-8de8-4881-89e0-dc7db5ff6e1a |
| ***Astragalus scaphoides*** | Henderson | 3733 | NY | United States of America: Idaho | 1977 | d40bee28-5b2a-4bf7-a3d0-7eed3cc6ff6c |
| ***Astragalus schachdarinus*** | Anders | 7233 | HUH | Afghanistan: Badakhshan | 1971 | f7b0f5c1-92c0-4202-8b93-ab6d4f355300 |
| ***Astragalus schahrudensis*** | Nikitin | | NY | Turkmenistan: Ashgabat | 1975 | ae0e2c48-ca97-4d4f-b735-56544160a1f6 |
| ***Astragalus schanginianus*** | Goloskokov | 4419 | NY | Kazakhstan | 1959 | ae0d4401-a3df-41f8-b9fe-40a3b42143e4 |
| ***Astragalus scheremetevianus*** | Gusev | 4580 | NY | Tajikistan: Gorno-Badakhshan | 1958 | adf422ec-8258-4393-9af0-16cc88bfd37b |
| ***Astragalus schimperi*** | Barkley | 33Ir5304 | NY | Iraq: Al Anbar | 1963 | ce752161-59c3-4d36-92d3-4e4a2c3f9e71 |
| ***Astragalus schistocalyx*** | Assadi | 2131 | NY | Islamic Republic of Iran: Kerman | 1977 | d54e75e4-19b3-47f5-bd4e-38967e67e65e |
| ***Astragalus schmalhausenii*** | Vaisk | 152 | NY | People’s Republic of China | 1989 | ade600df-120f-424f-b8d1-e1f5358b443b |
| ***Astragalus schrenkianus*** | Bajtenov | 5773 | HUH | Kazakhstan | 1963 | 00e462f3-4037-46fe-9f65-f7eda14bb6ad |
| ***Astragalus schugnanicus*** | [Cyrillic] | 188 | NY | [Cyrillic] | 1959 | ad288896-1b64-4f52-b51f-4dce1f511147 |
| ***Astragalus scleroxylon*** | Botschantzev | 6238a | HUH | Uzbekistan: Bukhara | 1937 | 009cca2c-e074-400b-970b-284a5f5e6762 |
| ***Astragalus scoparius*** | Goloskokov | 4420 | NY | Kazakhstan: Almaty | 1959 | ad272d51-3435-4317-8de6-b7eec7abf0f7 |
| ***Astragalus scopulorum*** | Franklin MA | 6530 | NY | United States of America: Utah: Utah | 1989 | d3f44e6b-d7d5-4358-9d5d-dbd2c69fba34 |
| ***Astragalus scorpiurus*** | Stewart | 23,604 | NY | Pakistan: Punjab: Attock | 1949 | aaa90af4-98bc-4ed1-9c37-d9294afe0642 |
| ***Astragalus scutaneus*** | Holmes | CES-338 | TEX | Mexico: Jalisco: Chapala | 1998 | 20e57763-2dab-418e-828e-07261f4dfdab |
| ***Astragalus semenovii*** | Goloskokov | 4128 | NY | Kazakhstan | 1955 | ad1e4201-94a2-4763-94ad-4cc6cd196540 |
| ***Astragalus sempervirens*** | Echevarria | MAF 154466 | TEX | Spain: Madrid | 1996 | 20b84e9b-29d1-431d-bbd9-038eb9e32d6b |

| | | | | | | |
|---|---|---|---|---|---|---|
| ***Astragalus sepultipes*** | DeDecker | 5930 | NY | United States of America: California: Inyo | 1987 | d3eb5c09-0672-4ce2-adb2-3d39a5a2f2d2 |
| ***Astragalus serenoi*** | Tiehm | 14965 | NY | United States of America: Nevada: Churchill | 2005 | d41a5915-c0dc-4b8a-a6eb-3fe52e9a8283 |
| ***Astragalus serpens*** | Atwood | 11796 | NY | United States of America: Utah: Wayne | 1985 | d3e2f51c-2296-4afa-b5b5-f386fd60c83f |
| ***Astragalus sesameus*** | Miller | 283 | MO | Morocco: SW | 1974 | 163fb5e0-6d45-4ef9-9f21-f81b6d260c14 |
| ***Astragalus sesamoides*** | Goloskokov | 4471 | NY | Kazakhstan: Zhambyl | 1963 | ad15d5cb-ef03-432c-8c62-45be3a8000dc |
| ***Astragalus sesquiflorus*** | Holmgren | 10,641 | NY | United States of America: Utah: San Juan | 1954 | d3d0638c-6538-483c-9071-2227281dbc9b |
| ***Astragalus setulosus*** | USDA-ARS | 383597 | NY | Turkey: Nidgen Pernenial: Forty km | | aa63dc51-4ae8-4854-bf9f-1850fedb03b3 |
| ***Astragalus sevangensis*** | Sytin | 6535 | NY | Armenia: Gegharkunik | 1981 | d0914b68-17ed-4423-80ba-cc0158701cbc |
| ***Astragalus sewertzovii*** | Czerepanov | 7068 | NY | Tajikistan: Sughd | 1965 | ad0e8c1d-be9c-4f01-b408-2c53d047d4d2 |
| ***Astragalus shagalensis*** | Sytin | | NY | Armenia: Lori | 1981 | d08841a1-9a7b-4049-b7af-833364f9e830 |
| ***Astragalus sheldonii*** | Gray | 5221 | NY | United States of America: Idaho: Nez Perce | 2005 | d37c2128-e540-4592-a3b8-5cda78b09253 |
| ***Astragalus shevockii*** | Shevock | 5633 | NY | United States of America: California: Tulare | 1977 | d3784f3a-cc4e-411a-bcf0-2718c0f225fd |
| ***Astragalus shinanensis*** | Furuse | s.n. | HUH | Japan: Shinano: Nagano | 1962 | 007d2455-ddce-484e-9486-448a49a2817d |
| ***Astragalus shortianus*** | Elliott | 6701 | NY | United States of America: Colorado: Chaffee | 1999 | d3ccd8a6-7454-459e-8255-8c3db8fabf93 |
| ***Astragalus shultziorum*** | Lichvar | 3926 | NY | United States of America: Wyoming: Lincoln | 1980 | d2e3f7c3-57ed-49e1-ba84-a8fece70c9c0 |
| ***Astragalus siahderrensis*** | Lamond | 2421 | NY | Afghanistan: Paktia: Gardez | 1965 | aa49d71c-3f10-49ca-9802-ced5c79bde5b |
| ***Astragalus sieberi*** | Ledingham | 3469 | NY | Canada: Saskatchewan | 1962 | d03499f1-d5f5-4e1a-bcd6-e8cfc07abed7 |
| ***Astragalus sieversianus*** | [Cyrillic] | s.n. | NY | Russian Federation | 1975 | acf49d98-cee8-4676-b922-6689f76e1a70 |
| ***Astragalus sikokianus*** | Kim | | NY | Korea (Republic of) | 2003 | acefc368-d4b2-4700-a1d3-c587c36d8188 |
| ***Astragalus siliceus*** | Barneby | 17870 | NY | United States of America: New Mexico: Torrance | 1983 | d3bd1d1a-39c1-433b-bbd0-aee50ed159d7 |
| ***Astragalus siliquosus*** | Bleak | 330696 | NY | Islamic Republic of IranMalayer | 1974 | aa5e2172-b180-434e-8cbf-2fed9fe3c916 |
| ***Astragalus simplicifolius*** | Lichvar | 2722 | NY | United States of America: Wyoming: Fremont | 1980 | d37167cf-2f63-4856-91c7-99856c8bcf30 |
| ***Astragalus sinicus*** | Sakiya | 42 | NY | Japan | 2004 | ace8aa15-d0ba-48d1-8968-7f69f8f909b4 |

| | | | | | | |
|---|---|---|---|---|---|---|
| ***Astragalus sinuatus*** | Spellenberg | 1651 | NY | United States of America: Washington: Chelan | 1967 | d2f5d646-68a3-48ae-ba0a-a0cac6fd7d83 |
| ***Astragalus skorniakowi*** | Ledingham | 3676 | NY | Canada: Saskatchewan | 1964 | acdebd89-6212-49f6-9914-b5a19e7c1c83 |
| ***Astragalus skythropos*** | Boufford | 26916 | NY | People's Republic of China: Qinghai: Chindu Xian | 1995 | acd8b177-7dbc-4999-865b-4dfcfa0bac91 |
| ***Astragalus solandri*** | Podlech | 53637 | MO | Morocco: Tiznit | 1997 | 165474bb-5e1e-455f-9980-e271e8d7fa7b |
| ***Astragalus solitarius*** | Packard | 79-35 | NY | United States of America: Oregon: Malheur | 1979 | d36b240f-a374-4c2d-a1e1-6d1734b53463 |
| ***Astragalus somcheticus (Astragalus demetrii)*** | Bleak | 314060-061 | NY | Russian Federation: Stavropol Krai | 1974 | cfd35c2e-0999-4944-b919-9325d7817bca |
| ***Astragalus somcheticus (Astragalus polygala)*** | Cuba | s.n. | NY | Russian Federation: Caucasus | 1980 | ae66b316-0f60-4a60-9544-e8ff826b3a2c |
| ***Astragalus sophoroides*** | Atwood | 25368 | NY | United States of America | 2000 | d35f9e39-5bc4-46ad-8aff-eb393db037f3 |
| ***Astragalus soxmaniorum*** | Givens | 3564 | FLAS | United States of America: Louisiana: Caddo | 1984 | ea7d2782-d7da-46bb-947b-32b07aeb1b08 |
| ***Astragalus spachianus*** | Barneby | 184 | NY | Islamic Republic of Iran | 1977 | aa5ca764-38ae-41dd-b384-7a4dbf08374b |
| ***Astragalus spaldingii*** | Joyal | 1173 | NY | United States of America: Oregon: Baker | 1986 | d2f5892b-61f8-400c-872d-b12ee87f6adf |
| ***Astragalus sparsiflorus*** | Weber | 13294 | NY | United States of America: Colorado: El Paso | 1967 | d34b2023-2f4b-4b36-b949-605565340213 |
| ***Astragalus spatulatus*** | Isely | 9753 | NY | United States of America: North Dakota: Golden Valley | 1965 | d2eca01b-c4e0-46e5-8c89-ef47a331fc32 |
| ***Astragalus speirocarpus*** | Hufford | 2716 | NY | United States of America: Washington: Kittitas | 1998 | d2eaf5c9-79de-4d7e-9962-2ca139f560c7 |
| ***Astragalus spinosus*** | Fagerstrom | 220 | NY | Saudi Arabia: Riyadh: Sadus | 1979 | cd94670c-aaf0-47e3-9857-76d168972455 |
| ***Astragalus sprucei*** | Iltis | E-559 | NY | Ecuador: Chimborazo | 1977 | d1e17c85-fe5d-4277-abf3-383edc090048 |
| ***Astragalus squarrosus*** | Popov | 6239 | HUH | Turkmenistan: Balkan | 1931 | 00702fb4-6294-4022-8f95-cbe2e3b54f3d |
| ***Astragalus stalinskyi*** | Rasulova | 350 | NY | Tajikistan: Sughd | 1955 | ad7a375c-292f-413a-9939-81a7961b012d |
| ***Astragalus stella*** | Miller | 549 | MO | Morocco: SW | 1974 | 16412939-c849-451d-b421-7789ecf78d61 |
| ***Astragalus stevenianus*** | Lachashvili | 352 | NY | Georgia: Kakheti: Dedoplistskaro | 2006 | d0692ca5-ff85-4d2f-aebe-6be607de0233 |
| ***Astragalus stewartii*** | Stewart | 22631 | NY | India: Kashmir | 1946 | aafe44f8-b3bf-4254-9ed0-be83754d0277 |
| ***Astragalus stipulatus*** | Ohashi | 771423 | HUH | Nepal | 1977 | f7d15a5b-1c52-4916-bb5e-a1015eaf5b94 |
| ***Astragalus stocksii*** | Anders | 8792 | NY | Afghanistan: Kandahar | 1972 | aa8b360f-1489-40ee-b8c9-d6215ab20090 |
| ***Astragalus straturensis*** | Atwood | 4909 | NY | United States of America: Utah: Washington | 1973 | d342979e-417f-46ad-9f41-18b837920cd4 |

| | | | | | | |
|---|---|---|---|---|---|---|
| ***Astragalus striatiflorus*** | Atwood | 23915 | NY | United States of America: Utah: Kane | 1998 | d2e3de91-437d-4175-b109-2506fc511e7e |
| ***Astragalus strictus*** | Leta Kharka | 20105042 | HUH | Nepal: Dhawalagiri: Mustang | 2001 | fc6c2e41-4d5f-4e19-a071-972af95bdb07 |
| ***Astragalus subcinereus*** | Higgins | 26026 | NY | United States of America: Arizona: Coconino | 2004 | d2dba836-f8f3-4b3c-898e-a179d79ef4bc |
| ***Astragalus suberosus*** | Samuelsson | 3677 | NY | Syrian Arab Republic | 1933 | a9dc964b-4ec8-4bfb-ba3f-13f70c73c01e |
| ***Astragalus submitis*** | Ledingham | 4058 | NY | Islamic Republic of Iran: Tehran | 1965 | a9ef9b33-adac-4d1e-b405-f60b296aa686 |
| ***Astragalus substipitatus*** | Vasak | s.n. | NY | Tajikistan | 1900 | ad78b8d1-5916-4d42-933a-5cd5ccd7cfed |
| ***Astragalus subuliformis*** | Singh | 4227 | NY | India: Ladakh | 1972 | a9ed6de2-d2e9-4a78-802a-a6588e664937 |
| ***Astragalus subumbellatus*** | Ledingham | 4048 | NY | Canada: Saskatchewan | 1965 | ac65f99d-505a-4a9e-a1b2-15a0d8e2d40e |
| ***Astragalus subvestitus*** | Twisselmann | 17286 | NY | United States of America: California: Tulare | 1970 | d2ce2520-9239-4192-8527-2632fb1f172d |
| ***Astragalus suffalcatus*** | USDA-ARS | 380723 | NY | Islamic Republic of Iran | 1977 | a9ea32c5-8a91-470c-869c-20a83e14efcc |
| ***Astragalus sulcatus*** | [Cyrillic] | | HUH | Russian Federation | 1967 | fead2a20-af8e-4f68-b381-32770fc4b9f1 |
| ***Astragalus sumbari*** | [Cyrillic] | | NY | Turkmenistan | 1975 | ad76ac6a-4d7f-47f3-bc82-20ab19cd0505 |
| ***Astragalus sungpanensis*** | Podlech | 34562 | HUH | People’s Republic of China | 2005 | fe52f1db-4281-4268-9305-a0fd79e0edfd |
| ***Astragalus susianus*** | Ledingham | 4081 | NY | Islamic Republic of Iran | 1965 | a9e7f920-2a31-4fe0-a809-2eb96ac09ff1 |
| ***Astragalus szovitsii*** | Barkworth | s.n. | NY | Armenia: Ararat: Ararat | 2003 | d06769f4-f6dd-4803-97c3-a01f4e1369aa |
| ***Astragalus tabrizianus*** | Grant | 16283 | MO | Islamic Republic of Iran: East Azerbaijan | 1964 | 0c657308-17d2-46a5-8a1e-f407795a455d |
| ***Astragalus takharensis*** | Anders | 6770 | NY | Afghanistan: Takhar | 1971 | a95c346b-20b2-43ae-a9d8-553ef24bd910 |
| ***Astragalus tarijensis*** | Ledingham | 4489 | NY | Argentina | 1966 | d1d6c4b8-f3d1-4123-a6d0-25d23baef610 |
| ***Astragalus taschkendicus*** | Podlech | 30156 | NY | Afghanistan: Takhar: Farkhar | 1977 | a9576c0f-eb44-4b53-96c7-0831cf466ee4 |
| ***Astragalus tatjanae*** | Anders | 5650 | HUH | Afghanistan | 1971 | fc698b5a-5862-424f-ae88-cc28201913b0 |
| ***Astragalus tegetarioides*** | Meinke | 6086 | NY | United States of America: Oregon: Harney | 1991 | d33b4fcb-456f-4c68-a267-189f7b99fcbc |
| ***Astragalus tegetarioides (Astragalus anxius)*** | Nelson | 5988 | NY | United States of America: California: Lassen | 1980 | d5fbce34-5c78-436a-a250-cdaed1c2fc51 |
| ***Astragalus teheranicus*** | Ledingham | 4200 | NY | Islamic Republic of Iran: Kermanshah: Harsin | 1965 | a9549361-0e11-4146-9f63-083f8d7c6756 |
| ***Astragalus tenellus*** | Kirkpatrick | 1610 | NY | United States of America: Colorado: Pitkin | 2010 | cc177f99-9208-41ef-9246-8fceb8a99ae9 |
| ***Astragalus tener*** | Crampton | 3315 | NY | United States of America: California: Yolo | 1956 | d3204468-c2b6-4ab7-b21b-9b7693dee684 |

| | | | | | | |
|---|---|---|---|---|---|---|
| ***Astragalus tephrodes*** | Crosswhite | 700 | FLAS | United States of America: Arizona: Yavapai | 1960 | ea7e801a-e927-492c-bcfe-321f6b7f8b74 |
| ***Astragalus tephrolobus*** | Sumnevicz | s.n. | NY | Russian Federation: Altai | 1931 | acd70b61-1b18-4485-a6dd-1fdc76f6d262 |
| ***Astragalus terminalis*** | Lesica | 2730 | NY | United States of America: Montana: Beaverhead | 1983 | d408142f-d0a9-432c-9e1b-5ef18ecdcb47 |
| ***Astragalus terrae-rubrae*** | Butkov | 6270 | NY | Uzbekistan: Qashqadaryo | 1936 | acd0dd0f-01bc-4756-aeda-13a91ca5d61a |
| ***Astragalus testiculatus*** | Dubovyk | s.n. | NY | Ukraine: Luhansk: Milove | 1958 | cf9b710f-4a62-457f-ae6b-f57e67e97416 |
| ***Astragalus tetrapterus*** | Packard | 78-64 | NY | United States of America: Oregon: Malheur | 1978 | d3b77726-edb8-4a29-96c8-0acb8a079cca |
| ***Astragalus thurberi*** | Rena | 2003-406 | CAS | Mexico: Sonora: Fronteras | 2003 | 169bd0f5-7c34-4251-a8fc-602ba8a31a97 |
| ***Astragalus tibetanus*** | Dorn | 5274 | NY | United States of America: Wyoming: Fremont | 1991 | d3b60cda-6624-46b0-9767-df913290c92c |
| ***Astragalus tidestromii*** | Sanders | 39143 | NY | United States of America: California: Inyo | 2011 | d3a757d3-67c5-4f5b-bd02-9305400f7f2f |
| ***Astragalus tiehmii*** | Morefield | 5483 (dupl. c) | NY | United States of America: Nevada: Washoe | 1991 | d39f27af-1fe3-44c1-ab76-0cff37a37e18 |
| ***Astragalus titanophilus*** | Reichenbacher | 1687 | NY | United States of America: Arizona: Coconino | 1985 | d399154c-db6c-4208-aa38-e05cb3c22225 |
| ***Astragalus toanus*** | Holmgren | 4949 | NY | United States of America: Idaho: Owyhee | 1971 | d407d980-97dd-4cad-b970-4d1fdffd5a74 |
| ***Astragalus tolucanus*** | Rzedowski | 25902 | CAS | Mexico: M√©xico | 1968 | 1643370d-44d1-404a-ab06-24188c1cbc18 |
| ***Astragalus tongolensis*** | Boufford | 37356 | NY | People’s Republic of China: Sichuan | 2006 | aca8ab5a-1d46-4782-8800-2253144fb48f |
| ***Astragalus toquimanus*** | Tiehm | 14506 | NY | United States of America: Nevada: Nye | 2004 | d3faffe8-0d4e-469f-b694-2fddd692b359 |
| ***Astragalus tortuosus*** | Lamond | 4474 | NY | Islamic Republic of Iran: Kordestan | 1971 | aae5e30f-e13f-4aed-94aa-aefe8989179d |
| ***Astragalus trachycarpus*** | Bochantsev | 1330 | NY | Tajikistan | 1960 | acba38d2-fc85-44db-a908-3fdb3a0f7910 |
| ***Astragalus transoxanus*** | Kurdrjaschev | 6271 | NY | Uzbekistan | 1929 | acbcd986-6441-43de-9975-51e548438475 |
| ***Astragalus traskiae*** | Junak | SN-005 | NY | United States of America: California: Ventura | 1983 | d3964ecf-d692-4cb1-af40-7d0dc3f89bda |
| ***Astragalus tribuloides*** | O.H Volk | 1898 | NY | Afghanistan: Kabul | 1951 | ce4ddc73-22ba-46fe-b4cd-4610879e0c53 |
| ***Astragalus tricarinatus*** | Liston | 852 | NY | United States of America: California: Riverside | 1992 | d393ca5d-c3a6-4d54-b34d-89dba26eb207 |
| ***Astragalus trichanthus*** | Goloskokov | 4821 | NY | Kazakhstan | 1966 | aca3b531-ae6b-4c80-89a3-18d5acc7f617 |

| | | | | | | |
|---|---|---|---|---|---|---|
| ***Astragalus tricholobus*** | Ledingham | 4182 | NY | Islamic Republic of Iran: Khorasan-e Jonubi | 1965 | ce3d0636-9a3c-497b-9f9b-9368f852985e |
| ***Astragalus trichopodus*** | Reveal | 6902 | CAS | Mexico: Baja California: Ensenada | 1988 | 169a5b28-8233-4c4c-9412-887a1a447f41 |
| ***Astragalus triflorus*** | Weigend | 97/741 | NY | Peru: Arequipa: Caraveli | 1997 | ce38dd57-3171-49d8-930a-933387556260 |
| ***Astragalus trifoliolatus*** | Ledingham | 3471 | NY | Canada: Saskatchewan | 1962 | ab192507-4fd7-4155-9cff-b62d41f64408 |
| ***Astragalus trigonus*** | Amin | s.n. | MO | Egypt | 1976 | 163cda8d-cefe-486c-b39d-9752b1cbf8b4 |
| ***Astragalus trimestris*** | Davis | D. 48556 | NY | Morocco | 1969 | d02b3a47-782b-4cfc-8fc5-110eb0c0da37 |
| ***Astragalus triqueter*** | G.F. Ledingham | 4063 | NY | Islamic Republic of Iran | 1965 | ce368241-66cf-4125-b226-6814fb1b6413 |
| ***Astragalus troglodytus*** | Mielke | H2173 | NY | United States of America: Arizona: Coconino | 1983 | d387d4c3-c17a-4ec7-b524-d26d5877296d |
| ***Astragalus tugarinovii*** | Kharkevich | 537 | NY | Russian Federation: Kamchatskiy | 1975 | ac9ef16a-c8e3-4d8a-be14-f172c59f5164 |
| ***Astragalus turbinatus*** | Va[?][?]sosna | 400 | NY | Uzbekistan | 1968 | ac9bd420-52d8-4b9e-9a1d-0a9dccb7357e |
| ***Astragalus turcomanicus*** | Vasak | s.n. | NY | Turkmenistan: Ashabad | 1986 | ac8d6224-05a0-4ea2-842c-fa322fd07170 |
| ***Astragalus turczaninovii*** | Putschkov | 6272 | NY | Kazakhstan | 1866 | ac8d3fbc-1123-4600-a80c-a210fecf6049 |
| ***Astragalus turkmenorum*** | Borissova | 3350 | NY | Turkmenistan | 1934 | ac850f07-aa13-4e5d-87dc-1b1c45aa0da7 |
| ***Astragalus turolensis*** | Reverchon | 6. 92. | NY | Spain: Teruel | 1892 | cf9e35b7-ba2a-45b3-a03d-f786e5ae33fe |
| ***Astragalus tweedyi*** | Hufford | 3270 | NY | United States of America: Washington: Benton | 1999 | d31b4b94-4eb5-4c6a-b9cb-d8d34d05f5e5 |
| ***Astragalus tyghensis*** | Hitchcock | 20736 | NY | United States of America: Oregon: Wasco | 1955 | d40a6611-91dc-4adb-a626-8f00fe1a8e1f |
| ***Astragalus uliginosus*** | USDA-ARS | 420694 | NY | Russian Federation: Irkutsk | 1981 | ad6b48cc-0b0c-4542-ad3e-a27c8772749d |
| ***Astragalus umbellatus*** | Welsh | 10057 | NY | Canada: Yukon Territory | 1970 | d288a3de-9c31-4817-9510-db9793afe879 |
| ***Astragalus umbraticus*** | Liston | 872 | NY | United States of America: Oregon | 1992 | d281d11b-2627-480c-a0f8-16a05d45120c |
| ***Astragalus uncialis*** | Welsh | 20619 | NY | United States of America: Utah: Millard | 1981 | d27ea0bd-c1d2-4401-bf57-0e46dc3ca0b8 |
| ***Astragalus uniflorus*** | Feuerer | 11040b | NY | Bolivia (Plurinational State of): Bautista Saavedra: La Paz | 1982 | d1ca603c-53d3-4bb1-9864-aa8bce5b4984 |
| ***Astragalus unifoliolatus*** | Belianina | 161 | HUH | Turkmenistan | 1975 | fe6b0c77-e847-42e9-a799-8ebf0cd92c61 |
| ***Astragalus uraniolimneus*** | v.l komarovi | 661 | NY | United States of America | 1956 | ad6a7912-a5f9-4b15-b71a-3c229cba9183 |
| ***Astragalus utahensis*** | Foster | 10410 | FLAS | United States of America: Wyoming: Uinta | 1991 | ea6c3742-0857-4fcc-bc46-fbe481766062 |
| ***Astragalus vaccarum*** | Yen | 2669 | CAS | Mexico: Chihuahua | 1994 | 16867fcf-9cfe-426d-81d1-4f11f7250653 |

| | | | | | | |
|---|---|---|---|---|---|---|
| ***Astragalus valerianensis*** | Zollner | 5701 | NY | Chile: Curico | 1971 | d23b0b99-3074-41fe-98bc-2afc2d195bf3 |
| ***Astragalus vallaris*** | Atwood | 12067 | NY | United States of America: Idaho: Washington | 1986 | d2c67619-16aa-4e4f-b3ea-f23f7f9e94b4 |
| ***Astragalus varius*** | Dubovyk | s.n. | NY | Ukraine: Luhansk: Milove | 1958 | cfbe44d2-c49e-4447-877f-575e15afc80e |
| ***Astragalus varzobicus*** | Vvedensky | 6242 | HUH | Uzbekistan | 1929 | fe2b9650-a9e3-4eef-b06f-ed2e8a80ae2b |
| ***Astragalus vegetus*** | Barneby | 384777 | NY | Islamic Republic of Iran | 1979 | ce34f589-e8f1-4a9a-89a8-0192e310a438 |
| ***Astragalus versicolor*** | Popov | 3827 | HUH | Russian Federation: Irkutsk | 1952 | fe24af05-3573-4e50-9c50-fa861709abe3 |
| ***Astragalus verticillatus*** | Ricardi | 3182 | NA | Chile: Colchagua: San Fernando | 1955 | e741730b-806a-48cc-b05e-cc5b5691a10a |
| ***Astragalus verus (Astracantha meana)*** | [Cyrillic] | s.n. | MO | [Cyrillic] | 1957 | 0c287c56-356e-4028-9822-0ae0c74b571a |
| ***Astragalus verus (Astracantha multifoliolata)*** | [Cyrillic] | s.n. | MO | Turkmenistan | 1954 | 0c315497-bddc-4caf-b655-cbabcb5301d9 |
| ***Astragalus verus (Astracantha piletoclada)*** | [Cyrillic] | s.n. | MO | Turkmenistan | 1953 | 0c54bbac-3871-4eb9-bd54-72955cab949f |
| ***Astragalus verus (Astracantha meschhedensis)*** | Kurbanov | 1474 | MO | Turkmenistan | 2001 | 0c2c24f9-1785-4619-a8c7-70c8db77f153 |
| ***Astragalus verus (Astracantha strobilifera)*** | Stewart | 22962a | NY | Pakistan: Kashmir | 1946 | aa79ccfe-496a-46ae-bfd8-f9da3935d730 |
| ***Astragalus verus (Astracantha pulvinata)*** | Vvedensky | 6267 | HUH | Turkmenistan | 1927 | 009c6e6a-ac26-4552-bda5-298d0ac7b463 |
| ***Astragalus vesicarius*** | Charpin | 8325 | NY | France: Hautes-Alpes: Laragne | 1969 | cfaf483a-69fb-4d67-a2c4-d8281e96d060 |
| ***Astragalus vesiculosus*** | Ellenberg | 4625 | NY | Chile | 1971 | d23aea77-7b17-43f9-881a-b4a70922bd83 |
| ***Astragalus vexans*** | Podlech | 29748 | NY | Afghanistan: Herat: Sudhang des Sabzak-Passes | 1977 | ce298ce4-eb32-4022-b0bd-a32b5ab217b5 |
| ***Astragalus vicarius*** | Goloskokov | 4469 | NY | Kazakhstan: Zhambyl | 1963 | ad6641da-3f4a-4929-af78-881bcf8665ac |
| ***Astragalus villosus*** | Herring | 1108 | FLAS | United States of America: Florida: Columbia | 1993 | e9daa50b-2770-467b-b971-075413e87fc1 |
| ***Astragalus viridiflorus*** | Popov | 6244 | HUH | Kazakhstan | 1930 | fe130d4e-709d-429d-91bc-cd8f6aca1851 |
| ***Astragalus vulpinus*** | Androssov | 7424 | NY | Kazakhstan: Aktjubinsk | 1908 | cfbe144e-0be4-4b18-8f68-6706b4f530fb |
| ***Astragalus wardii*** | M. Ben Franklin | 7191 | FLAS | United States of America: Utah: Garfield | 1990 | ea8cf49b-3491-4f91-888f-dad412a383f4 |
| ***Astragalus waterfallii*** | Spellenberg | 4638 | NY | United States of America: New Mexico: Otero | 1977 | d27a648d-9876-4e78-b94c-ab141299a452 |
| ***Astragalus webberi*** | Hanson | s.n. | NY | United States of America: California: Plumas | 1987 | d2bdf26b-9aed-49ed-9539-e4cb5e4cc26e |

| | | | | | | |
|---|---|---|---|---|---|---|
| ***Astragalus webbianus*** | Singh | 4232 | NY | India: Kashmir | 1972 | ce1cc552-74c5-4757-b33a-7d6b814d11a7 |
| ***Astragalus weberbaueri*** | Sagdstegui | 16084 | NY | Peru: Santiago de Chuco | 1997 | d22a7d82-0ed0-4ada-838b-f142fbf80e89 |
| ***Astragalus weddellianus*** | Weigend. M. | 2000/478 | NY | Peru: Puno: Juli | 2000 | d226ddb7-28fa-4125-9ab1-0269199d9d9c |
| ***Astragalus welshii*** | Welsh | 6477 | NY | United States of America: Utah: Wayne | 1967 | d272c4d9-ce87-4fc8-8baa-3b583017f7a8 |
| ***Astragalus wetherillii*** | Welsh | 6235 | NY | United States of America: Colorado: Garfield | 1967 | d2bd4007-0215-4885-ba4e-655d99ff86f3 |
| ***Astragalus williamsii*** | Kowalczyk | 9 | NY | Canada: Yukon Territory | 1971 | d2bc095e-13c4-4683-a6a4-6582c26d385a |
| ***Astragalus wingatanus*** | d Atwood | 22045 | NY | United States of America: Utah | 1997 | d26cc0a0-b87d-4ec6-a58e-70b3da543d70 |
| ***Astragalus wittmannii*** | Spellenberg | 5991 | NY | United States of America: New Mexico: Harding | 1981 | d2b31d7f-4315-4afe-a50e-7006fda3ed9f |
| ***Astragalus woodruffii*** | Atwood | 8681 | NY | United States of America: Utah: Wayne | 1982 | d2692522-babc-4815-bd08-cf663ce8028c |
| ***Astragalus wootonii*** | Ventura | 632 | CAS | Mexico: Hidalgo: Tepeapulco | 1975 | 167fb7e0-4791-40b3-b1cf-fc749e1700e3 |
| ***Astragalus wrightii*** | Tharp | 44041 | NY | United States of America: Texas: Travis | 1944 | d2b0a388-ad58-458f-b573-2517d27d6d6e |
| ***Astragalus xanthomeloides*** | Vvedensky | 6245 | HUH | Unknown Country | 1930 | fde0bf9e-b017-4f03-8a0f-b4203cd52fad |
| ***Astragalus xiphidioides*** | Hevron | 698b | NY | United States of America: Arizona: Navajo | 1990 | d2a90952-64ca-4d81-a826-2515bdc43ebc |
| ***Astragalus xiphidium*** | Lachashvili | 499 | NY | Georgia: East Georgia: Dedoplistkali District | 2006 | cf208803-d32d-4e55-b8e6-2977672e074f |
| ***Astragalus yoder-williamsii*** | Rosentreter | 3489 | NY | United States of America: Nevada: Humboldt | 1984 | d2a06b8f-5fa0-4796-9220-8aa2104cb8f7 |
| ***Astragalus yunnanensis (Astragalus fenzelianus)*** | Boufford | 31818 | HUH | People's Republic of China: Xizang (Tibet) | 2004 | fa38fcff-372c-448e-9f76-4b79ecd2e65c |
| ***Astragalus zacharensis*** | Cao | 53 | FLAS | People's Republic of China: Heilongjiang: Daqingshan | 1991 | ea772c5c-0aef-499b-bb23-1e45b02d0957 |
| ***Astragalus zagrosicus*** | Ledingham | 4103 | NY | Islamic Republic of Iran: Fars | 1965 | ce0e36f5-c79c-414f-be06-5bf20e2fb993 |
| ***Astragalus zanskarensis*** | Webster | 5784 | HUH | Pakistan: Baltistan: Skardu | 1955 | fd62c73b-aa74-438e-83bc-e3eb0b5371b5 |
| ***Astragalus zingeri*** | Skvortsov | s.n. | NY | Russian Federation: Saratov: Hvalynsk | 1993 | cf1c4ddb-9cc5-459a-8336-5dee469e1b1a |
| ***Astragalus zionis*** | Neese | 16890 | NY | United States of America: Utah: Washington | 1985 | d25cab54-46aa-432e-89ea-31af68d6cd92 |
| ***Biserrula epiglottis*** | Wilczek | 178 | MO | Morocco | 1928 | 1579f02c-9844-41d7-abc9-307a179d6358 |

| | | | | | | |
|---|---|---|---|---|---|---|
| ***Carmichaelia violacea*** | Chapman | 258520 | MO | New Zealand | 1970 | 17abaf6f-55c2-4cb5-a7aa-28ea2ed1f278 |
| ***Clianthus puniceus*** | Breteler | 13684 | MO | New Zealand: North Island | 1996 | 172f84d3-6023-48a6-8fbd-1426cbe05a34 |
| ***Colutea abyssinica*** | Simon | 857 | MO | Tanzania: Arusha: Monduli | 2001 | 16e6a46a-09bc-4471-8a31-d8ac5a80dd74 |
| ***Eremosparton flaccidum*** | Belyanina | 26 | MO | Turkmenistan: Kara-Kumy: Repetek | 1979 | 6ce6a816-4321-4e3f-8c67-e64506df10ac |
| ***Erophaca baetica*** | Lewalle | 12219 | MO | Morocco | 1989 | 1511ff5a-b9e2-4899-aa11-64ef7fec40c4 |
| ***Hedysarum boreale*** | Kass | 3262 | FLAS | United States of America: Wyoming: Sweetwater | 1991 | f0b6e23e-7346-42df-a499-aa05d5d52a6b |
| ***Lessertia capitata*** | Goldblatt | 3673 | MO | South Africa: Cape | 1976 | 5048e728-0a1c-48b2-9b7a-9e872442c8b6 |
| ***Onobrychis conferta*** | Dubuis | 13246 | MO | Algeria: M’Sila: Bou-Saada | 1986 | 6b5615a8-6aa7-42b6-8be0-225c14135843 |
| ***Oxytropis aciphylla*** | Shiquan | 20[5?]4 | MO | [Chinese] | 1980 | 04e037e9-79f2-4c3b-aa3d-a66bad0da0a2 |
| ***Oxytropis adamsiana*** | Petrovsky | 6576 | MO | Russian Federation: Sakha: Bulunsky | 1984 | 04f9c33c-cfb6-4535-bd55-711579fe7562 |
| ***Oxytropis albana*** | Atha | 2698 | MO | Georgia: Mtskheta-Mtianeti: Kazbegi | 2002 | 05223403-4639-4282-b021-9102a14d4568 |
| ***Oxytropis arctica*** | Gillespie | 8954-2 | MO | Canada: Northwest Territories | 2009 | 0cfa9a7b-6f17-469e-a789-65d95b6312ec |
| ***Oxytropis aspera*** | Vvedensky | 6274 | MO | Uzbekistan | 1929 | 0539eb06-7bcf-4d0b-8aad-f7c63af4882b |
| ***Oxytropis aucheri*** | Rechinger | 35436 | MO | Afghanistan: Paktia | 1967 | 05166e43-8196-4b6e-b63a-9b947ad18a59 |
| ***Oxytropis besseyi*** | Foster | 10424 | FLAS | United States of America: Wyoming: Sweetwater | 1991 | ec7e5497-5fd0-474b-907c-e0910a64d888 |
| ***Oxytropis bicolor*** | Wang | 91244 | MO | People’s Republic of China: Gansu | 1991 | 04f08eb9-86a4-43f0-9e41-acb43d8802c9 |
| ***Oxytropis campanulata*** | Elias | 7094 | MO | Russian Federation: Altay: Ust-Kansky | 1983 | 04f16b2c-8c6d-4aa3-9a25-ffab9d5bbab2 |
| ***Oxytropis campestris*** | Welsh | 24742 | FLAS | United States of America: South Dakota: Custer | 1991 | ec7bf570-b8a8-49c7-a6b8-221923f9dccd |
| ***Oxytropis chionobia*** | Sodombekov | KPL_00628 | MO | Kyrgyzstan: Naryn: Kochkor | 2006 | 05085790-3aed-4d74-a6c6-dabee0d99fe3 |
| ***Oxytropis ciliata*** | Cao | 130 | FLAS | People’s Republic of China: Hohhot: Daqingshan | 1991 | ec623678-7df4-4f99-bbd8-3485e280989a |
| ***Oxytropis coerulea*** | Wang | 368 | MO | People’s Republic of China | 1998 | 051c37de-48c5-4142-a307-0ba2b0c42b29 |
| ***Oxytropis czukotica*** | Kharkevich | 550a | MO | Russian Federation: Kamchatka: Olyutorsky | 1974 | 0536240f-cae5-4dac-b8e1-b3479df1b86e |
| ***Oxytropis deflexa*** | Harriman | 18215 | FLAS | United States of America: Colorado: Park | 1984 | ec72c93d-6656-4081-9ad2-9aa7b6eed9f5 |

| | | | | | | |
|---|---|---|---|---|---|---|
| ***Oxytropis eriocarpa*** | Khanmichun | 19 | MO | Russian Federation: Siberia: Tuva | 1973 | 054c6d23-aae8-412f-a80b-453df977ed4f |
| ***Oxytropis exserta*** | Kharkevich | 552c | MO | Russian Federation: Kamchatka: Olyutorsky | 1975 | 054fbe06-1ce0-4843-b0c9-cbc45e45ed14 |
| ***Oxytropis falcata*** | Ho | 1534 | MO | People's Republic of China: Qinghai: Hainan Zangzu | 1993 | 05200ba5-732a-4c5a-a1e6-5087e6ac107a |
| ***Oxytropis grandiflora*** | Budantzev | 759 | MO | Mongolia: Dornod: Khalkhgol | 1987 | 052e9530-04f2-43dd-9584-bf4f82d60f89 |
| ***Oxytropis hirta*** | Li | 2548 | MO | [Chinese] | Not Shown | 054b72af-3449-4b19-bd0a-852bd68ef3e1 |
| ***Oxytropis humifusa*** | Chukarina | 4592 | MO | Tajikistan: Pamir | 1971 | 0c029dfb-5c36-4a15-88a7-db95d004bfde |
| ***Oxytropis immersa*** | AchmetsPeople's Republic of China | s.n. | MO | Tajikistan | 1968 | 054b6cd4-6dd5-45a8-bb86-374946740672 |
| ***Oxytropis jacquinii*** | Bussman | 43239 | MO | Slovenia: Gori≈°ka | 1988 | 04dc0e5f-a417-485c-a2e6-39e798ce42c2 |
| ***Oxytropis jonesii*** | Franklin | 7198 | BRIT | United States of America: Utah: Garfield | 1990 | 6c5bac9c-b0a6-4372-8eae-9174920812c5 |
| ***Oxytropis jordalii*** | Barker | BG07-058B | MO | United States of America: Alaska: Valdez Cordova | 2007 | 0cfa8401-a53e-4807-b100-c2d328f2d260 |
| ***Oxytropis kansuensis*** | Ho | 165 | MO | People's Republic of China: Qinghai: Hainan Zangzu | 1993 | 0558ad39-fa5e-471a-9fd4-34eda1cad5c4 |
| ***Oxytropis kobukensis*** | Lipkin | 84-68 | MO | United States of America: Alaska | 1984 | 0ce21f3d-dba9-4a54-943c-d7bd6969b4dd |
| ***Oxytropis koyukukensis*** | Murray | 3692 | MO | United States of America: Alaska | 1973 | 0ce1a27f-f8ed-4e21-bfad-9057629c43fa |
| ***Oxytropis lagopus*** | Henderson | 90-65 | MO | United States of America: Texas: Donley | 1990 | 0cdeadb3-df15-4abf-a135-980ee2e7ae28 |
| ***Oxytropis lambertii*** | Welsh | 24764 | FLAS | United States of America: Wyoming: Weston | 1991 | ec6b7cfa-5fc7-4556-986a-beb4c68e073e |
| ***Oxytropis lapponica*** | Oxborne | 516 | MO | Kyrgyzstan: Talas: Talas | 2008 | 0558d0b6-43c4-4b8e-aaf5-b0dda32da2fe |
| ***Oxytropis macrocarpa*** | Sodombekov | KPL_00379 | MO | Kyrgyzstan: Chui: Ysyk-Ata | 2005 | 055d0761-1305-4f61-abe5-4a771e96b6bc |
| ***Oxytropis maydelliana*** | Parker | 8885 | MO | United States of America: Alaska: Bethel | 1999 | 0cdf5509-b999-451d-9837-2a18fe044ce4 |
| ***Oxytropis melanocalyx*** | Ho | 421 | MO | People's Republic of China: Qinghai: Golog Autonomous Tibetan Prefecture | 1993 | 056166d5-71c1-4d52-97d3-98c33a20bf4e |
| ***Oxytropis mertensiana*** | Murray | 3172 | MO | United States of America: Alaska | 1970 | 0cdb0bb8-da1f-4b2f-b43c-a558ce6a920b |
| ***Oxytropis michelsonii*** | Abdusaljamora | 5212 | MO | Tajikistan | 1967 | 056b7240-b44b-4ba9-9ad1-77d036bceee6 |
| ***Oxytropis microphylla*** | Iokawa | 40011 | MO | Nepal: Gandaki Pradesh: Mustang | 2003 | 056c0db5-9f71-4374-86b8-f3286463b4f0 |

| | | | | | | |
|---|---|---|---|---|---|---|
| ***Oxytropis mollis*** | Iokawa | 40004 | MO | Nepal: Gandaki Pradesh: Mustang | 2003 | 05829b3a-6084-4d45-b522-c63677f78684 |
| ***Oxytropis multiceps*** | King | 11696 | MO | United States of America: Wyoming: Albany | 2001 | 0cd03c8c-483e-4d49-86b4-e18f057447c6 |
| ***Oxytropis nana*** | Meyer | 1172 | MO | United States of America: Wyoming: Albany | 2011 | 0cd298a0-65fe-46f5-85e5-d84698c32886 |
| ***Oxytropis neglecta*** | Charpin | 19752 | MO | Italy: Cureo | 1985 | 04e46b41-cd0e-4760-b088-3bb9ec799dcb |
| ***Oxytropis nigrescens*** | Lipkin | 82 | MO | United States of America: Alaska: Bethel | 1993 | 0cf0184b-7b2c-4e80-9820-1ccba3ffb174 |
| ***Oxytropis nutans*** | Sodombekov | KPL_00412 | MO | Kyrgyzstan: Chui | 2005 | 05610514-f170-43a4-af0e-26bf48d4cd83 |
| ***Oxytropis obnapiformis*** | Weber | 12656 | TEX | United States of America: Colorado: Moffat | 1965 | 10da7bb1-890c-41ee-b9f8-22cce4331915 |
| ***Oxytropis ochotensis*** | [Cyrillic] | s.n. | MO | Russian Federation: Magadan: Khasynskiy | 1969 | 0569f460-f5b2-4912-8996-f9cece956a9e |
| ***Oxytropis ochrantha*** | Cao | 66 | FLAS | People's Republic of China: Inner Mongolia: Hohhot | 1991 | ec4cf8bb-e12c-4096-bc4c-d20095534237 |
| ***Oxytropis ochrocephala*** | Boufford | 40287 | MO | People's Republic of China: Sichuan: Aba Zangzu Qiangzu | 2007 | 056db88c-af48-4a6f-9a20-1e4407dfdeac |
| ***Oxytropis ochroleuca*** | Smith | 11377 | MO | People's Republic of China: Sikang: Kangting | 1934 | 05896fe4-4142-440d-b620-66cdb6f0fa24 |
| ***Oxytropis oreophila*** | Welsh | 26992 | MO | United States of America: Utah: Garfield | 1998 | 0ccb45a1-2f4c-414b-aa48-ff6e351fb8d6 |
| ***Oxytropis oxyphylloides*** | Malyschev | 4031 | MO | Russian Federation: Irkutsk | 1955 | 058b0554-b121-4870-a5b6-9ef5cb4943bc |
| ***Oxytropis parryi*** | Huber | 3813 | MO | United States of America: Utah: Wasatch | 1998 | 0cc63caa-d350-43f7-9546-427cc835b4b1 |
| ***Oxytropis pauciflora*** | Ho | 768 | MO | People's Republic of China: Qinghai: Golog Tibetan | 1993 | 05943296-8355-4be9-a80e-bb0bdd489a5e |
| ***Oxytropis pilosa*** | Kleineberg | 6112 | MO | Germany | 1973 | 04c7443c-a46d-4dc5-ac86-93f4b134976a |
| ***Oxytropis podoloba*** | Goloskokov | 4421 | MO | Kazakhstan | 1959 | 059f86d6-c3fb-4833-ae53-02e23d250220 |
| ***Oxytropis poncinsii*** | Medvedev | 49 | MO | Tajikistan: Pamir | 1985 | 05a977b1-100b-4503-898a-89fc411a2a34 |
| ***Oxytropis qinghaiensis*** | Ho | 589 | MO | People's Republic of China: Qinghai: Golog Tibetan | 1993 | 052ae417-7fdd-431c-a0af-9ea618573822 |
| ***Oxytropis racemosa*** | [Chinese] | 1139 | MO | [Chinese] | 1983 | 05bbb0e5-a1e9-4974-ba4e-52fb59f1d49c |
| ***Oxytropis recognita*** | Zomonosova | s.n. | MO | Russian Federation: Altay: Kosh-Agachskiy | 1982 | 05b29410-45d2-480d-9727-6fe332657c58 |
| ***Oxytropis riparia*** | Rechinger | 18122 | MO | Afghanistan: Bamian | 1962 | 05ca7139-12a7-40fc-a2fe-7605a4e6ee26 |

| | | | | | | |
|---|---|---|---|---|---|---|
| ***Oxytropis roseiformis*** | Botschantzev | s.n. | MO | Tajikistan | 1960 | 05cc36db-c66c-4b46-b7a1-b661a402f629 |
| ***Oxytropis savellanica*** | Akhmetshina | s.n. | MO | Tajikistan | 1968 | 058a88f8-b8e2-4674-b9ef-cb8687849342 |
| ***Oxytropis scammaniana*** | Parker | 8768 | MO | United States of America: Alaska: Bethel | 1999 | 0cbe9132-c327-4533-b681-902d6295a7cf |
| ***Oxytropis sericea*** | Foster | 10416 | FLAS | United States of America: Wyoming: Uinta | 1991 | ec6918a2-3596-4ad3-933b-7529e4921f43 |
| ***Oxytropis splendens*** | van der Werff | 26017 | MO | United States of America: New Mexico: Taos | 2015 | 0cba2d72-282e-420d-a1ca-1a6bbbb1a2cd |
| ***Oxytropis squammulosa*** | Kamelin | 155 | MO | Mongolia: Dornod: Matad | 1987 | 056a94fc-ce24-41c0-aa30-f8e379a919f9 |
| ***Oxytropis strobilacea*** | [Cyrillic] | s.n. | MO | Russian Federation: Tyva Republic: Mongun-Tayginsky | 1980 | 0597ccfa-f23a-40a3-a8b0-959ec354319a |
| ***Oxytropis talassica*** | Sodombekov | KPL_00603 | MO | Kyrgyzstan: Naryn: Kochkor | 2006 | 05a47f68-ed30-47df-9d58-8a9eb015c16d |
| ***Oxytropis taochensis*** | Wang | 901101 | MO | People’s Republic of China: Gansu: Linxia Hui | 1991 | 05ab4d36-3482-4596-9788-8767772d8f7a |
| ***Oxytropis tianschanica*** | Stanjukovicz | 7844 | MO | Tajikistan: Gorno-Badakhshan | 1952 | 05b54e51-e2c6-4493-b485-3c57de93e589 |
| ***Oxytropis vassilievii*** | Kaharkevich | 1118 | MO | Russian Federation: Kamchatka: Ayano-Maysky | 1977 | 05c8944c-17e5-4f8e-be6b-642dc840faea |
| ***Oxytropis viscida*** | Scott | 8130 SAL | FLAS | United States of America: Wyoming: Fremont | 1991 | ec660bed-8cc7-405e-9b96-5aa4aa326060 |
| ***Oxytropis williamsii*** | Ross | TUCH-MO 66 | MO | Nepal: Gandaki Pradesh: Manang | 2009 | 05cb8d42-5817-4b80-bdf1-4bd173c735b4 |
| ***Oxytropis yunnanensis*** | Ho | 1640 | MO | People’s Republic of China: Qinghai: Yushu | 1996 | 05db621b-0925-4efb-ad22-dd71787d9206 |
| ***Phyllolobium balfourianum (Astragalus prattii)*** | Boufford | 28339 | HUH | People’s Republic of China: Sichuan Province: Xiangcheng Xian | 1998 | 0102421d-c383-40d2-8f49-26961abc066a |
| ***Phyllolobium balfourianum*** | Rock | 5327 | NY | People’s Republic of China: Yunnan | 1922 | ced84ef6-9072-4d71-855c-906d9da645f3 |
| ***Phyllolobium donianum*** | Ludlow | 19492 | HUH | Bhutan: Bumthang | 1949 | fc851e2f-7835-430e-8afb-782f9d879a1b |
| ***Phyllolobium tribulifolium*** | Boufford | 26850 | NY | People’s Republic of China | 1995 | ad73aef4-06b6-4df1-a240-3dd854d0dd8e |
| ***Phyllolobium tribulifolium*** | Stewart | 22,839 | NY | India: Punjab | 1946 | ce1449be-a037-4155-981d-563220506d48 |
| ***Podlechiella vogelii*** | Mokhtar | 5 | MO | Egypt | 1980 | 163a83ab-3015-4aa5-af58-ce9d55b56959 |
| ***Podolotus hosackioides*** | Rashid | 25853 | NY | Pakistan: Azad Kashmir | 1954 | a9f94d10-8570-4ee9-abb0-99d8a86ca999 |
| ***Smirnowia turkestana*** | Belianina | 160 | MO | Turkmenistan | 1975 | e143d7fc-df8a-44d7-ac16-8c642460dafa |
| ***Sphaerophysa salsula*** | Hatle | 167 | FLAS | United States of America: Wyoming: Fremont | 1990 | 1967e7c1-a057-45ef-903c-f9a3f7541399 |

| | | | | | | |
|---|---|---|---|---|---|---|
| ***Sutherlandia microphylla*** | van Wyk | 2660 | TEX | South Africa: Cape | 1986 | 0a73748a-3142-4baf-8877-b624a1f03324 |
| ***Swainsona formosa*** | Herman | 658 | MO | South Africa: Transvaal | 1982 | 175c43b9-bf41-4a2f-89a2-6943e974414f |
| ***Wisteria sinensis*** | Degener | 35249 | | United States of America: Hawaii | 1981 | 2f27b596-4028-4b40-b9f3-d222890dd14e |

**Table S1.** Accession table.

**Table S2.** Results for phylogenetic signal tests.

| Variable | λ | *p*-value | *p*-value (Hochberg-corrected) |
|---|---|---|---|
| BIO1 | 0.860019 | 1.14E-48 | 1.03E-47 |
| BIO2 | 0.6706 | 4.28E-122 | 4.28E-121 |
| BIO3 | 0.8874 | 1.86E-134 | 2.05E-133 |
| BIO4 | 0.755717 | 2.79E-42 | 2.23E-41 |
| BIO7 | 0.687896 | 5.76E-30 | 2.88E-29 |
| BIO12 | 0.585947 | 2.01E-13 | 2.01E-13 |
| BIO15 | 0.71018 | 1.67E-26 | 5.01E-26 |
| elevation | 0.856952 | 1.46E-35 | 1.02E-34 |
| nitrogen | 0.545634 | 1.94E-29 | 7.76E-29 |
| pH | 0.312943 | 3.59E-24 | 7.18E-24 |
| aridity | 0.637648 | 8.45E-31 | 5.07E-30 |

**Table S3.** Detailed results for two-way MANOVA, with variance partitioned by taxonomic group, biogeographic region, and their interaction. Boldface *p*-values were considered significant.

| Predictor | Sum of Squares (normalized) | Mean Squares (normalized) | *p*-value for *F*-test |
|---|---|---|---|
| bio1: taxonomic group | 47.34 | 6.7627 | **6.48E-12** |
| bio1: biogeographic region | 178.33 | 9.9072 | **<2E-16** |
| bio1: group*biogeography | 49.62 | 1.0786 | **0.008478** |
| bio2: taxonomic group | 303.892 | 43.413 | **<2E-16** |
| bio2: biogeographic region | 104.09 | 5.783 | **<2E-16** |
| bio2: group*biogeography | 19.495 | 0.424 | 0.5081 |
| bio3: taxonomic group | 303.892 | 43.413 | **<2E-16** |
| bio3: biogeographic region | 104.09 | 5.783 | **<2E-16** |
| bio3: group*biogeography | 19.495 | 0.424 | 0.5081 |
| bio4: taxonomic group | 36.85 | 5.2648 | **5.83E-11** |
| bio4: biogeographic region | 270.08 | 15.0042 | **<2E-16** |
| bio4: group*biogeography | 35.43 | 0.7701 | 0.06181 |
| bio7: taxonomic group | 17.04 | 2.4339 | **0.000371** |
| bio7: biogeographic region | 56.23 | 14.2353 | **<2E-16** |
| bio7: group*biogeography | 32.13 | 0.6984 | 0.282644 |
| bio12: taxonomic group | 28.34 | 4.0486 | **1.53E-05** |
| bio12: biogeographic region | 94.49 | 5.2495 | **5.76E-15** |
| bio12: group*biogeography | 68.63 | 1.492 | **0.0007799** |
| bio15: taxonomic group | 67.56 | 9.6516 | **6.86E-15** |
| bio15: biogeographic region | 105.68 | 5.8709 | **<2E-16** |
| bio15: group*biogeography | 35.91 | 0.7806 | 0.473 |
| bio17: taxonomic group | 30.37 | 4.3383 | **6.48E-07** |
| bio17: biogeographic region | 139.01 | 7.7227 | **<2E-16** |
| bio17: group*biogeography | 83.03 | 1.8051 | **2.85E-07** |
| elevation: taxonomic group | 104.28 | 14.8975 | **<2E-16** |
| elevation: biogeographic region | 142.32 | 7.9067 | **<2E-16** |
| elevation: group*biogeography | 40 | 0.8697 | 0.078 |
| nitrogen: taxonomic group | 54.27 | 7.7529 | **<2E-16** |
| nitrogen: biogeographic region | 229.23 | 12.7352 | **<2E-16** |
| nitrogen: group*biogeography | 67.37 | 1.4645 | **7.74E-08** |
| carbon: taxonomic group | 52.793 | 7.5418 | **<2E-16** |
| carbon: biogeographic region | 272.794 | 15.1552 | **<2E-16** |
| carbon: group*biogeography | 72.415 | 1.5742 | **1.86E-11** |

| | | | |
|---|---|---|---|
| ph: taxonomic group | 95.15 | 13.5933 | **<2E-16** |
| ph: biogeographic region | 129.81 | 7.2118 | **<2E-16** |
| ph: group*biogeography | 28.76 | 0.6253 | 0.6932 |
| sand: taxonomic group | 176.99 | 25.2847 | **<2E-16** |
| sand: biogeographic region | 61.67 | 3.4259 | **1.26E-10** |
| sand: group*biogeography | 27.35 | 0.5945 | 0.7274 |
| coarsefragment: taxonomic group | 84.76 | 12.1086 | **<2E-16** |
| coarsefragment: biogeographic region | 104.06 | 5.7809 | **<2E-16** |
| coarsefragment: group*biogeography | 27.98 | 0.6083 | 0.8357 |
| needleleaf: taxonomic group | 97.42 | 13.9174 | **<2E-16** |
| needleleaf: biogeographic region | 72.56 | 4.0312 | **1.98E-10** |
| needleleaf: group*biogeography | 12.51 | 0.2721 | 1 |
| deciduousbroadleaf: taxonomic group | 49.93 | 7.1323 | **1.61E-09** |
| deciduousbroadleaf: biogeographic region | 44.13 | 2.4518 | **0.0001021** |
| deciduousbroadleaf: group*biogeography | 55.35 | 1.2033 | **5.43E-02** |
| herbaceous: taxonomic group | 25.7 | 3.6717 | **0.0002146** |
| herbaceous: biogeographic region | 80.49 | 4.4715 | **1.27E-10** |
| herbaceous: group*biogeography | 27.94 | 0.6074 | 0.9508394 |
| aridity: taxonomic group | 44.66 | 6.38 | **7.39E-13** |
| aridity: biogeographic region | 198.75 | 11.0417 | **<2E-16** |
| aridity: group*biogeography | 83.26 | 1.8099 | **4.03E-10** |
